# Expanding the ligandable proteome by paralog hopping with covalent probes

**DOI:** 10.1101/2024.01.18.576274

**Authors:** Yuanjin Zhang, Zhonglin Liu, Marsha Hirschi, Oleg Brodsky, Eric Johnson, Sang Joon Won, Asako Nagata, Matthew D. Petroski, Jaimeen D. Majmudar, Sherry Niessen, Todd VanArsdale, Adam M. Gilbert, Matthew M. Hayward, Al E. Stewart, Andrew R. Nager, Bruno Melillo, Benjamin Cravatt

## Abstract

More than half of the ∼20,000 protein-encoding human genes have at least one paralog. Chemical proteomics has uncovered many electrophile-sensitive cysteines that are exclusive to a subset of paralogous proteins. Here, we explore whether such covalent compound-cysteine interactions can be used to discover ligandable pockets in paralogs that lack the cysteine. Leveraging the covalent ligandability of C109 in the cyclin CCNE2, we mutated the corresponding residue in paralog CCNE1 to cysteine (N112C) and found through activity-based protein profiling (ABPP) that this mutant reacts stereoselectively and site-specifically with tryptoline acrylamides. We then converted the tryptoline acrylamide-N112C-CCNE1 interaction into a NanoBRET-ABPP assay capable of identifying compounds that reversibly inhibit both N112C- and WT-CCNE1:CDK2 complexes. X-ray crystallography revealed a cryptic allosteric pocket at the CCNE1:CDK2 interface adjacent to N112 that binds the reversible inhibitors. Our findings thus provide a roadmap for leveraging electrophile-cysteine interactions to extend the ligandability of the proteome beyond covalent chemistry.

---

Paralogous proteins originate from gene duplication events and retain sequence and structural relatedness, while having evolved to perform overlapping but distinct functions in cells^1–3^. The shared functionality among paralogs is thought to contribute to the robustness of cellular systems, a hypothesis that is supported by the discovery of many synthetic lethality relationships for paralogs in cancer cells^4,5^. Examples include ENO1-ENO2^6^, SMARCA2-SMARCA4^7^, ARID1A-ARID1B^8^ and STAG1-STAG2^9,10^, and recent large-scale CRISPR screens are uncovering several other paralog dependencies^11–13^.

Depending on the unique versus shared functions of paralogous proteins in physiology and disease, chemical probes can be pursued that selectively target one versus several paralogs as a means to optimize drug efficacy and safety. One attractive way to achieve paralog-restricted activity with small molecules is through covalent chemistry targeting specific nucleophilic amino acid residues (e.g., cysteines) found in only one or a subset of related proteins. Covalent ligands targeting paralog-restricted cysteines have been described for kinases (e.g., JAK3_C909^14^, JAK1_C817^15^, FGFR4_C552^16^, PIK3CA_C862^17^) and RNA-binding proteins (NONO_C145)^18^ as well as for somatically mutated forms of KRAS (G12C)^19,20^ and TP53 (Y220C)^21^. While many of these compounds were developed using structure-guided approaches, the covalent chemical probes targeting JAK1_C817 and NONO_C145 were alternatively discovered by chemical proteomics^15,18^, and recent activity-based protein profiling (ABPP) investigations of stereochemically defined sets of electrophilic compounds (‘stereoprobes’) have identified covalent ligands for numerous paralog-restricted cysteines in structurally and functionally diverse proteins^22–24^.

While much attention has been given to targeting paralog-restricted cysteines with covalent chemistry, we wondered whether the discovery of such small molecule-protein interactions might offer even broader insights into the ligandability of the human proteome, if, for instance, paralogs lacking the covalently liganded cysteine might still possess a conserved pocket suitable for reversibly binding small molecules. To address the question of pocket conservation across paralogous proteins, we describe herein a strategy where electrophile-sensitive cysteines discovered by ABPP are introduced into a cysteine-less paralog by site-directed mutagenesis to enable screening with stereoprobe libraries, where stereoselective and site-specific covalent binding to the cysteine mutant protein is interpreted as evidence of a ligandable pocket. We then convert the stereoprobe ligands into high throughput screening-compatible probes to identify reversibly binding compounds capable of also targeting the wild-type paralog. We apply this approach to the cyclin paralogs CCNE2 and CCNE1, which are key regulators of cyclin-dependent kinase activity in cancer^25^ and possess or lack a ligandable cysteine, respectively (C109 in CCNE2^24^ and N112 in CCNE1). We identify tryptoline acrylamide stereoprobe ligands for the N112C-CCNE1 mutant and show that these probes display competitive binding with reversible 2,6-diazaspiro[3.4]octane inhibitors of CCNE1/CDK2 complexes recently reported in the patent literature^26^. Structural studies uncover a cryptic allosteric binding pocket for the reversible inhibitors residing at the interface of CCNE1 and CDK2 in proximity to N112 of CCNE1. Our findings thus describe a “paralog-hopping” strategy to illuminate reversible small molecule-binding pockets in the proteome from electrophile-cysteine interaction maps.

## Results

### Stereoprobe reactivity with a N112C-CCNE1 mutant

A recent in-depth ABPP study of stereoprobe reactivity with proteins in human cancer cell lines identified tryptoline acrylamides that stereoselectively and site-specifically react with C109 of CCNE2 (**Fig. 1a-c**)^24^. The stereoselective engagement of CCNE2 by (*R*, *R*)-tryptoline acrylamide WX-03-59 and its alkyne analog WX-03-348 was observed by protein- (**Fig. 1b**) and cysteine- (**Fig. 1c**) directed ABPP, and we verified this reactivity profile with purified recombinant CCNE2:CDK2 complexes by gel-ABPP, where we found that WX-03-348 stereoselectively reacted with WT-CCNE2, but not a C109A-CCNE2 mutant (**Fig. 1d**).

**Figure 1.**
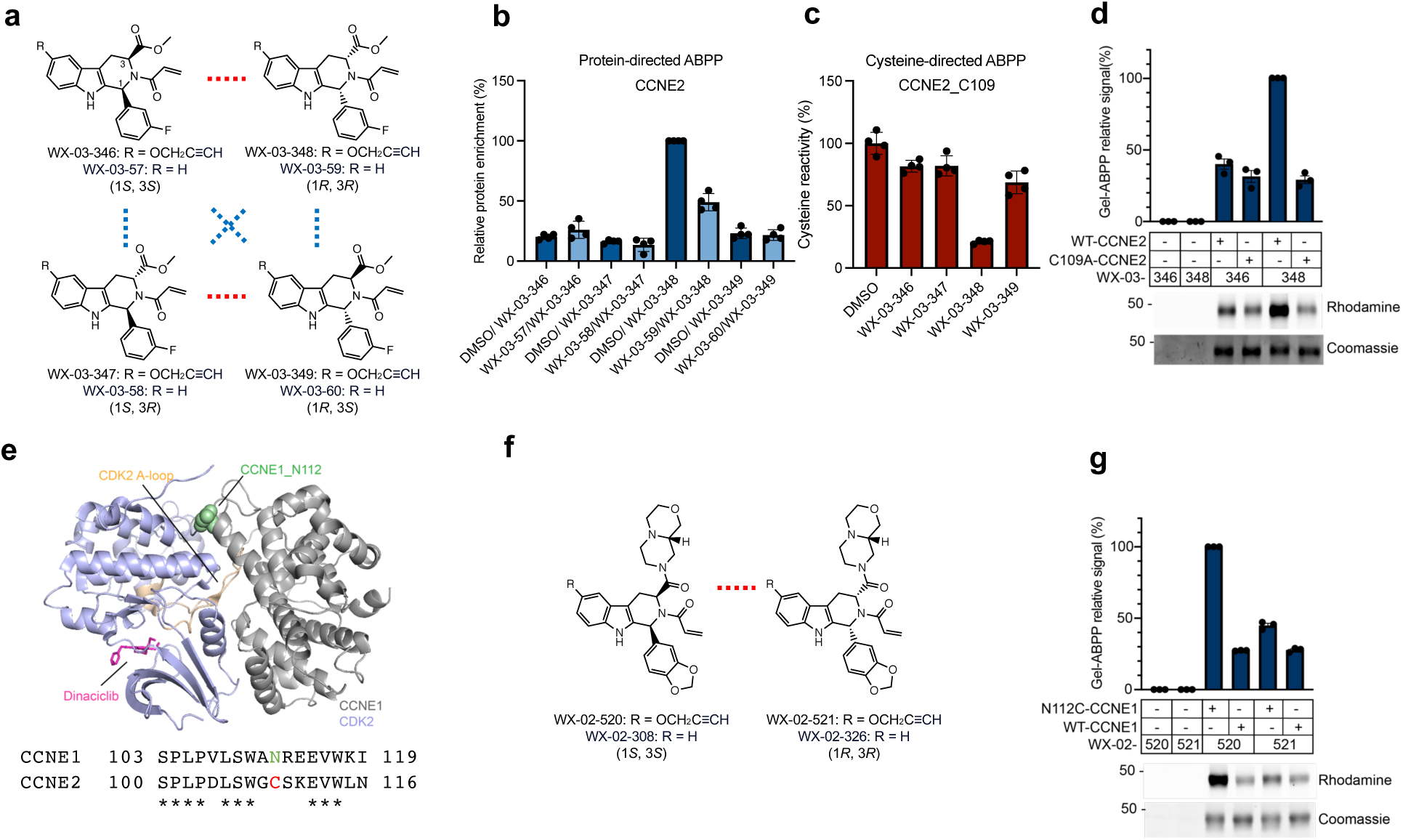
Covalent stereoprobes targeting CCNE2 and an N112C-CCNE1 mutant. **a**, Structure of tryptoline acrylamide stereoprobes WX-03-348 and WX-03-59 that stereoselectively engage CCNE2_C109 alongside inactive stereoisomers. Red lines connect enantiomers and blue lines connect diastereomers. **b**, Protein-directed ABPP data showing stereoselective enrichment of CCNE2 by WX-03-348 (5 µM, 1 h) and blockade of this enrichment by pre-treatment with WX-03-59 (20 µM, 2 h) in 22Rv1 cells. Data are from ref. ^24^ and represent average values ± S.D., n = 4. **c**, Cysteine-directed ABPP data showing stereoselective engagement of CCNE2_C109 by WX-03-348 (20 µM, 3 h) in 22Rv1 cells. Data are from ref. ^24^ and represent average values ± S.D., n = 4. **d**, Gel-ABPP data showing stereoselective engagement of recombinant WT-CCNE2, but not a C109-CCNE2 mutant, by WX-03-348 (bottom, representative gel-ABPP data; top, quantification of data). Purified CCNE1:CDK2 complexes (1 µM) were treated with WX-03-348 or enantiomer WX-03-346 (5 µM, 1 h) followed by conjugation to a rhodamine-azide reporter tag (Rh-N_3_) by copper-catalyzed azide-alkyne cycloaddition (CuAAC^72,73^) chemistry, SDS-PAGE, and in-gel fluorescence scanning. Coomassie blue signals correspond to WT- or C109A-CCNE2. Data represent average values ± s.e.m., n = 3. **e**, Top, crystal structure of a WT-CCNE1:CDK2 (gray: purple) complex bound to the orthosteric inhibitor dinaciclib (magenta)^74^ (PDB 5L2W), showing location of N112 of CCNE1 (green sphere) in proximity to the CDK2 binding interface. Also highlighted is the CDK2 A-loop (activation loop) (orange). Bottom, local sequence alignment for CCNE1 and CCNE2 around CCNE1_N112/CCNE2_C109 with conserved residues marked by asterisks. **f**, Structures of an alkyne and non-alkyne pair of stereoprobes – WX-02-520 and WX-02-308, respectively – that stereoselectively engage the N112C-CCNE1 mutant, alongside the corresponding inactive enantiomers WX-02-521 and WX-02-326. **g**, Gel-ABPP data showing stereoselective engagement of recombinant N112C-CCNE1, but not WT-CCNE1 by WX-02-520 (5 µM, 1 h) (bottom, representative gel-ABPP data; top, quantification of data). Purified CCNE1:CDK2 complexes were assayed as described in **d**. Data represent average values ± s.e.m., n = 3.

CCNE2 is one of two E-type cyclins in human cells that bind to and activate CDK2 to promote cell cycle progression^25^. The heightened expression of CCNE2 and its paralog CCNE1 supports the growth of diverse cancers and plays a role in drug resistance^27–30^. While both CCNE1 and CCNE2 are assigned as *strongly selective* cancer dependencies in the Cancer Dependency Map^31^, many more human cancer cell lines appear to require CCNE1 for growth (**Extended Data Fig. 1**), likely reflecting its frequent amplification in tumor types like ovarian cancer^32^. CCNE1 and CCNE2 share ∼55% sequence identity, but CCNE2_C109 is not conserved and corresponds to N112 in CCNE1 (**Fig. 1e**). Crystal structures localize N112 to a CCNE1:CDK2 interface that also harbors the CDK2 A-loop (activation loop) (**Fig. 1e**), suggesting the potential for ligands binding at this site to allosterically regulate CDK2 function.

Acknowledging the broader apparent cancer relevance of CCNE1 and its generally high sequence relatedness to CCNE2, we wondered if CCNE1 might also possess the stereoprobe-binding pocket discovered in CCNE2. We set out to test this hypothesis by evaluating a focused library of alkyne-modified tryptoline acrylamides for reactivity with purified WT-CCNE1:CDK2 and N112C-CCNE1:CDK2 complexes by gel-ABPP (**Extended Data** Figs. 2 and 3). Multiple tryptoline acrylamides showed preferential reactivity with N112C-CCNE1 compared to WT-CCNE1, with WX-02-520 displaying the strongest stereoselectivity (**Fig. 1f, g** and **Extended Data** Figs. 2 and 3). We also confirmed that pretreatment with the non-alkyne analog WX-02-308 (**Fig. 1f**) stereoselectively blocked WX-02-520 reactivity with N112C-CCNE1 (**Extended Data Fig. 4a**). Interestingly, WX-02-520 and WX-02-308 are (*S*, *S*) tryptoline acrylamides and did not engage CCNE2_C109 (**Extended Data Fig. 4b**), while WX-03-348 – the (*R*, *R*) tryptoline acrylamide ligand targeting CCNE2_C109 – did not stereoselectively engage CCNE1_N112C (**Extended Data Fig. 2**). We confirmed these respective stereoprobe reactivity preferences by mass spectrometry with purified N112C-CCNE1:CDK2 and CCNE2:CDK2 complexes (**Extended Data Fig. 5**). Our results thus indicate the presence of a small molecule binding pocket proximal to N112 and C109 in CCNE1 and CCNE2, respectively, that shows a distinct structure-activity relationship for stereoprobe engagement in each protein.

### Stereoprobe reactivity with N112C-CCNE1 measured by NanoBRET-ABPP

We next set out to establish a higher-throughput assay to screen for small molecules that can block tryptoline acrylamide stereoprobe reactivity with N112C-CCNE1. We first synthesized derivatives of WX-02-308 that possess PEG3-methyl ester linkers at different locations on the tryptoline core (**Extended Data Fig. 6a**) and tested these compounds for blockade of WX-02-520 reactivity with N112C-CCNE1 by gel-ABPP. We observed superior stereoselective reactivity for the compound bearing a PEG3-methyl ester extended from the amide group on C3 of the tryptoline core (WX-02-588 conjugate) compared to the compound with a PEG3-methyl ester appended to C6 of the tryptoline core (WX-02-520 conjugate) (**Extended Data Fig. 6b**). We accordingly coupled a NanoBRET^TM^ 590 dye to WX-02-588 and its enantiomer WX-02-589 to generate fluorescent analogs YZ-01-A and YZ-01-B, respectively (**Fig. 2a**).

**Figure 2.**
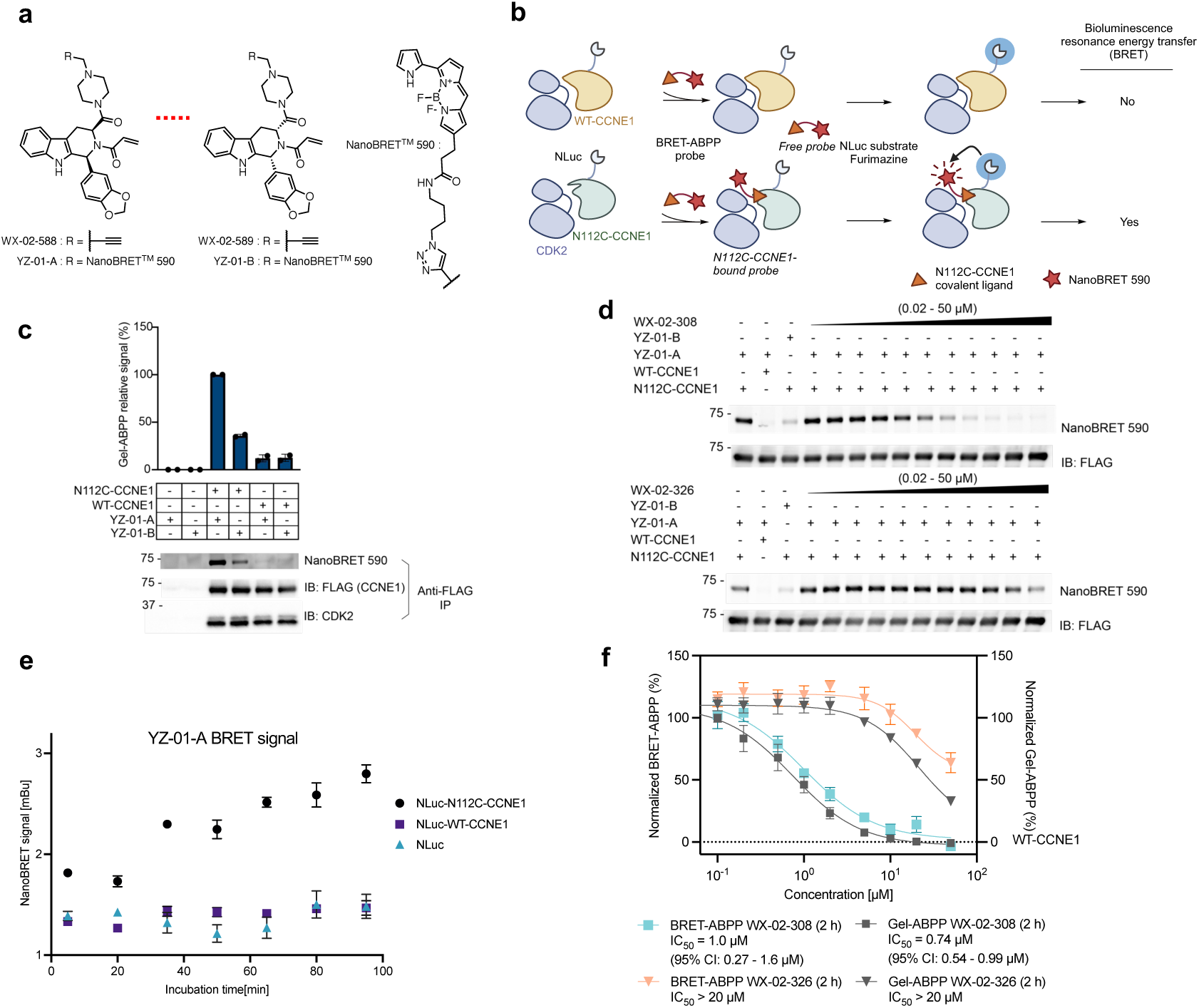
NanoBRET-ABPP assay for measuring stereoprobe interactions with N112C-CCNE1. **a**, Left, structures of NanoBRET stereoprobe YZ-01-A and the corresponding alkyne analog WX-02-588, alongside the corresponding enantiomers YZ-01-B and WX-02-589. Right, structure of NanoBRET^TM^590 fluorophore and linker conjugated to WX-02-588/WX-02-589 by CuAAC. **b**, Schematic for NanoBRET-ABPP assay to measure stereoprobe interactions with N112C-CCNE1. Created with BioRender.com. **c**, Gel-ABPP data showing stereoselective engagement of recombinant NLuc-N112C-CCNE1-3xFLAG, but not NLucWT-CCNE1-3xFLAG by YZ-01-A (bottom, representative gel-ABPP data; top, quantification of data). NLuc-CCNE1 proteins were co-expressed with CDK2 in HEK293T cells by transient transfection, and cell lysates treated with YZ-01-A or YZ-01-B (10 µM, 90 min), after which CCNE1 proteins were immunoprecipitated with an anti-FLAG antibody and processed for gel-ABPP. Data represent average values ± s.e.m., n = 2. **d**, Gel-ABPP data showing concentration-dependent, stereoselective blockade of YZ-01-A engagement of NLuc-N112C-CCNE1 by WX-02-308 compared to WX-02-326. Lysates from HEK293T cells co-transfected with NLuc-N112C-CCNE1 or NLuc-WT-CCNE1 and CDK2 were treated with WX-02-308 or WX-02-326 (0.02 – 50 µM, 2 h) followed by YZ-01-A (10 µM, 90 min), anti-FLAG immunoprecipitation, and processing for gel-ABPP. See panel **f** for quantification of gel-ABPP data. **e,** NanoBRET data showing time-dependent increase in YZ-01-A (10 µM, 2 h) reactivity with NLuc-N112C-CCNE1 vs NLuc-WT-CCNE1 or NLuc control proteins. NanoBRET-ABPP assays were performed in transfected HEK293T cell lysates. **f**, Comparison of gel- and NanoBRET-ABPP data showing concentration-dependent, stereoselective blockade of YZ-01-A engagement of NLuc-N112C-CCNE1 by WX-02-308 compared to WX-02-326 (0.2 – 50 µM, 2 h). Dashed horizontal line marks background signals for NanoBRET-ABPP assays performed with YZ-01-A (10 µM, 90 min) and NLuc-WT-CCNE1. CI, confident intervals. Data represent average values ± s.e.m., n = 3.

We first verified by gel-ABPP that YZ-01-A stereoselectively reacted with a purified N112C-CCNE1:CDK2 complex to a much greater extent than a purified WT-CCNE1:CDK2 complex (**Extended Data Fig. 7a**). We next established a NanoBRET-ABPP assay^33^ for measuring the reactivity of YZ-01-A with N112C-CCNE1. NanoBRET measures the energy transfer from a donor nanoluciferase (NLuc)-fusion protein-of-interest (in our case, N112C-CCNE1) to an acceptor fluorophore conjugated to an interacting biomolecule (in our case, YZ-01-A) (**Fig. 2b**). NanoBRET systems have the advantages of serving as homogeneous assays for identifying small molecules that block biomolecular interactions and compatibility with assaying these interactions in native biological systems (e.g., cell lysates or even *in cellulo*). We recombinantly co-expressed NLuc-N112C-CCNE1 or NLuc-WT-CCNE1 with CDK2 by transient transfection in HEK293T cells and then incubated whole-cell lysates with YZ-01-A or YZ-01-B (10 µM, 90 min). Analysis of the samples by gel-ABPP revealed stereoselective engagement of NLuc-N112C-CCNE1, but not NLuc-WT-CCNE1, by YZ-01-A (**Fig. 2c** and **Extended Data Fig. 7b**), and we further confirmed that this interaction was stereoselectively blocked in a concentration-dependent manner by pre-treatment with WX-02-308 (**Fig. 2d**).

We next evaluated the reactivity of YZ-01-A with NLuc-N112C-CCNE1 by NanoBRET-ABPP, which revealed a time-dependent increase in BRET signal that was much greater than the signals observed with NLuc-WT-CCNE1-transfected or NLuc-transfected cell lysates (**Fig. 2e**). The specific reactivity of YZ-01-A continued to increase across the 100 min assay window, indicating incomplete engagement of NLuc-N112C-CCNE1 that should be compatible with measuring the impact of reversibly or irreversibly binding small-molecule competitors, as has been shown with other formats for ABPP^34,35^. We selected an assay time point of 90 min for subsequent NanoBRET-ABPP assays of YZ-01-A reactivity with NLuc-N112C-CCNE1, as it offered a good signal over background ratio in lysates expressing NLuc-WT-CCNE1 (**Fig. 2e**). We finally used the NanoBRET-ABPP assay to measure stereoselective engagement of NLuc-N112C-CCNE1 by WX-02-308, as well as a related tryptoline acrylamide WX-02-14, which furnished IC50 values matching those generated by gel-ABPP (**Fig. 2f** and **Extended Data Fig. 7c-e**). As expected for irreversible ligands, both WX-02-308 and WX-02-14 showed greater potency in 2 h pretreatment vs no pretreatment conditions in NanoBRET assays (**Extended Data Fig. 7f**).

### NanoBRET-ABPP identifies allosteric inhibitors of CCNE1:CDK2 complexes

The high cancer relevance of CCNE1 has motivated efforts to discover small molecules that can inhibit CCNE1:CDK2 complexes. While orthosteric (ATP site-competitive) CDK2 inhibitors have been described^36,37,38^, these compounds face selectivity challenges across the CDK family and broader kinome and also do not discriminate among the various cyclin (A and E-type):CDK2 complexes. We reasoned that the NanoBRET-ABPP assay could identify alternative types of reversibly binding compounds to N112C-CCNE1:CDK2 complexes that would also have the potential to bind WT-CCNE1:CDK2 complexes. In this regard, a class of 2,6- diazaspiro[3.4]octane inhibitors of CCNE1:CDK2 complexes was recently reported in the patent literature^26^. We synthesized two representative compounds from the patent – I-125A and I-198 (**Fig. 3a**) – and first confirmed that they inhibited the activity of purified WT-CCNE1:CDK2 and N112C-CCNE1:CDK2 complexes using a C-terminal peptide of retinoblastoma protein 1 (RB1) as a substrate (**Fig. 3b**, left; and **Extended Data Fig. 8a**). I-125A was more active than I-198 as a CCNE1:CDK2 inhibitor (IC50 values of 0.13 and 0.75 µM, respectively), and we observed similar relative potencies for the compounds when tested with a second CDK2 substrate – histone H1 protein – although, in this case, neither compound fully inhibited WT-CCNE1:CDK2 or N112C-CCNE1:CDK2 complexes (**Fig. 3b**, right; and **Extended Data Fig. 8a**). The orthosteric CDK2 inhibitor dinaciclib fully blocked the activity of WT-CCNE1:CDK2 and N112C-CCNE1:CDK2 complexes with either RB1 peptide or H1 protein substrates (**Fig. 3b** and **Extended Data Fig. 8a**). In contrast, the tryptoline acrylamides WX-02-308 and WX-02-14 did not inhibit N112C-CCNE1:CDK2 complexes in either substrate assay (**Fig. 3b**) at concentrations that fully engaged N112C in ABPP experiments (**Fig. 2f** and **Extended Data Fig. 7c-e**). Finally, neither I-125A or I-198 affected the activity of CCNE2:CDK2 complexes in RB1 peptide or H1 protein assays (**Extended Data Fig. 8b**), suggesting that these compounds act as paralog-restricted inhibitors of CCNE1:CDK2 complexes.

**Figure 3.**
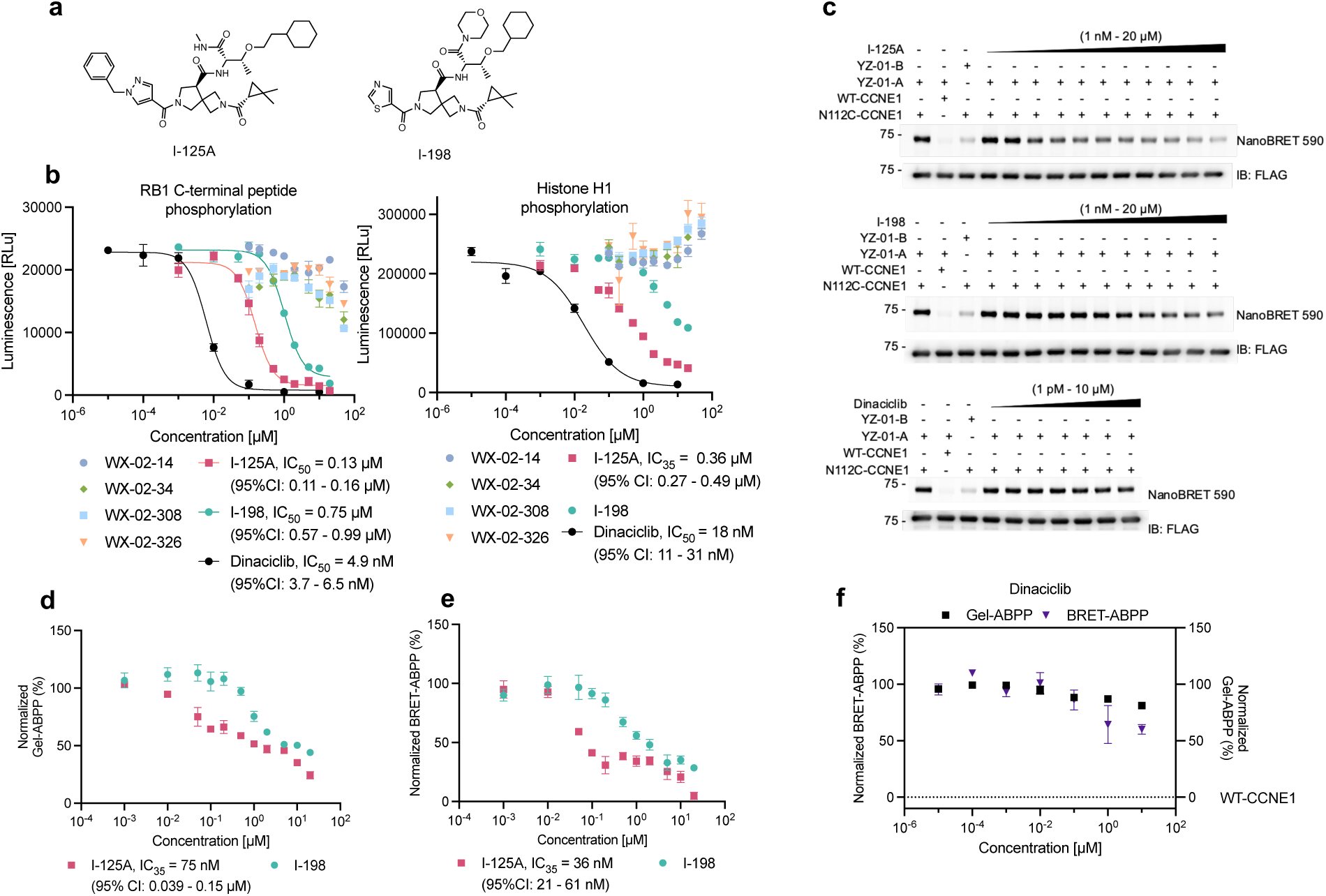
Characterizing allosteric inhibitors of the CCNE1:CDK2 complex. **a**, Structures of two inhibitors of the CCNE1:CDK2 complex – I-125A and I-198 – reported in a patent^26^. **b**, ADP- Glo data showing effects of the indicated compounds on the activity of a purified N112C-CCNE1:CDK2 complex as measured by phosphorylation of the C-terminal peptide of RB1 (left) or the histone H1 protein (right). The N112C-CCNE1:CDK2 complex was pre-treated with compounds for 2 h (37 °C) before initiating phosphorylation reactions by addition of substrate. CI, confident intervals. Data represent average values ± s.e.m., n = 3. **c**, Gel-ABPP data showing concentration-dependent partial blockade of YZ-01-A engagement of N112-CCNE1 by I-125A and I-198, but not dinaciclib. Assays were performed in lysates from transfected HEK293T cells co-expressing NLuc-N112C-CCNE1 or NLuc-WT-CCNE1 and CDK2 that were treated with the indicated concentration ranges of compounds followed by YZ-01-A (10 µM, 90 min), anti-FLAG immunoprecipitation, and processing for gel-ABPP. **d**, Quantification of gel-ABPP data shown in **c**. Data represent average values ± s.e.m., n = 3. **e**, NanoBRET-ABPP data showing concentration-dependent partial blockade of YZ-01-A engagement of N112-CCNE1 by I-125A and I-198. Samples were treated with compounds as described in **c** prior to NanoBRET measurement. Data represent average values ± s.e.m., n = 3. **f**, Quantification of gel- and NanoBRET-ABPP data showing that dinaciclib does not substantially affect YZ-01-A engagement of N112C-CCNE1. Data represent average values ± s.e.m., n = 3.

We next tested the effects of I-125A and I-198 on YZ-01-A reactivity with N112C-CCNE1. Using either gel- (**Fig. 3c, d**) or NanoBRET-ABPP (**Fig. 3e**), we observed a clear concentration-dependent blockade of YZ-01-A engagement of N112C-CCNE1 by both I-125A and I-198. Consistent with the substrate assays, I-125A showed greater potency than I-198 in blocking the YZ-01-A-N112C-CCNE1 interaction. Interestingly, the inhibitory effects of I-125A and I-198 were incomplete in both the gel- and NanoBRET-ABPP assays, resembling the partial suppression of CCNE1:CDK2 activity observed for these compounds in the histone H1 substrate assay. We confirmed that the residual ∼30-40% I-125A/I-198-insensitive reactivity of N112C-CCNE1 with YZ-01-A persisted in gel-ABPP assays performed at different time points (**Extended Data Fig. 8c**) and maintained site-specificity (i.e., was not observed with WT-CCNE1) and stereoselectivity (i.e., was not observed with YZ-01-B) (**Extended Data Fig. 8d**). For the more active I-125A ligand, we were able to estimate “IC35 values” of ∼35-75 nM by gel- and NanoBRET-ABPP (**Fig, 3d, e**) that were similar, albeit somewhat more potent than the IC35 values measured with purified CCNE1:CDK2 complexes and a histone H1 substrate (IC35 values of ∼130-360 nM; **Fig. 3b**). While we do not yet understand the mechanistic basis for partial inhibition of CCNE1:CDK2 complexes by I-125A and I-198 in certain functional assays, it is possible that these complexes are a composite of different CCNE1:CDK2 proteoforms (e.g., a mixed population of phosphorylation states) that show distinct sensitivities to the 2,6-diazaspiro[3.4]octane inhibitors. Finally, we found that I-125A and I-198 did not affect the binding of the orthosteric (ATP site) NanoBRET tracer K-10 to WT-CCNE1:CDK2 or N112C-CCNE1:CDK2 complexes, which contrasted with the full inhibitory activity observed for dinaciclib in this assay (**Extended Data Fig. 8e**). And, on the other hand, dinaciclib did not alter YZ-01-A interactions with N112C-CCNE1 in gel- or NanoBRET-ABPP assays (**Fig. 3f**).

Taken together, these data demonstrate that ABPP assays measuring covalent tryptoline acrylamide stereoprobe-N112C-CCNE1 interactions can detect the binding and activity of a recently described class of structurally distinct and reversible 2,6-diazaspiro[3.4]octane inhibitors of CCNE1:CDK2 complexes, even though the stereoprobes themselves do not inhibit this complex. Considering further that the ABPP assays were insensitive to an orthosteric CDK2 inhibitor dinaciclib, and, conversely, the binding of the orthosteric K-10 tracer to CDK2 was unaffected by the 2,6-diazaspiro[3.4]octane inhibitors, we conclude that CCNE1:CDK2 complexes possesses at least two distinct small molecule-binding pockets (one orthosteric and one or more allosteric). We next set out to gain structural insights into the modes of allosteric compound binding to CCNE1:CDK2 complexes.

### Crystal structures of allosteric inhibitor-CCNE1:CDK2 complexes

Co-crystal structures of a WT-CCNE1 (residues 96-378):CDK2 complex with I-125A and I-198 were solved at 1.9 and 2.0 Å resolution, respectively (**Supplementary Table 1**), and found to largely resemble previously reported structures of CCNE1-CDK2 complexes in the apo form^39^ or in complex with orthosteric inhibitors such as PF-06873600^13^. A clear difference, however, was electron density corresponding to a single molecule of I-125A or I-198 bound at a cryptic pocket at the CCNE1-CDK2 interface (**Fig. 4a, b** and **Extended Data Fig. 9a-c**). The binding pocket is adjacent to the kinase C-lobe and A-loop, but distal to the ATP binding site and adjacent to α-helices α1, α7, and α8 of CCNE1 (**Fig. 4a, b** and **Extended Data Fig. 9a-c**). N112 of CCNE1 is located at the origin of α1 within 10 Å of the I-125A and I-198 molecules (**Fig. 4a, b** and **Extended Data Fig. 9a-c**), providing a rationale for competitive binding of these compounds with covalent stereoprobes targeting the N112C-CCNE1 mutant. Key protein contacts with the 2,6-diazaspiro[3.4]octane inhibitors include hydrogen-bonding interactions between the side-chain guanidine of R122 of CDK2 and the (*gem*-dimethyl)cyclopropyl amide carbonyl of I-125A and I-198 and between the backbone amide of T182 of CDK2 and the thiazole or pyrazole nitrogen of I-125A and I-198, respectively, as well as numerous hydrophobic and solvent mediated interactions (**Fig. 4c** and **Extended Data Fig. 9d**).

**Figure 4.**
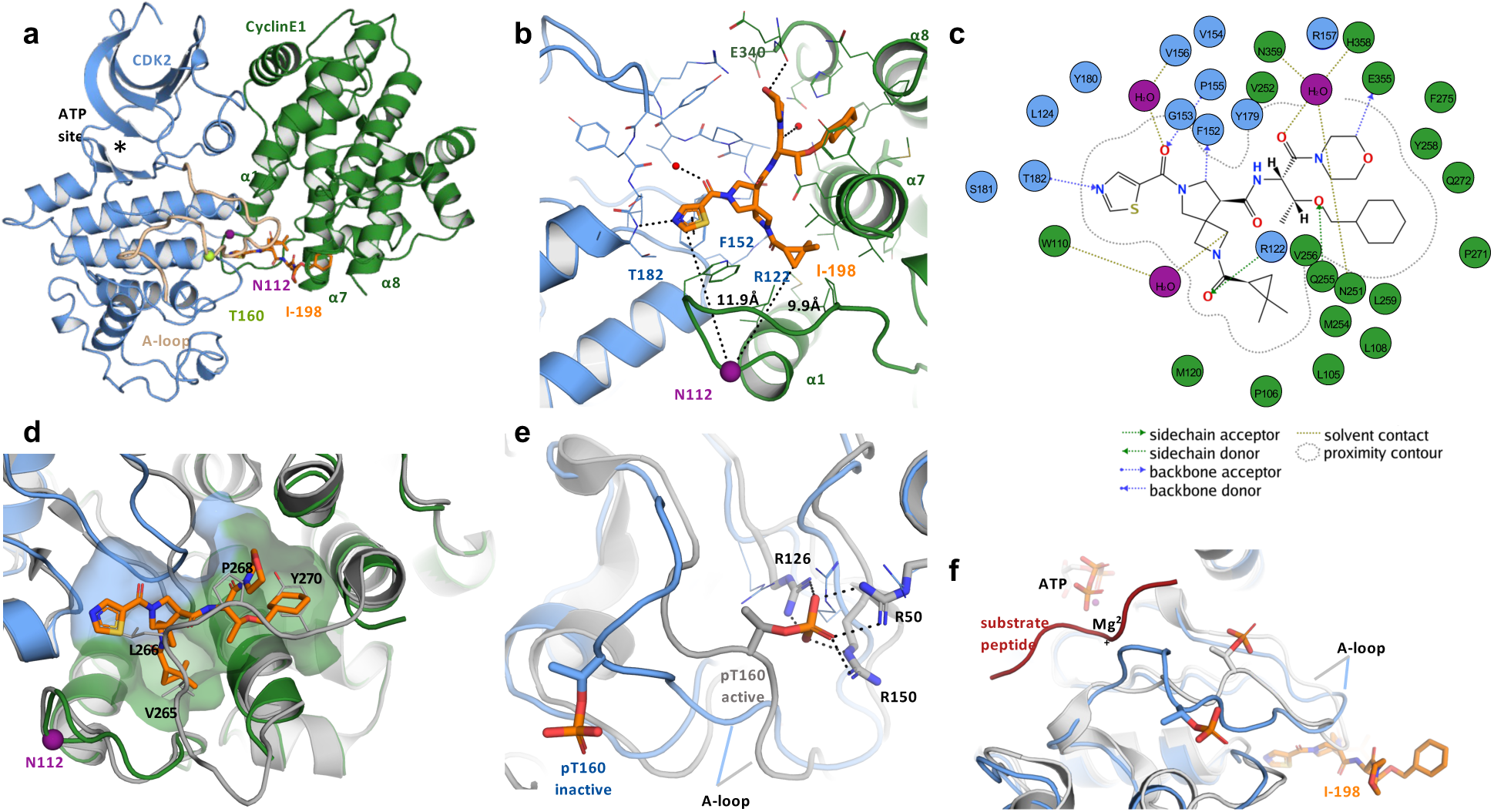
Crystal structure of an I-198-CCNE1:CDK2 complex. **a**, Ribbon diagram structure of CCNE1:CDK2 (green: blue) complex with the I-198 compound shown in orange, the ATP-binding pocket in black (asterisk), the CDK2 activation loop (A-loop) in beige (also containing the phosphorylated T160 (pT160) residue in lime, and N112 in purple. **b**, Detailed view of the I-198 binding site, protein-ligand interactions, and the distance to N112 indicated by dashes. **c**, Protein-ligand interaction plot between CCNE1:CDK2 and I-198 (generated in Molecular Operating Environment software). CCNE1 and CDK2 residues are shown in green and blue, respectively, and water molecules are in magenta. **d**, I-198 binds in a cryptic pocket that is not present in the CCNE1:CDK2 apo structure (grey). The CDK2 A-loop and CCNE1 loop containing amino acids L244-P266 are reorganized to create a pocket in the I-198-bound structure. **e**, Overlay of apo (gray; W981QMZ) and I-198-bound (blue) CCNE1:CDK2 structures showing the position of pT160. In the apo structure, pT160 is coordinated by three arginines (R50, R126 and R150), which is a hallmark of a fully activated cyclin:CDK2 complex. I-198 induces an A-loop conformation that moves pT160 away from the arginines, creating a structure that is consistent with an inactivated kinase. **f**, Overlay of activated CCNA2:CDK2 with bound ATP and substrate peptide (gray and red; 1QMZ) and I-198-bound (blue) CCNE1:CDK2 structures showing the conformational change in the A-loop induced by I-198 binding that results in clashes with substrate peptide binding.

An overlay of the I-198 co-crystal structure with an apo structure of CCNE1:CDK2^39^ helped to explain the origins of the cryptic allosteric pocket, as substantial structural rearrangements are required to accommodate ligand binding (**Fig. 4d**). Indeed, the I-198 binding pocket, which has a volume of ∼80 Å^3^, is mostly absent in the apo CCNE1:CDK2 structure (**Fig. 4d**) and is formed by the movement of two loops – the loop connecting helices α7 and α8 of CCNE1 (residues L244-P266; **Fig. 4d**) and the A-loop of CDK2 (**Fig. 4e**). Notably, the cyclohexyl group of I-125A/I-198 swaps positions with the sidechain of Y270 of CCNE1 in the apo structure, which appears to contribute to the remodeling of the α7-α8 loop (**Fig. 4d**). The A-loop movement disrupts the canonical coordination of the oxygen atoms of phosphorylated T160 of CDK2 with sidechain nitrogen atoms from R50, R126 (of the HRD motif), and R150 (**Fig. 4e**) that is required for CDK2 to adopt a fully active state^40^. The rearrangement of the A-loop further appears to be incompatible with substrate peptide binding, as superposition of the I-198 co-crystal structure with that of activated CCNA2:CDK2 with bound ATP and substrate peptide^41^ revealed clashes between residues of the A-loop and the substrate peptide (**Fig. 4f**).

Although we cannot rule out that different A-loop conformation(s) or alternate substrate sequences may be compatible with simultaneous binding of I-125A/I-198 and substrate, these structural data provide mechanistic insights that help to explain how allosteric small molecules binding at the CCNE1:CDK2 interface can remodel this complex to inhibit CDK2 activity. Finally, we noted that most of the residues contacting I-125A/I-198 are conserved between CCNE1 and CCNE2 (**Extended Data Fig. 9e**), suggesting that other structural differences, such as residues interfacing with CDK2, may contribute to the CCNE1-restricted activity displayed by these allosteric inhibitors.

### Distinct functional effects of allosteric ligands binding the CCNE1:CDK2 complex

The discovery that N112 of CCNE1 is located proximal to the allosteric binding pocket for the 2,6-diazaspiro[3.4]octane inhibitors suggested that diverse types of chemistry are capable of binding to this site in the CCNE1:CDK2 complex. Nonetheless, tryptoline acrylamides like WX-02-308 and WX-02-14, despite engaging N112C-CCNE1 with high stereoselectivity and stoichiometry, did not inhibit CDK2 catalytic activity, indicating that different ligands binding the allosteric site may have variable functional effects on CCNE1:CDK2 complexes. Such divergence between small-molecule binding and activity has been observed for allosteric sites in other proteins^42,43^, and we wondered whether the tryptoline acrylamides acted as silent ligands or might instead cause distinct functional changes in CCNE1:CDK2 complexes. To begin to explore this concept, we performed immunoprecipitation-mass spectrometry (IP-MS) experiments on FLAG epitope-tagged WT-CCNE1:CDK2 and N112C-CCNE1:CDK2 complexes from HEK293T cells treated with tryptoline acrylamides or I-125A. Among the co-immunoprecipitated proteins, two cyclin-dependent kinase regulatory subunits CKS1B and CKS2 showed substantially greater enrichment with FLAG-N112C-CCNE1:CDK2 complexes from cells treated with WX-02-308 or WX-02-14 (**Fig. 5a, b** and **Extended Data Fig. 10a** and **Supplementary Dataset 1**). This effect was both stereoselective (not observed with inactive enantiomers WX-02-326 and WX-02-34) and site-specific (not observed with WT-CCNE1:CDK2 complexes) and also not caused by I-125A, which instead impaired the co-enrichment of a single protein PRC1 (**Fig. 5a, b**).

**Figure 5.**
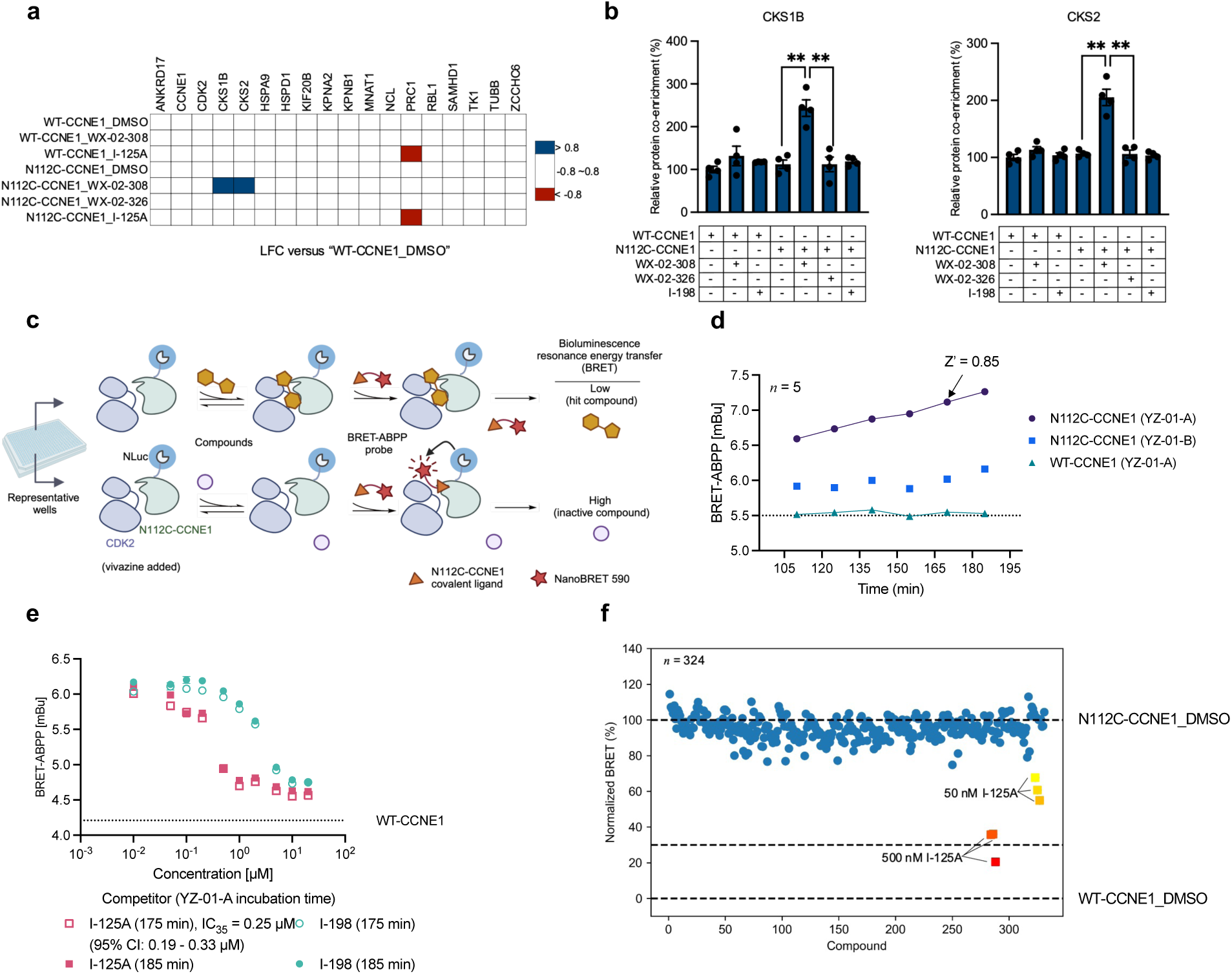
Functional analysis and high-throughput screening of the allosteric pocket in CCNE1:CDK2 complexes. **a**, Heat map showing effects of the indicated compounds on proteins that co-immunoprecipitate with CCNE1-3xFLAG. HEK293T cells recombinantly expressing WT- or N112C-CCNE1-3xFLAG and CDK2 were treated with WX-02-308 or WX-02-326 (20 µM), I-125A (1 µM), or DMSO for 6 h, followed by cell lysis, anti-FLAG enrichment, and MS-based proteomic analysis. Shown are proteins with ≥ 3 spectral counts that were enriched ≥ 4-fold in DMSO-treated WT-CCNE1-3xFLAG cells compared to DMSO-treated mock-transfected cells. Data are normalized to CCNE1 signals within each experiment. LFC, log2 (fold change). **b**, Quantification of data for CKS1B and CKS2 in co-immunoprecipitation experiments performed in **a**. Data represent average values ± s.e.m., n = 4. Unpaired t test with Welch’s correction was used to test statistical significance. p values: 0.0026 and 0.0023 for CKS1B in WX-02-308-treated vs DMSO-treated and WX-02-326-treated N112C-CCNE1-expressing cells, respectively; 0.0054 and 0.0028 for CKS2 in WX-02-308-treated vs DMSO-treated and WX-02-326-treated N112C-CCNE1-expressing cells, respectively. **c**, Schematic for high-throughput screening-compatible NanoBRET-ABPP assay with an N112C-CCNE1:CDK2 complex. HEK293T cell lysate recombinantly expressing NLuc-N112C-CCNE1 and CDK2 (20 µL of cell lysates at 2 mg protein/µL) are mixed with vivazine (used at 1:100 as recommended by Promega) and YZ-01-A (10 µM), then dispensed into a 384-well plate that is pre-plated with individual compounds per well (0.05 µL of 4 mM compound stocks in DMSO; final compound concentration of 10 µM). Compound effects on the reaction between YZ-01-A with NLuc-N112C-CCNE1:CDK2 are measured by BRET with signals from a reaction with YZ-01-A and NLuc-WT-CCNE1:CDK2 serving as a background control. Created with BioRender.com. **d**, Time-dependent assessment of signal intensity for the NanoBRET assay performed as described in **c**. Arrow marks the 170-min time point chosen for HTS because, at this time point, the reaction between YZ-01-A and NLuc-N112C-CCNE1:CDK2 is still incomplete and a good Z’ score is achieved (0.85). Data are average values ± s.e.m., n = 5. **e**, NanoBRET-ABPP data acquired with HTS assay conditions at 175 or 185 min time points showing concentration-dependent partial blockade of YZ-01-A engagement of N112C-CCNE1 by I-125A and I-198. Data are average values ± s.e.m., n = 4. **f**, A pilot NanoBRET-ABPP screen performed as described in **c** of 324 compounds from the ChemBridge macrocycles collection (10 µM final concentration) alongside the indicated concentrations of I-125A. Dashed lines mark maximum signal (top, DMSO control with NLuc-N112C-CCNE1:CDK2 complex), 70% inhibition of signal (middle), and background signal (bottom, DMSO control with NLuc-WT-CCNE1:CDK2 complex).

CKS1B and CKS2 are small (∼80 amino acid) paralogous adaptor proteins that regulate the activity of cyclin-CDK complexes^44^. The CKS proteins have been implicated in promoting multisite phosphorylation of substrates by CDKs, which is important for integrating diverse signaling inputs in cellular processes like mitosis^45^. Most recently, CKS1B has been shown to diversify the substrate scope of CDK1 by promoting multisite phosphorylation events that include noncanonical (non-proline-directed) sites^46^. An overlay of the crystal structures of CKS1B:CDK2^47^ and CCNE1:CDK2 complexes showed that CKS1B and CCNE1 bind to opposing and non-overlapping interfaces of CDK2, which places CKS1B at a distal location from the presumed binding pocket for tryptoline acrylamides (**Extended Data Fig. 10b**). We therefore speculate that tryptoline acrylamide-induced stabilization of the CKS1B/CKS2:CDK2:N112C-CCNE1 complex occurs in an allosteric manner. Based on current understanding of CKS1B function, such stabilization might in turn influence the global substrate scope of CDK2-CCNE1 complexes.

Intrigued by the finding that 2,6-diazaspiro[3.4]octanes and tryptoline acrylamides confer different functional effects on CCNE1:CDK2 complexes (inhibition of CDK2 activity vs remodeling of complex composition, respectively), we set out to develop a high-throughput screening (HTS) assay that can be used to identify additional structural classes of compounds that bind to the allosteric site in this complex. To achieve this goal, we made multiple modifications to NanoBRET-ABPP assay (**Fig. 5c**). We first discovered that the original NLuc substrate furimazine was insufficiently stable for HTS, as we observed a time-dependent erosion of its signal intensity over the ∼10 min time period needed to scan a full 384-well plate on a ClarioStar instrument. Switching to an alternative substrate vivazine generated luminescence signal that was stable for ∼10 h. We also extended the time course analysis of NanoBRET-ABPP reactions between YZ-01-A and the N112C-CCNE1:CDK2 complex to 185 min, which produced strong stereoselective BRET signals without showing evidence of reaction completion over the incubation time period (**Fig. 5d**). We selected a time point for HTS of 170 min, at which we observed excellent Z’ (0.85) and signal-to-noise (∼30) values in 384-well plate format (**Fig. 5d** and **Extended Data Fig. 11a**). Using this optimized NanoBRET-ABPP assay, we first verified the relative potency of engagement of N112C-CCNE1:CDK2 complexes by I-125A and 1-198 (**Fig. 5e**), as well as the stereoselective reactivity of WX-02-308 with N112C-CCNE1 (**Extended Data Fig. 11b**). As a proof-of-principle, we then performed a pilot screen of 324 compounds from the ChemBridge macrocycles collection (https://chembridge.com/screening-compounds/macrocycles/#) tested at 10 µM alongside two concentrations of I-125A, and this screen proved capable of “rediscovering” I-125A as a hit ligand for N112C-CCNE1 (**Fig. 5f**). We accordingly expect that the optimized NanoBRET-ABPP assay will be suitable for larger HTS to identify additional types of chemistry binding to the allosteric pocket in CCNE1:CDK2 complexes.

## Discussion

Paralogous proteins, due to their prevalence, structural relatedness, and overlapping, but distinct functions, present fascinating opportunities for chemical biology. For instance, small-molecule probes can be developed that target paralogous proteins with a range of tailored selectivity profiles. This has been achieved with groups of related kinases for which pan- and paralog-restricted inhibitors have been found to produce different pharmacological effects on immune^48,14,49,15^ and cancer^50–54^ signaling pathways.

The discovery of orthosteric chemical probes for paralogous proteins has often relied on comparative structural biology methods using purified proteins or protein domains^55,50,52^. Such structure-guided approaches, however, may overlook opportunities for identifying paralog-restricted chemical probes acting by allosteric mechanisms that involve intact proteins functioning in more native biological environments (e.g., as parts of physiological complexes or pathways). Allosteric, paralog-restricted inhibitors targeting the pseudokinase domains of TYK2 and JAK1, have alternatively been discovered by phenotypic screening^56^ and cysteine-directed ABPP^15^, respectively. These findings underscore the diverse and sometimes serendipitous ways that chemical probes for paralogous proteins have been discovered.

The incorporation of non-native cysteine residues into proteins to confer sensitivity to electrophilic compounds also has a rich history for chemical probe discovery^57–62^. Nonetheless, the selection of sites for introducing the cysteine has, in most cases, relied on structural knowledge of an already mapped small molecule-binding or functional pocket. The paralog hopping strategy described herein is complementary and potentially generalizable in that it does not require prior structural information, but instead leverages public chemical proteomic datasets containing many covalently liganded paralog-restricted cysteines (e.g., > 60 such cysteines are described in ref. ^24^). In the case of CCNE1 and CCNE2, this allowed for the discovery of a cryptic ligandable pocket that was nonobvious from apo-structures of CCNE1-CDK2 complexes. The successful engineering of covalent ligandability into ‘cysteine-less’ pockets in related proteins can then leverage various cysteine-directed assays for compound screening, including, in addition to NanoBRET-ABPP, tethering^57^ and and fluorescence polarization-ABPP^34^. NanoBRET-ABPP has the advantage of being compatible with assaying proteins in complex proteomes, as shown here for CCNE1:CDK2 complexes, as well as in living cells, as has recently been demonstrated with activity-based probes for caspases^63^.

The proximity of N112 of CCNE1 to the binding site for I-125A and I-198, as well as the blockade of stereoprobe interactions with N112C-CCNE1 by these reversible inhibitors, suggests that the different classes of CCNE1 ligands studied herein bind to the same pocket. We cannot, however, exclude the possibility of partly overlapping or non-overlapping proximal pockets for the covalent and non-covalent N112C-CCNE1 ligands. Indeed, we do not yet understand why I-125A and I-198 incompletely block stereoprobe reactivity with N112C-CCNE1 or why such partial inhibition was also observed for I-125A and I-198 with the Histone H1 substrate, but not an RB1 peptide substrate for both WT- and N112C-CCNE1. The potential for for such differences to be caused by heterogeneity in the modification state or broader protein interactions of CCNE1:CDK2 complexes isolated from human cells that in turn influence ligand binding or activity is an important topic for future investigation.

Some limitations of the paralog hopping strategy should be discussed. Depending on the structural conservation across paralogs, it is possible that a ligandable pocket will only be found in a subset of related proteins. We are encouraged by our success with CCNE1 and CCNE2 in that these proteins only share ∼50% overall sequence identity; however, our structural studies revealed a surprising degree of conservation for residues contacting I-125A/I-198, and it remains an open question whether paralog hopping can be accomplished with proteins displaying lower sequence/structural relatedness. We also acknowledge that covalent chemistry has advantages for targeting shallow and dynamic pockets in proteins, and such sites in cysteine-less paralogs may prove more difficult to address with reversible chemistry.

Nonetheless, we are encouraged by recent examples of success in progressing from covalent to non-covalent ligands for challenging protein targets, as has been described for oncogenic mutant forms of KRAS^64,65^. From a technical perspective, our paralog hopping method currently uses heterologously expressed proteins, but we envision the potential to extend the strategy to endogenously expressed proteins by leveraging, for instance, base editing^66^ to introduce codons encoding cysteine residues into the genome.

Finally, the different functional effects displayed by 2,6-diazaspiro[3.4]octane and tryptoline acrylamide ligands targeting N112C-CCNE1 – inhibition of CDK2 and stabilization of CKS1B/CKS2 interactions, respectively – underscore the potential for allosteric sites to modulate distinct biochemical activities of proteins^67–69^. While we do not yet understand how promoting interactions with CKS1B/CKS2 may impact the function of CCNE1:CDK2 complexes, it is possible that tryptoline acrylamides displaying this property may function as substrate-selective agonists in cells^45,46^. More generally, the diverse functional effects conferred by allosteric pockets points to the value of fully exploring their small molecule interaction potential. We further speculate that ‘binding-first’ assays^70^ such as NanoBRET-ABPP may represent a preferred approach for screening allosteric pockets, as they should be capable of identifying hit compounds displaying a broad range of functional activities, including antagonists, agonists, and silent ligands. Indeed, other binding-first assays, such as structure-guided fragment screening, have been used to identify ligands for allosteric sites that are initially silent and then converted through medicinal chemistry into functional inhibitors^71^. We anticipate that the application of paralog hopping to allosteric sites will offer a powerful strategy for expanding chemical probe discovery throughout the proteome.

## Supporting information

Supplementary Chemistry Information

PDB Validation Reports

## Data Availability

Proteomic data are available via ProteomeXchange with identifier PXD048328. The atomic coordinates and structure factors have been deposited in the Protein Data Bank, www.pdb.org (PDB ID codes 8VQ3 (I-125A-CCNE1:CDK2 structure) and 8VQ4 (I-198-CCNE1:CKD2 structure)).

## Acknowledgements

This work was supported by the NCI (R35 CA231991) and Pfizer. We thank You-Ai He for purification of recombinant CCNE1:CDK2 complexes used for crystallography. We thank Travis J. Matthewson for preparing compound plates for the NanoBRET-ABPP high-throughput screen.

## Methods

### Protein expression and purification

Full-length human CDK2 gene (UniProt ID: P24941) was codon optimized for insect cell expression, synthesized de novo, and subcloned into a modified pFastBac vector containing no affinity tags. Full-length human CCNE1 gene (UniProt ID: P24864) was codon optimized for insect cell expression, synthesized de novo, and subcloned into a modified pFastBac vector containing N-terminal Flag affinity tag followed by a Tobacco Etch Virus (TEV) protease cleavage site. Full-length human CCNE2 gene (UniProt ID: O96020) was codon optimized for insect cell expression, synthesized de novo, and subcloned into a modified pFastBac vector containing N-terminal hexahistidine tag followed by a Tobacco Etch Virus (TEV) protease cleavage site. CCNE1_N112C and CCNE2_C109A mutants were generated by site directed mutagenesis of the wild-type plasmids. In order to recombinantly express the CDK2:CCNE1 or CDK2:CCNE2 heterodimers, a 2 liter culture of Spodoptera frugiperda (Sf21) insect cells at a density of 2e6/mL was infected with equal volumes of high-titer stocks of recombinant baculovirus generated from each of the above plasmids, and allowed to grow in a shake flask for 72 hours at 27 ⁰C. Cell pellets were harvested by centrifugation, and frozen at -80 ⁰C for at least 24 h prior to purification.

All purification steps were performed at 4 ⁰C. Cells were resuspended in lysis buffer containing 50 mM Tris [pH=8.0], 250 mM NaCl, 0.25 mM Tris(2-carboxyethyl)phosphine hydrochloride (TCEP), 1/75 mL Roche EDTA-free protease inhibitor tablets, 10 µM leupeptin, and 10 µM E-64 protease inhibitor, then clarified by centrifugation. Binary CDK2:CCNE2 protein complexes were purified using nickel ProBond resin (Life Technologies). N-terminal poly-histidine tag was subsequently cleaved off CCNE2 using TEV protease, and protein sample passed over nickel Probond resin again, retaining flow-through fractions. Binary CDK2:CCNE1 protein complexes were purified using anti-Flag resin (Pierce). All protein samples were concentrated and passed over a gel filtration chromatography column (SRT-500, Sepax Technologies) equilibrated in 25 mM HEPES pH 7.5, 250 mM NaCl, and 0.25 mM TCEP. Peak fractions were pooled, concentrated to 1-2 mg/mL, flash frozen and stored at -80 ⁰C until further use.

### Mass Spectrometry Reactivity Assay (MSRA)

3 µM thawed protein samples in reaction buffer (25 mM HEPES [pH=7.5], 250 mM NaCl, 0.25 mM TCEP) were incubated for 3 h at ambient temperature in the presence of 50 µM of each ligand. Reactions were quenched with 0.5% formic acid (FA). A Waters Acquity UPLC system equipped with RP column (Opti-TRAP MICRO 5uL, Optimize Technologies Inc.) was used to separate protein samples. 10 µL aliquot of each sample was injected onto the instrument, and a 2 min 10%-30% acetonitrile (ACN) gradient in water containing 0.2% FA at 0.6 mL/min was used to elute protein samples. LC/MS analysis of eluted samples were performed using Waters Xevo G2-XS system. The instrument was operated in positive ion mode, with capillary and cone voltage set to 3 kV and 40 V, respectively. The source temperature was set to 100 ⁰C, and nitrogen was used as the desolvation gas at 800 L/h, 300⁰C. LC/MS data was processed using Waters MassLynx software, which calculated deconvoluted mass and peak intensities for each protein species. Peak intensities were used to approximate relative species abundance.

### DNA constructs and transfection

WT-CCNE1-3xFLAG (GenScript, Lot:U8948FB050-2/PD40693), N112C-CCNE1-3xFLAG (GenScript, Lot:U8948FB050-6/PD43863) plasmids used for mammalian cell over-expression were purchased from GenScript. NLuc-WT-CCNE1-3xFLAG, NLuc-N112C-CCNE1-3xFLAG plasmids were cloned into PvuI and EcoRI digested NLuc-CDK4 plasmid (Promega) using gBlocks obtained from IDT. All sequences were verified via next-gen sequencing (Primordium) before use. Untagged CDK2 expression plasmid was cloned by PCR with CDK2-NLuc plasmid (Promega) as the template. Transfections were performed using polyethylenimine (PEI-MAX, Polysciences, 24765).

### Antibodies

For western blotting, CDK2 mouse monoclonal antibody (HRP conjugated) [Clone ID: OTI2D9] (Origene, TA502935BM), monoclonal Anti-FLAG M2-peroxidase (HRP) (produced in mouse) (Sigma-Aldrich, A8592).

### Tissue culture

HEK293T (ATCC, CRL-3216) cells were maintained in DMEM supplemented with 10% vol/vol fetal bovine serum (FBS), 2 mM L-alanyl-L-glutamine (GlutaMax, Gibco, 35050061), penicillin (100 U mL^-1^), and streptomycin (100 μg mL^-1^).

### Gel-based ABPP experiments

For gel-based ABPP experiments using purified CCNE:CDK2 complexes, 1 µM CCNE:CDK2 complexes were treated with different compounds for desired period of time, then chased with 5 µM 1 h alkyne probe treatment at room temperature. Compound stocks were made in DMSO and final DMSO concentration did not exceed 1% vol/ vol. A 10x click master mix containing 125 µM rhodamine-azide, 10 mM TCEP, 1 mM Tris((1-benzyl-4-triazolyl)methyl)amine (TBTA; in 4:1 t-BuOH:DMSO) and 10 mM CuSO4 was made and added. After 1 h incubation, 4x laemmli loading dye was added. Samples were analyzed by SDS-PAGE.

For gel-based ABPP experiments using HEK293T cell lysate, HEK293T cells transiently expressing NLuc-WT-CCNE1 or NLuc-N112C-CCNE1 and CDK2 were harvested 48 h post transfection and stored at -80 °C. Cells were then lysed in DPBS by probe sonication (2x 15 pulses) and whole-cell lysate was normalized with Protein BCA assay (Pierce) to 2 mg mL^-1^. 500 µL normalized whole-cell lysate was treated with 10 µM YZ-01-A or 10 µM YZ-01-B for 90 min, then 50 µL 10% IGEPAL (in DPBS) was added and lysate was clarified by 16,000*g* 5 min centrifugation. Supernatant was taken and mixed with 20 µL pre-washed anti-DYKDDDK magnetic agarose bead slurry (Pierce) and rotated at 4 ℃ for 2 h. Beads were washed four times with wash buffer (DPBS, 0.2% IGEPAL) and once with DPBS, then protein was eluted by adding 50 µL 1x laemmli loading dye. Samples were analyzed by SDS-PAGE gel.

### CCNE1 NanoBRET-ABPP assays

The CCNE1 NanoBRET-ABPP assay was performed in a 384-well format (Greiner, 784075). Briefly, HEK293T cells transiently expressing NLuc-N112C-CCNE1 or NLuc-WT-CCNE1, and CDK2 were harvested 48 h post transfection and stored at -80 °C. Cells were lysed in BRET assay buffer (50 mM HEPES [pH = 7.5], 150 mM NaCl) by probe sonication (2x 15 pulses) and whole-cell lysate was normalized to 2 mg mL^-1^ with Protein BCA assay (Pierce). After normalization, lysate was mixed with YZ-01-A or YZ-01-B (final DMSO 1% vol/ vol) to 10 µM final concentration, then dispensed to 384-well plates pre-plated with tested compound and allowed for 90 min room-temperature incubation. 4 µL furimazine (0.36% vol/ vol diluted in BRET assay buffer, Promega) was added before readout on ClarioStar.

For CCNE1 NanoBRET-ABPP HTS-compatible assays, a similar protocol was followed with modifications. Normalized cell lysate was first mixed with vivazine (Promega) to reach 1% vol/vol as recommended by Promega, then mixed with YZ-01-A (final DMSO 1% vol/ vol) to 10 µM final concentration, then dispensed to 384-well plates pre-plated with 4 mM 0.05 µL tested compounds(10 µM final concentration) and allowed for 3 h room-temperature incubation. BRET signal was read out on ClarioStar.

### ADP-Glo kinase assays with purified CCNE:CDK2 complexes

The CDK2 activity assay was performed in a 384-well format (Greiner, 784075) using ADP-Glo Kinase Assay (Promega). Briefly, different CCNE: CDK2 purified complexes at 4 ng µL^-1^ were mixed 1:1 with 2x compound stock (Both are made in kinase reaction buffer (40 mM Tris [pH = 7.6], 20 mM MgCl2, 0.1 mg mL^-1^ BSA and 50 µM DTT) and incubated at 37 ℃ for 1 h. Then 6 ng compound treated CCNE: CDK2 complexes (3 µL) were mixed with 2µL ATP/ CDK2 substrate mix to make final concentration of 150 µM ATP and 0.5 µg substrate per reaction (Substrate used is either histone H1(Sigma-Aldrich, 14-155) or RB1 C-terminal peptide (MyBioSource, MBS143254)), then incubated at room temperature for 1 h. Subsequently, 5 µL ADP-Glo reagent was added and incubated for 40 min at room temperature, followed with 10 µL kinase glo reagent and another 30-min incubation. Luminescence was read out on ClarioStar.

### I-125A and I-198 CCNE1: CDK2 co-crystal structures

#### Protein production and purification

The CDK2/CCNE1 complex for crystallography was generated as previously described^13^ with modifications. Briefly, the cDNAs encoding human full-length CDK2 and residues 96–378 of human CCNE1 with an N-terminal His tag were synthesized and cloned into modified pFastBack1 vectors, and then co-expressed in sf21 cells at a ratio of 1:1. Harvested cells were resuspended in lysis buffer (40 mM HEPES, pH 7.5, 150 mM NaCl, 0.01% 1-thioglycerol, and 25 mM imidazole), and lysates were cleared by centrifugation. The complex was bound on Probond (Invitrogen) resin followed by lysis buffer wash, elution at 250 mM imidazole, TEV protease digestion and overnight dialysis. The protein was again flowed over Probond resin to remove cleaved His tags, and His-tagged TEV. The complex was then subjected to size exclusion chromatography using a HiLoad 26/600 Superdex 200 pg column equilibrated in 40 mM HEPES, pH7.5, 150 mM NaCl, 0.01% 1-thioglycerol. Intact mass analysis indicated the complex was >95% phosphorylated as isolated and required no additional activation.

#### Crystallization and crystallography

The phosphorylated CDK2/CCNE1 complex (8.3 mg/mL) was crystallized in MRC2 96-well sitting drops plates by mixing 200 nL protein plus 200 nL well solution (0.21 M sodium citrate, 19.1% PEG 3350) followed by incubation at 30°C overnight and further incubation at 21 °C. Crystals growth was completed in 2-4 days and apo crystals were subjected to soaking in 1mM of the I-125A or I-198 for ∼3 days, followed by cryoprotection with well solution plus 20% glycerol and flash-freezing in liquid N2. X-ray diffraction data were collected at the Advanced Photon Source beamline 17-ID and processed with autoPROC3 using STARANISO4 from Global Phasing. Structure solution and refinement were done using BUSTER5, PHENIX6 and COOT7 and figures were generated using PYMOL.

### CCNE1 IP-MS studies

C-terminally 3xFLAG epitope-tagged CCNE1 WT or N112C mutant were transiently co-expressed with untagged CDK2 in HEK293T cells. 48 h post-transfection, cells were replenished with fresh media and treated with different compounds (final DMSO 0.1 % vol/ vol) for 6 h. Cells were collected and washed with ice cold DPBS and stored at -80 °C. Frozen cell pellets were lysed in immunoprecipitation lysis buffer (50 mM Tris [pH = 7.5], 150 mM NaCl, 1% IGEPAL, 10% Glycerol) with cOmplete Protease Inhibitor Cocktail (Roche) and PhosSTOP Phosphatase Inhibitor Cocktail (Roche) by probe sonication (2x 15 pulses). The insoluble debris was removed by centrifuging at 16, 000*g* for 5 min. The supernatant was taken and normalized to 2 mg mL^-1^ with Protein BCA assay (Pierce). 2 mg proteome per sample was used to incubate with 40 µL anti-DYKDDDDK Magnetic Agarose bead slurry (washed twice with lysis buffer) for 3 h at 4 °C. Beads were then spun down at 2,000*g* for 2 min, washed four times with immunoprecipitation wash buffer (DPBS, 0.2% IGEPAL) and then once with DPBS. For I-125A treated samples, 1 µM I-125A was included in the lysis and wash buffers. The samples were eluted from beads using 50 µL 8 M urea in DPBS and 10-min 65 ℃ incubation, then reduced with 200 mM DTT at 65 ℃ for 15 min and alkylated with 400 mM iodoacetamide at 37 °C for 30 min. After diluted to 2 M Urea by the addition of DPBS, samples were digested using MS-grade trypsin (Promega) at 37 ℃ overnight, followed by TMT labeling (Thermo Scientific) and desalting using Peptide Desalting Spin Columns (Pierce) as described previously. The labeled peptides were analyzed in Orbitrap Fusion Mass Spectrometer (ThermoFisher).

---

**Extended Data Figure 1.**
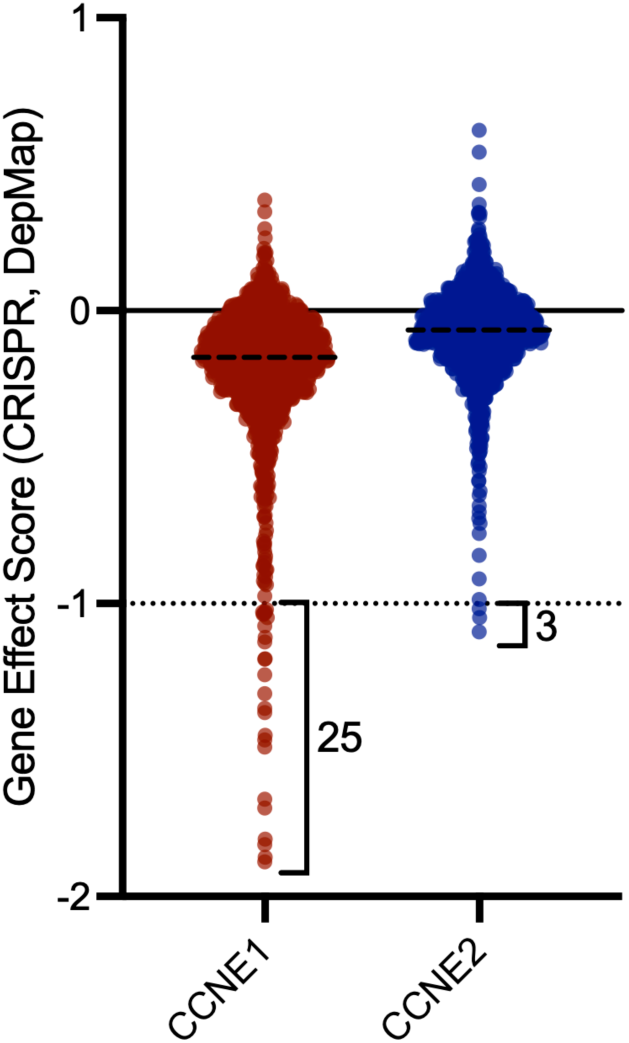
Cancer Dependency Map data for CCNE1 and CCNE2, indicating the number of human cancer cell lines showing gene effect scores < -1.0, reflecting strong dependency on the gene^31^.

**Extended Data Figure 2.**
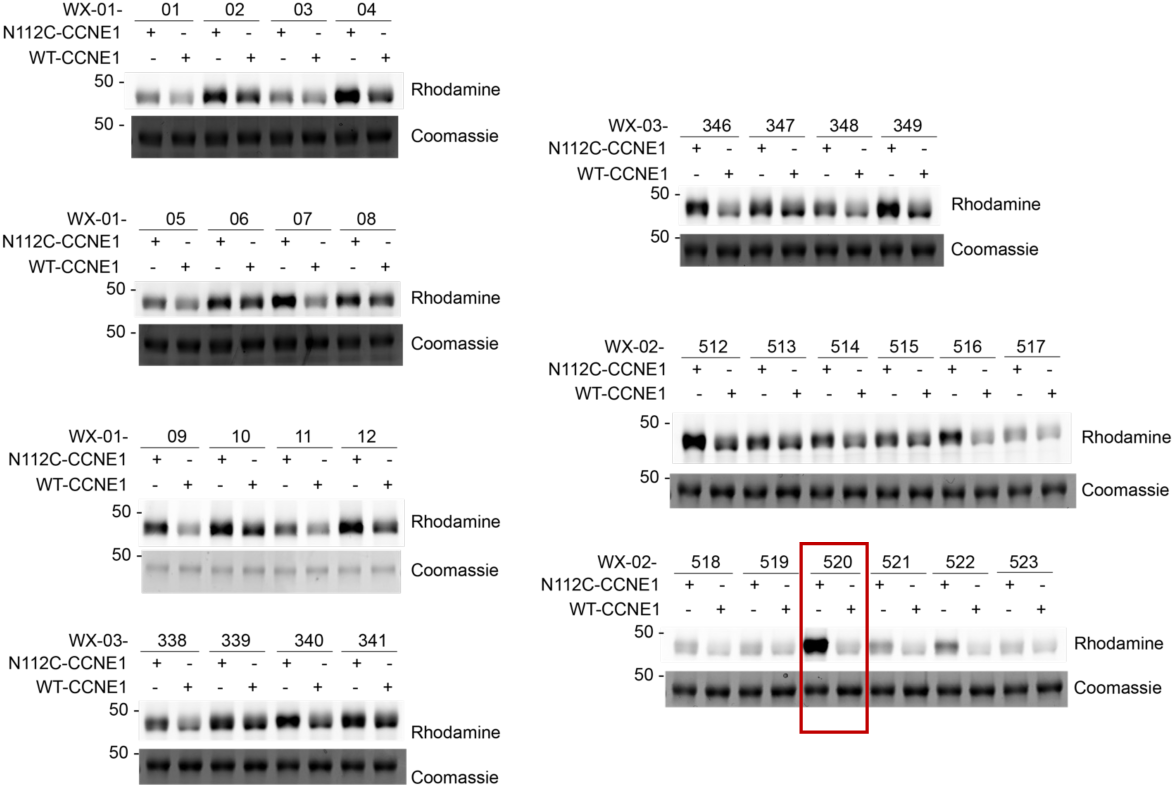
Gel-ABPP data for a N112C- and WT-CCNE1:CDK2 complexes treated with a focused library of alkyne-modified tryptoline acrylamides. Gel-ABPP data for purified N112C- or WT-CCNE1:CDK2 complexes (1 µM) exposed to alkyne-modified tryptoline acrylamides (5 µM, 1 h) followed by CuAAC conjugation to an Rh-N_3_ tag, SDS-PAGE, and in-gel fluorescence scanning. Coomassie blue signals correspond to WT- or N112C-CCNE1. Red box marks profile of WX-02-520, which reacts with N112C-CCNE1 in a stereoseletive (compared to enantiomer WX-02-521) and site-specific (compared to WT-CCNE1) manner.

**Extended Data Fig. 3.**
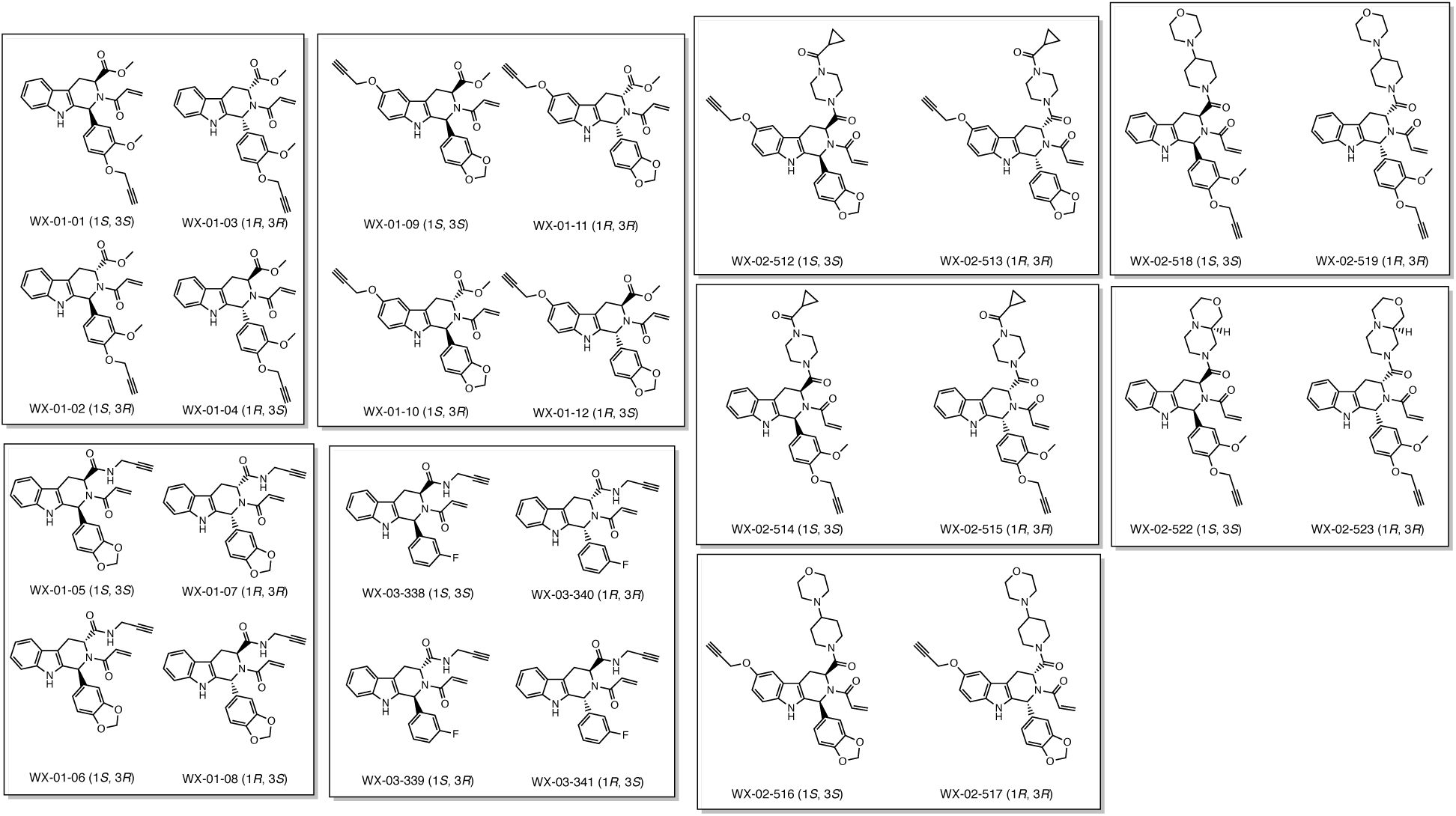
Structures of alkyne-modified tryptoline acrylamides screened for reactivity with N112C-CCNE1:CDK1 complexes.

**Extended Data Figure 4.**
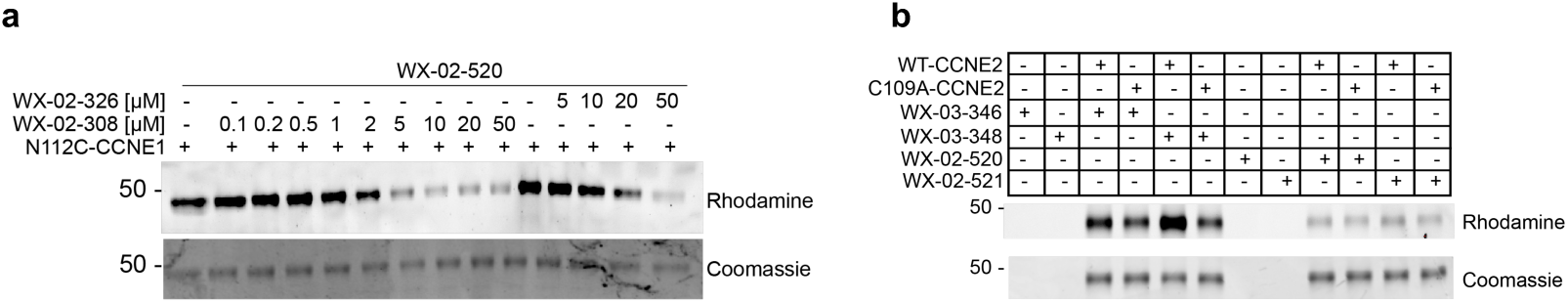
N112C-CCNE1 and WT-CCNE2 show differential reactivity profiles with tryptoline acrylamides. **a**, Gel-ABPP data showing concentration-dependent, stereoselective blockade of WX-02-520 engagement of purified N112C-CCNE1 by WX-02-308 in comparison to enantiomer WX-02-326. Purified N112C-CCNE1:CDK2 complex (1 µM) was pre-treated with the indicated concentrations of WX-02-308 or WX-02-326 (2 h) followed by WX-02-520 (1 µM) and processing for gel-ABPP. Data are from a single experiment. **b**, Gel-ABPP data showing stereoselective engagement of WT-CCNE2, but not C109A-CCNE2 by WX-02-346 in comparison to enantiomer WX-02-348 or the WX-02-520 and WX-02-521 stereoprobes. Purified WT- or C109A-CCNE2:CDK2 complexes were treated with the indicated concentrations of alkyne tryptoline acrylamides followed by processing for gel-ABPP.,

**Extended Data Figure 5.**
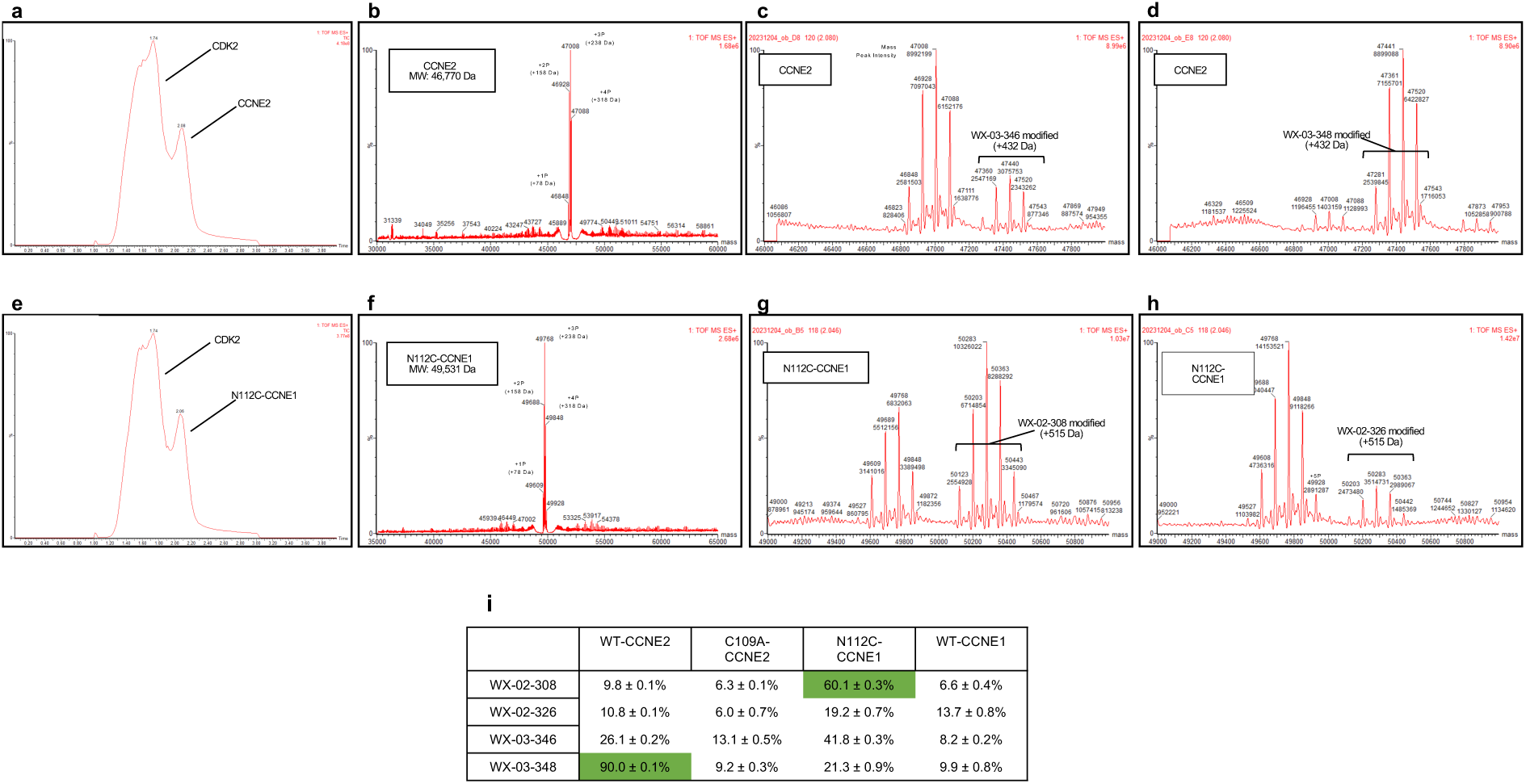
Mass spectrometry (MS) analysis of tryptoline acrylamide stereoprobe reactivity with purified CCNE2- and N112C-CCNE1:CDK2 complexes. a, b, Reverse-phase liquid chromatography-MS traces for purified recombinant WT-CCNE2 (**a**) and N112C-CCNE1 (**b**) complexes with CDK2. **c-e**, Deconvoluted intact mass analysis of the peak mass spectrum for CCNE2 with no compound treatment (**c**), WX-02-346 treatment (**d**), and WX- 02-348 treatment (**e**). Multiple phosphorylation states can be assigned to each peak based on expected mass and mass shifts (+80 Da / phosphate). **f-h**, Deconvoluted intact mass analysis of the peak mass spectrum for N112C-CCNE1 with no compound treatment (**f**), WX-02-308 treatment (**g**), and WX-02-326 treatment (**h**). For c-h, each peak is annotated with mass (top) and peak intensity (bottom). Peak intensities were used to semi-quantitatively assess relative species abundance (shown in table in **i**). Multiple phosphorylation states for each protein can be assigned to each peak based on expected mass and mass shifts (+80 Da / phosphate). **i,** Summary of results listing relative abundance of each modified protein species as percent of total across the four tested stereoprobes. Also included are results from experiments performed with control proteins (C109A-CCNE2 and WT-CCNE1) that show only minor and non-stereoselective reactivity with the tested stereoprobes. Green highlights values where substantial (> 50%) and stereoselective (>3X over enantiomer) stereoprobe engagement of the indicated CCNE protein was observed.

**Extended Data Figure 6.**
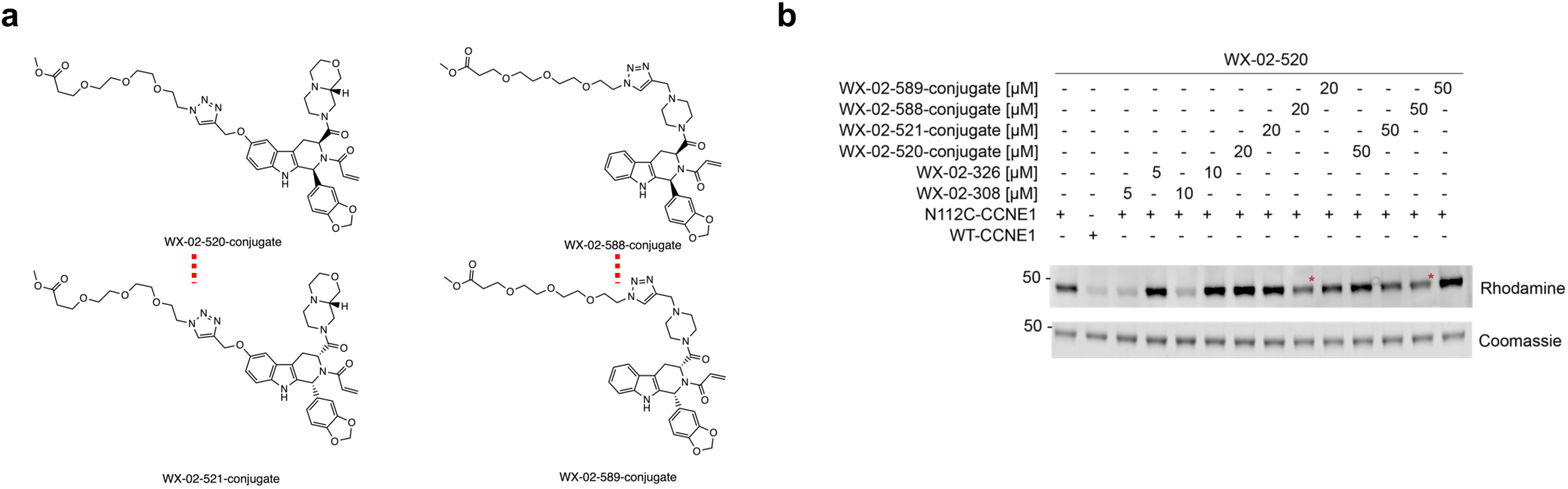
Reactivity of PEG-conjugated tryptoline acrylamides with N112C-CCNE1. a, Structures of PEG-conjugated tryptoline acrylamide stereoprobes. **b**, Gel-ABPP data showing stereoselective blockade of WX-02-520 engagement of purified N112C-CCNE1 by WX-02-588-conjugate in comparison to enantiomer WX-02-589-conjugate or additional stereoprobes WX-02-520-conjugate and WX-02-521-conjugate. Purified N112C-CCNE1:CDK2 complex (1 µM) was pre-treated with the indicated concentrations of PEG-conjugated stereoprobes (2 h) followed by WX-02-520 (1 µM, 1 h) and processing for gel-ABPP. Red asterisks mark decreased WX-02-520 reactivity with N112C-CCNE1 in samples pre-treated with WX-02-588-conjugate.

**Extended Data Fig 7.**
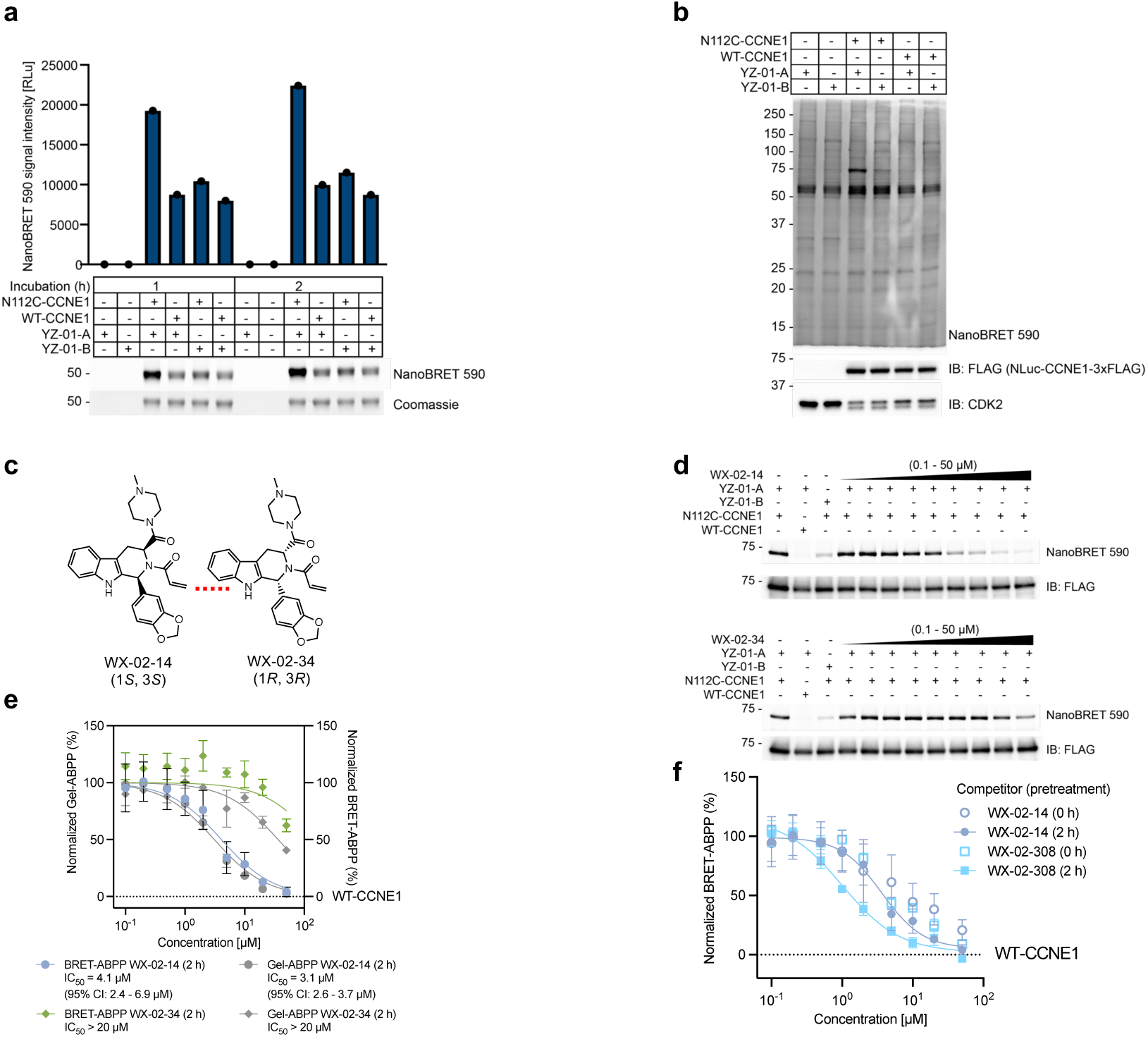
Characterization of YZ-01-A interactions with N112C-CCNE1. **a**, Gel-ABPP data showing that YZ-01-A stereoselectively engages purified N112C-CCNE1:CDK2 but not WT-CCNE1:CDK2 complexes. The purified N112C- or WT-CCNE1:CDK2 complexes were incubated with YZ-01-A or YZ-01-B (5 µM, 1 or 2 h) followed by processing of samples for gel-ABPP. Data are from a single experiment. **b**, Gel-ABPP data showing stereoselective engagement of N112C-CCNE1, but not WT-CCNE1 by YZ-01-A in lysates from HEK293T cells recombinantly co-expressing NLuc-N112C-CCNE1-3xFLAG (or NLuc-WT-CCNE1-3xFLAG) and CDK2 were treated with YZ-01-A or YZ-01-B (10 µM, 90 min) followed by processing of samples for gel-ABPP. Data are from a single experiment. **c**, Structures of WX-02-14 and WX-02-34. **d**, Gel-ABPP data showing concentration-dependent stereoselective blockade of YZ-01-A engagement of N112C-CCNE1 by WX-02-14 in comparison to WX-02-34. Lysates from HEK293T cells co-transfected with NLuc-N112C-CCNE1 or NLuc-WT-CCNE1 and CDK2 were treated with the indicated concentration ranges of compounds followed by YZ-01-A (10 µM, 90 min), anti-FLAG immunoprecipitation, and processing for gel-ABPP. **e**, Comparison of gel- and NanoBRET-ABPP data showing concentration-dependent, stereoselective blockade of YZ-01-A engagement of NLuc-N112C-CCNE1 by WX-02-14 in comparison to WX-02-34 (0.2 – 50 µM, 2 h). Dashed horizontal line marks background signals for NanoBRET-ABPP assays performed with YZ-01-A (10 µM, 90 min) and NLuc-WT-CCNE1. CI, confident intervals. Data represent average values ± s.e.m., n = 3. **f**, NanoBRET-ABPP data showing a time-dependent increase in potency of blockade of YZ-01-A reactivity with N112C-CCNE1 by WX-02-14 and WX-02-308. Data represent average values ± s.e.m., n = 3.

**Extended Data Fig. 8.**
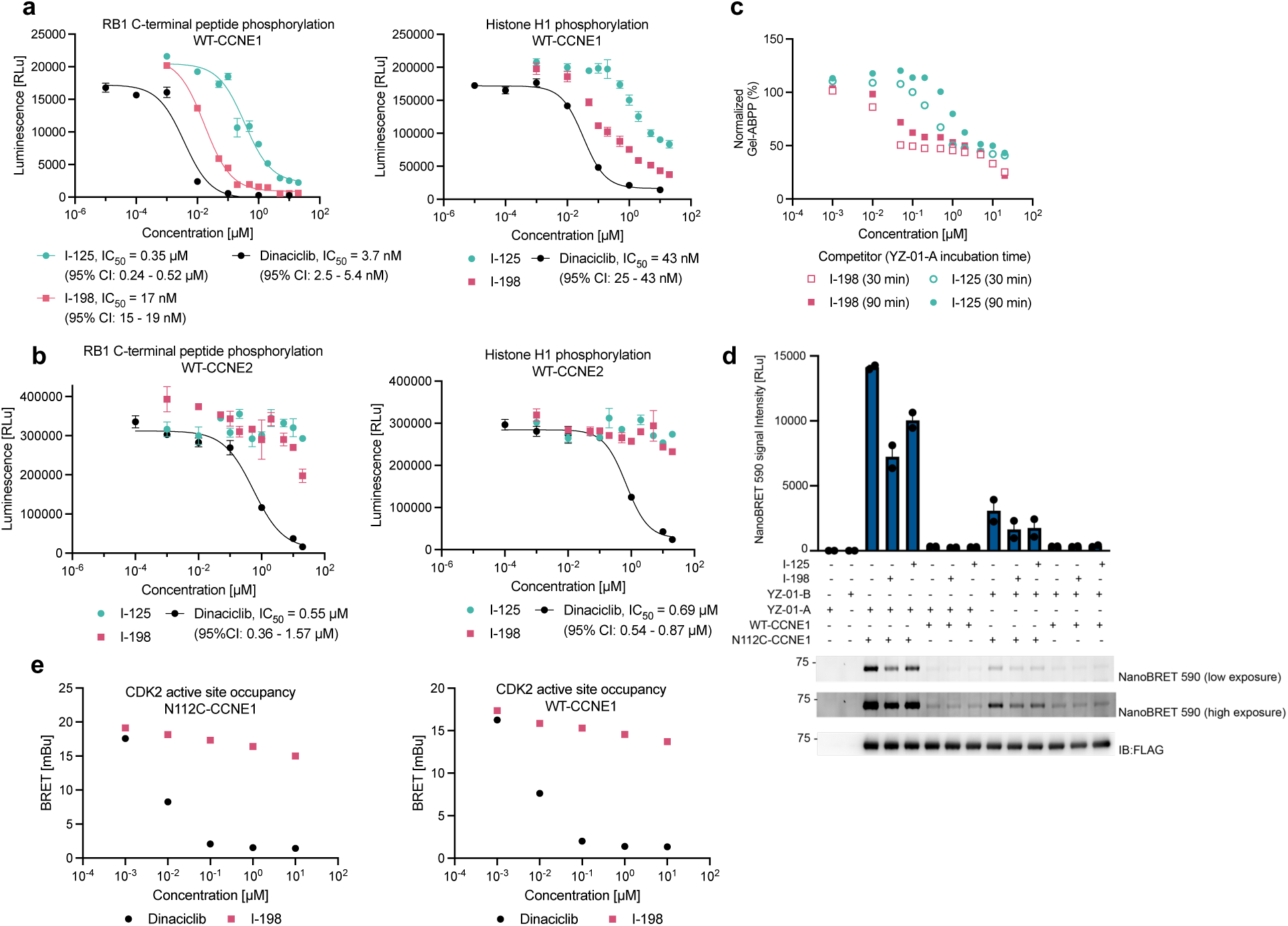
Additional characterization of allosteric inhibitors of CCNE1:CDK2 complexes. **a, b**, ADP-Glo data showing effects of the indicated compounds on the activity of a purified WT-CCNE1:CDK2 (**a**) or WT-CCNE2:CDK2 (**b**) complexes as measured by phosphorylation of the C-terminal peptide of RB1 (left) or the histone H1 protein (right). The WT-CCNE:CDK2 complexes were pre-treated with compounds for 2 h (37 °C) before initiating phosphorylation reactions by addition of substrate. CI, confident intervals. Data represent average values ± s.e.m., n = 3. **c,** Gel-ABPP data showing concentration-dependent blockade of YZ-01-A engagement of N112C-CCNE1 following incubation with I-125A or I-198 for 30 min or 90 min. Experiment was performed as described in Fig. 3c. Data are from a single experiment. **d**, Gel-ABPP data showing that residual YZ-01-A reactivity with N112C-CCNE1 in presence of 1 µM of I-125A or I-198 is stereoselective (greater than signals generated with YZ-01-B) and site-specific (greater than signals generated with WT-CCNE1). Experiments were performed as described in Fig. 3c. Data represent average values ± s.e.m., n = 2. **e**, NanoBRET data showing concentration-dependent blockade of active site-directed tracer K-10 (0.2 µM) binding to N112C-CCNE1:CDK2-NLuc (left) or WT-CCNE1:CDK2-NLuc complexes by dinaciclib, but not I-125A. NanoBRET assays were performed in lysates of HEK293T cells recombinantly expressing the CCNE1:CDK2-NLuc complexes. Data represent average values ± s.e.m., n = 3.

**Extended Data Fig. 9.**
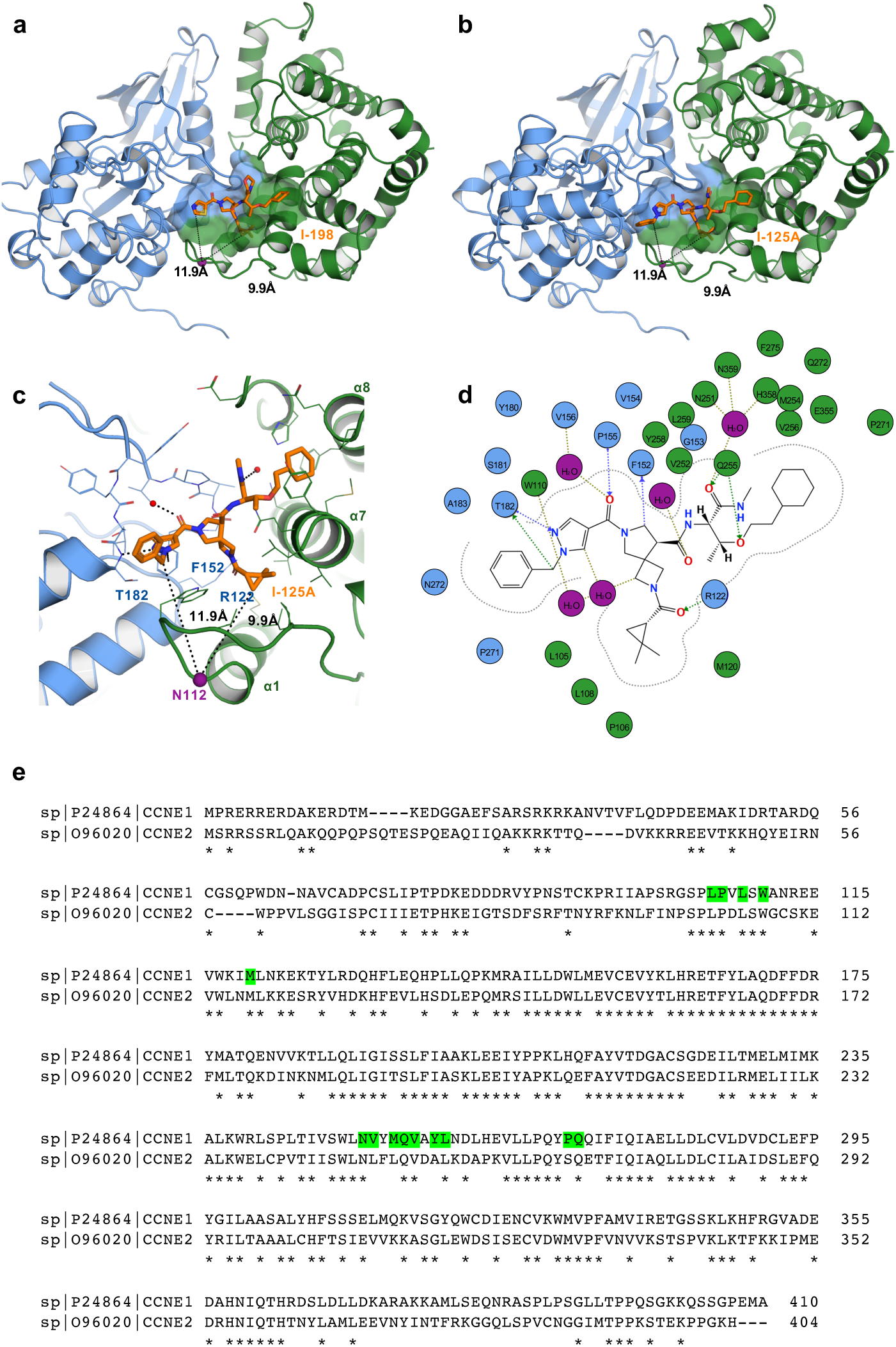
Crystal structures of I-125A and I-198-CCNE1:CDK2 complexes. **a**, Ribbon diagram structure of CCNE1:CDK2 (green: blue) complex with the I-198 compound shown in orange and the binding cavity shown as a transparent surface. **b**, Ribbon diagram structure of CCNE1:CDK2 (green: blue) complex with the I-125A compound shown in orange. For **a, b**, N112 is shown in purple and the distance from this residue to ligands indicated by dashes. **c**, Detailed view of the I-125A binding site, protein-ligand interactions, and the distance to N112 indicated by dashes. CCNE1 and CDK2 residues are shown in green and blue, respectively, and water molecules are in magenta. **d**, Protein-ligand interaction plot between CCNE1: CDK2 and I-125A. **e**, A global sequence alignment of CCNE1 and CCNE2. CCNE1 residues shown in Figure 4c and **Extended Data Fig. 9d** are highlighted in green.

**Extended Data Figure 10.**
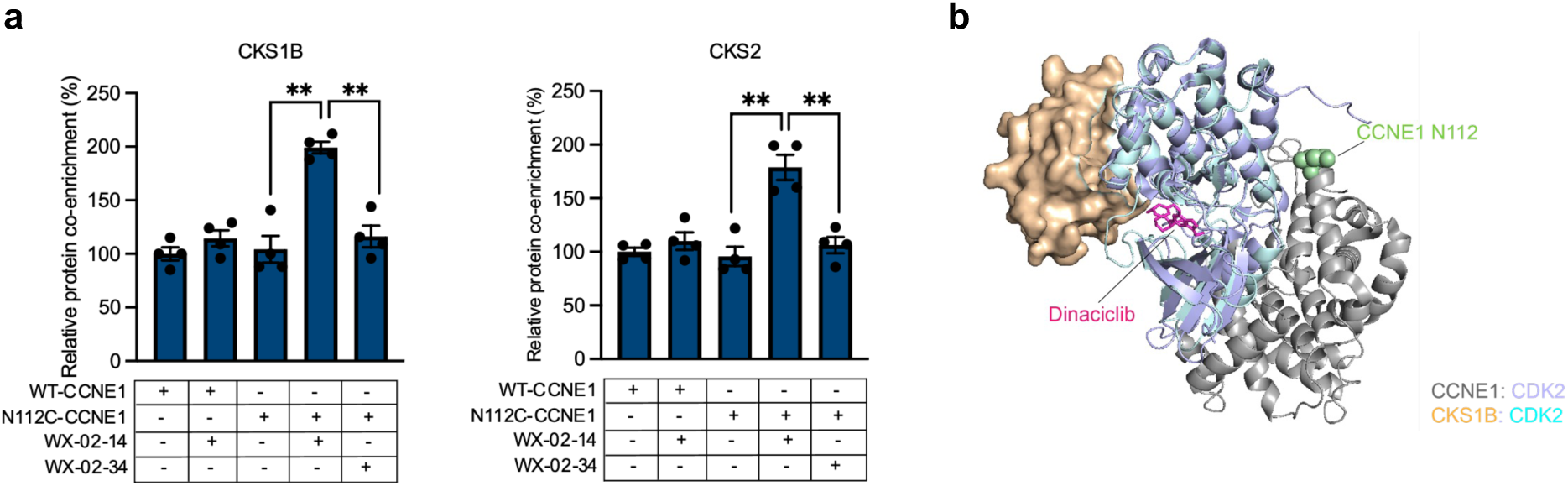
Impact of tryptoline acrylaimdes WX-02-14 and WX-02-34 on protein interactions of CCNE1:CDK2 complexes. **a**, Quantification of data for CKS1B and CKS2 in co-immunoprecipitation-MS experiments performed in HEK293T cells recombinantly expressing N112C-CCNE1-3xFLAG (or WT-CCNE1-3xFLAG) and CDK2 treated with WX-02-14 or WX-02-34 (20 µM, 6 h) or DMSO. Data represent average ± s.e.m., n = 4. Unpaired t test with Welch’s correction was used to test statistical significance. p values: 0.0011 and 0.0021 for CKS1B in WX-02-14-treated vs DMSO-treated and WX-02-34-treated N112C-CCNE1-expressing cells, respectively; 0.0017 and 0.0032 for CKS2 in WX-02-14-treated vs DMSO-treated and WX-02-34-treated N112C-CCNE1-expressing cells, respectively. **b,** Overlay of crystal structures of CKS1B:CDK2^47^ (PDB: 1BUH) and CCNE1:CDK2^74^ (PDB: 5L2W) showing the relative locations of N112 of CCNE1 and CKS1B.

**Extended Data Fig. 11.**
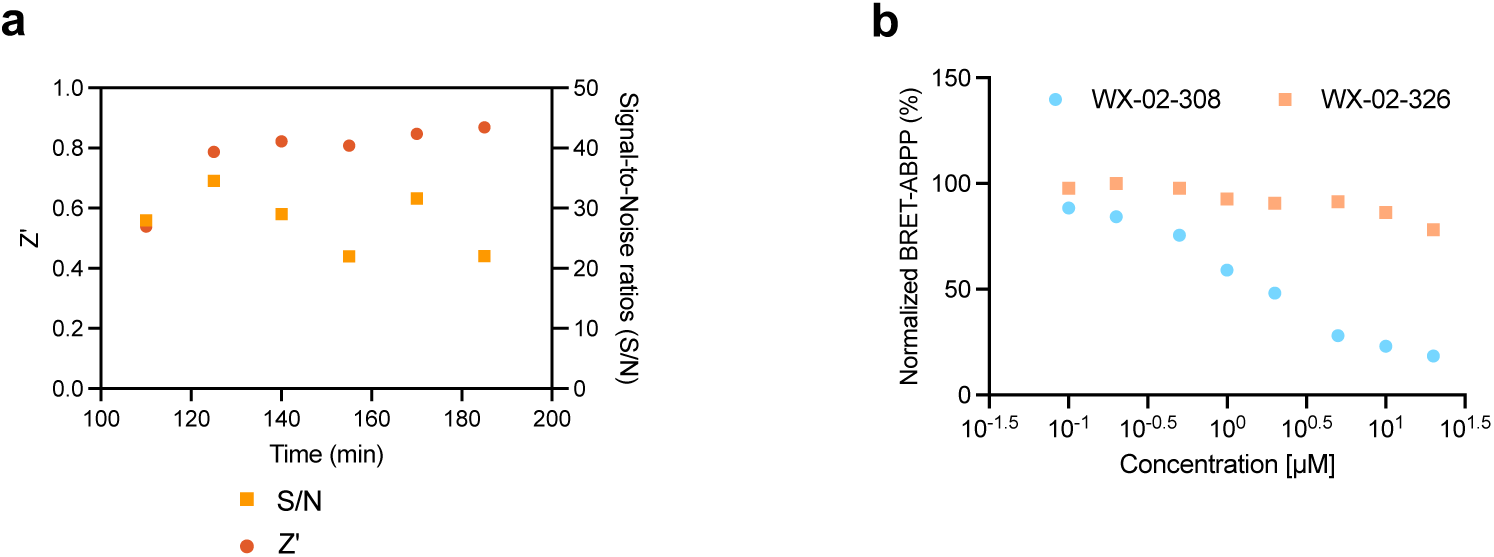
Further characterization of HTS-compatible NanoBRET-ABPP assay for N112C-CCNE1. **a**, Time-dependent assessment of signal-to-noise (S/N, signal: N112C-CCNE1 (YZ-01-A); noise: WT-CCNE1 (YZ-01-A) ) for the NanoBRET assay performed as described in Fig. 5c. **b**, NanoBRET-ABPP data acquired under optimized assay conditions at 185 min time point showing concentration-dependent stereoselective blockade of YZ-01-A engagement of N112C-CCNE1 by WX-02-308 in comparison to WX-02-326 (2 h pre-treatment with compounds) Data are average values ± s.e.m., n = 4.

