## Supplementary Chemistry Information for "Expanding the ligandable proteome by paralog hopping with covalent probes"

### Previously reported small-molecule probes

Stereoisomeric probes and other tool compounds used in this study that were previously reported in the literature are shown below (**Supplementary Figure 1**).[1-4]

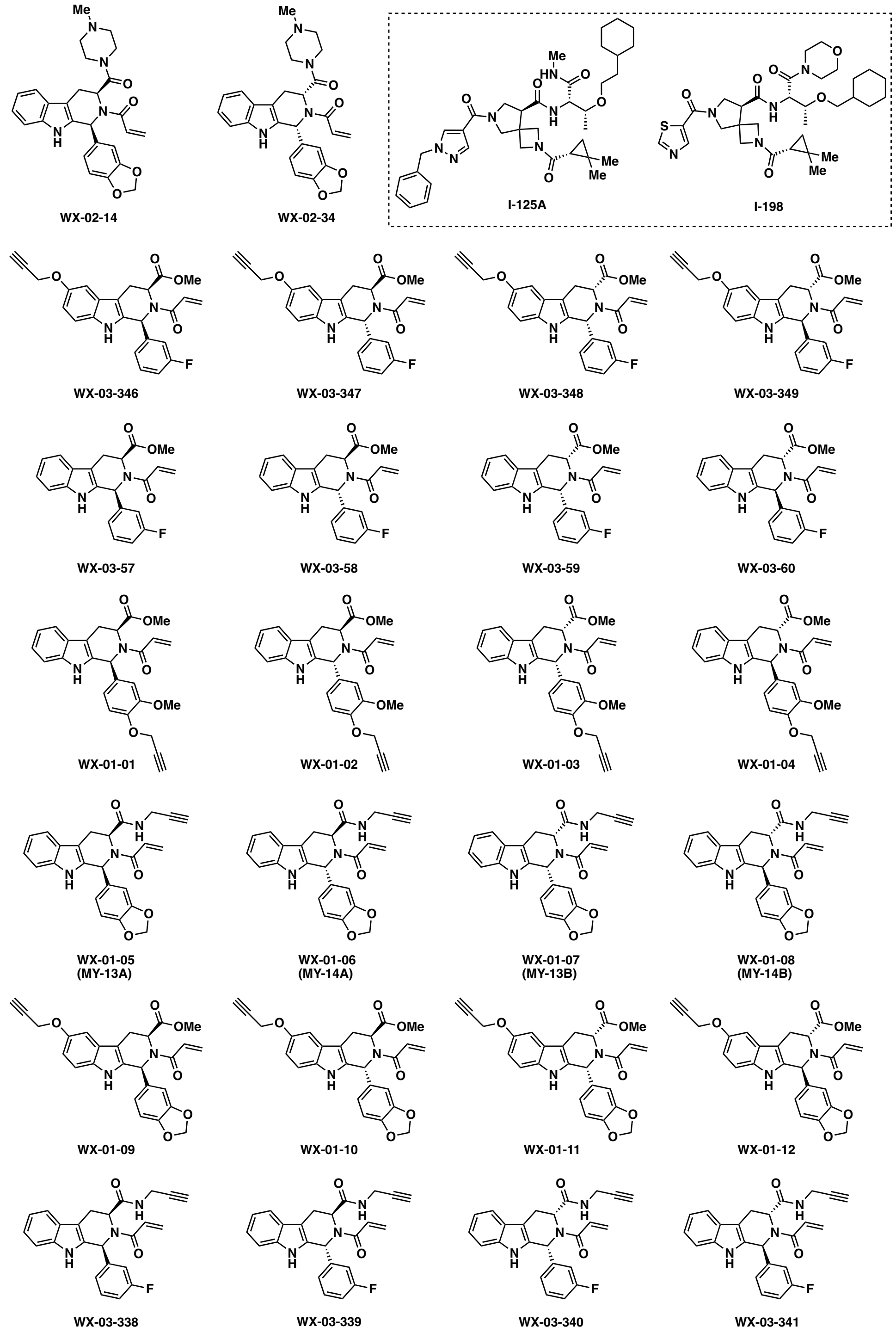


**Supplementary Figure 1. Chemical structures of previously reported stereoisomeric probes and tool compounds used in this study.**

### Synthesis of new stereoisomeric probes

Unless otherwise noted, new small-molecule probes used in this study were prepared by adapting previously reported procedures (**Supplementary Figure 2**, reproduced from Njomen et al.).[1] Analytical characterization data is provided for all new compounds.

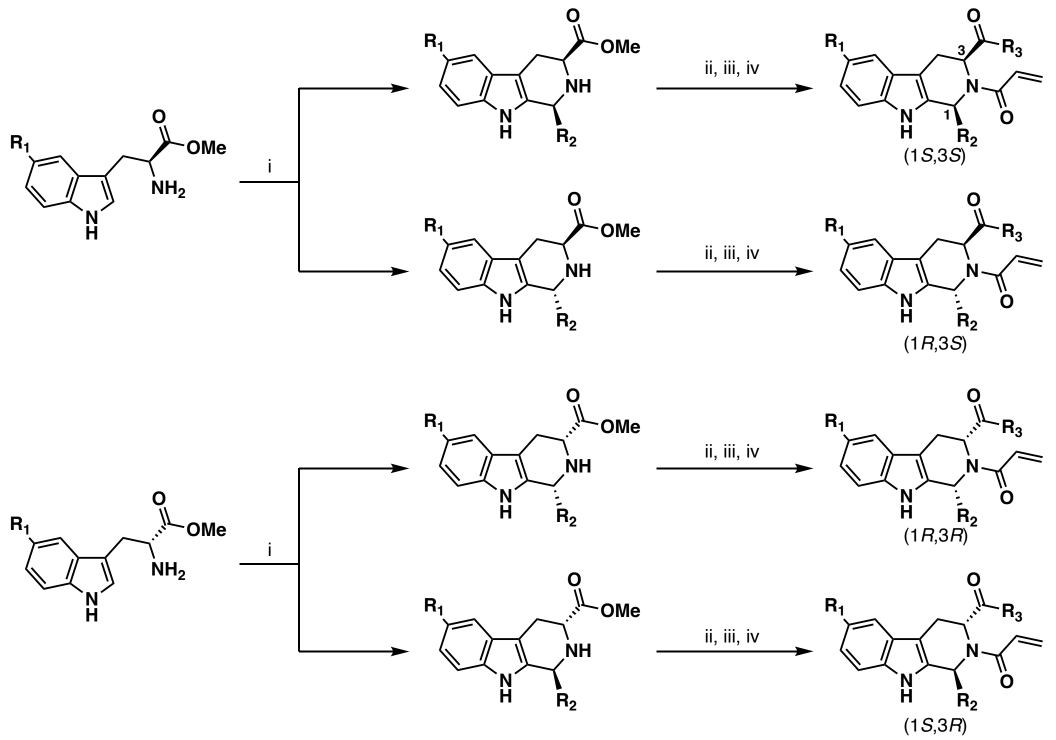


**Supplementary Figure 2. General scheme for the synthesis of tryptoline acrylamide stereoisomeric probes.[1]** Steps: (i) Pictet-Spengler cyclization followed by chromatographic separation of diastereomers; (ii) ester hydrolysis; (iii) amide coupling; (iv) acrylamide installation.

#### General considerations

All NMR spectra were recorded at 298 K unless otherwise noted. $^{1}$H NMR spectra were recorded on Bruker Avance III 400, Avance III HD 400, Avance Neo 400 spectrometers ($^{1}$H, 400 MHz). $^{13}$C NMR spectra were recorded on a Bruker Avance III HD 600 spectrometer ($^{1}$H, 600 MHz; $^{13}$C, 150 MHz). $^{1}$H NMR data are reported as follows: chemical shift (δ), multiplicity (s = singlet, d = doublet, t = triplet, m = multiplet; br = broad), coupling constants, and integration. Chemical shifts are reported in parts per million (ppm) using the appropriate solvent as reference[5]. Analytical supercritical fluid chromatography (SFC) was performed on a Shimadzu LC system (flow rate: 3 mL/min, back pressure: 100 Bar, column temperature: 35 ºC) equipped with a polydiode array detector. High-resolution mass spectra (HRMS) were recorded on an Agilent LC/MSD TOF mass spectrometer.

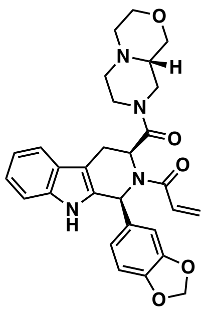

**1-((1*S*,3*S*)-1-(benzo[*d*][1,3]dioxol-5-yl)-3-((*S*)-octahydropyrazino[2,1-*c*][1,4]oxazine-8-carbonyl)-1,3,4,9-tetrahydro-2*H*-pyrido[3,4-*b*]indol-2-yl)prop-2-en-1-one (WX-02-308)**

$^{1}$H NMR (400 MHz, $CD_3OD$) δ 7.56 – 7.47 (m, 1H), 7.33 – 7.15 (m, 2H), 7.14 – 7.07 (m, 1H), 7.06 – 7.00 (m, 1H), 6.89 (br m, 1H), 6.76 (br m, 2H), 6.61 – 6.28 (m, 2H), 6.00 – 5.86 (m, 4H), 4.00 – 3.64 (m, 3H), 3.64 – 3.44 (m, 3H), 3.18 – 2.89 (m, 2H), 2.84 – 2.45 (m, 4H), 2.34 – 2.07 (m, 2H), 2.00 – 1.61 (br m, 1H), 1.48 – 1.21 (br m, 1H). $^{13}$C NMR (150 MHz, $CD_3OD$) δ 170.03, 168.92, 168.63, 149.19, 149.03, 148.84, 138.30, 138.26, 134.57, 133.64, 130.69, 130.40, 129.93, 129.39, 127.74, 127.63, 124.18, 123.26, 122.83, 119.94, 119.15, 112.04, 110.69, 109.96, 109.66, 108.99, 108.82, 102.60, 69.36, 69.24, 67.71, 60.68, 59.90, 56.71, 54.92, 54.74, 54.62, 53.66, 46.73, 45.83, 43.54, 42.82, 24.35, 22.98. HRMS ESI-TOF m/z calculated for $C_{29}H_{31}N_4O_5$ $[M+H]^+$ 515.2289. Found 515.2299.

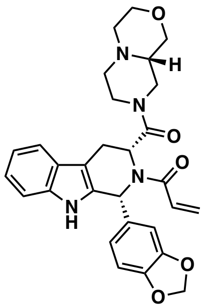

**1-((1*R*,3*R*)-1-(benzo[*d*][1,3]dioxol-5-yl)-3-((*S*)-octahydropyrazino[2,1-*c*][1,4]oxazine-8-carbonyl)-1,3,4,9-tetrahydro-2*H*-pyrido[3,4-*b*]indol-2-yl)prop-2-en-1-one (WX-02-326)**

$^{1}$H NMR (400 MHz, $CD_3OD$) δ 7.57 – 7.47 (m, 1H), 7.33 – 7.15 (m, 2H), 7.14 – 7.07 (m, 1H), 7.06 – 6.99 (m, 1H), 6.86 (br m, 1H), 6.81 – 6.70 (m, 2H), 6.66 – 6.31 (m, 2H), 6.06 – 5.86 (m, 4H), 4.03 – 3.33 (m, 7H), 3.17 – 2.89 (m, 2.5H), 2.64 – 2.46 (m, 2H), 2.33 – 1.53 (m, 4.5H).
HRMS ESI-TOF m/z calculated for $C_{29}H_{31}N_4O_5$ $[M+H]^+$ 515.2289. Found 515.2296.

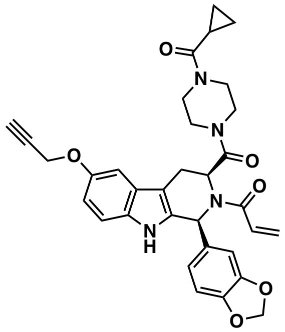

**1-((1*S*,3*S*)-1-(benzo[*d*][1,3]dioxol-5-yl)-3-(4-(cyclopropanecarbonyl)piperazine-1-carbonyl)-6-(prop-2-yn-1-yloxy)-1,3,4,9-tetrahydro-2*H*-pyrido[3,4-*b*]indol-2-yl)prop-2-en-1-one (WX-02-512)**

$^{1}$H NMR (400 MHz, $CD_3OD$) δ 7.30 – 7.08 (m, 3H), 6.93 – 6.80 (m, 3H), 6.75 (br, 1H), 6.62 – 6.29 (m, 2H), 6.04 – 5.73 (m, 4H), 4.73 (d, *J* = 2.4 Hz, 2H), 3.83 – 3.03 (m, 9H), 2.98 (dd, *J* = 15.4, 6.1 Hz, 1H), 2.90 (t, *J* = 2.4 Hz, 1H), 2.82 – 2.41 (m, 1H), 1.86 (s, 1H), 0.97 – 0.70 (m, 4H) .
HRMS ESI-TOF m/z calculated for $C_{33}H_{33}N_4O_6$ $[M+H]^+$ 581.2395. Found 581.2402.

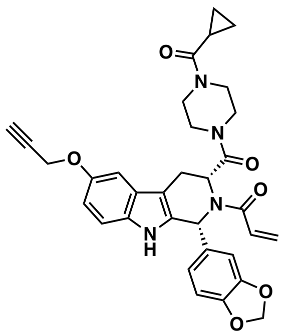

**1-((1*R*,3*R*)-1-(benzo[*d*][1,3]dioxol-5-yl)-3-(4-(cyclopropanecarbonyl)piperazine-1-carbonyl)-6-(prop-2-yn-1-yloxy)-1,3,4,9-tetrahydro-2*H*-pyrido[3,4-*b*]indol-2-yl)prop-2-en-1-one (WX-02-513)**

$^{1}$H NMR (400 MHz, $CD_3OD$) δ 7.30 – 7.07 (m, 3H), 6.95 – 6.80 (m, 3H), 6.73 (br, 1H), 6.64 – 6.29 (m, 2H), 6.02 – 5.73 (m, 4H), 4.74 (d, *J* = 2.4 Hz, 2H), 3.84 – 3.02 (m, 9H), 3.00 (dd, *J* = 15.4, 6.1 Hz, 1H), 2.90 (t, *J* = 2.4 Hz, 1H), 2.83 – 2.41 (m, 1H), 1.86 (s, 1H), 0.98 – 0.70 (m, 4H).
HRMS ESI-TOF m/z calculated for $C_{33}H_{33}N_4O_6$ $[M+H]^+$ 581.2395. Found 581.2404.

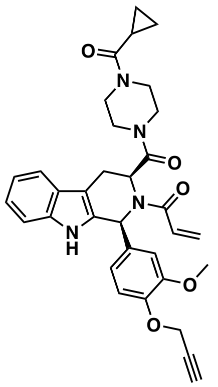

**1-((1*S*,3*S*)-3-(4-(cyclopropanecarbonyl)piperazine-1-carbonyl)-1-(3-methoxy-4-(prop-2-yn-1-yloxy)phenyl)-1,3,4,9-tetrahydro-2*H*-pyrido[3,4-*b*]indol-2-yl)prop-2-en-1-one (WX-02-514)**

$^1$H NMR (400 MHz, $CD_3OD$) δ 7.55 (dt, *J* = 7.6, 1.1 Hz, 1H), 7.40 – 6.81 (m, 7.5H), 6.67 – 6.30 (m, 1.5H), 6.08 – 5.35 (m, 2H), 4.73 (d, *J* = 2.4 Hz, 2H), 3.70 (s, 3H), 3.62 – 2.78 (m, 11H), 3.62 – 2.78 (br d, 1H), 1.84 (br s, 1H), 0.95 – 0.72 (m, 4H).

HRMS ESI-TOF m/z calculated for $C_{33}H_{35}N_4O_5$ $[M+H]^+$ 567.2602. Found 567.2613.

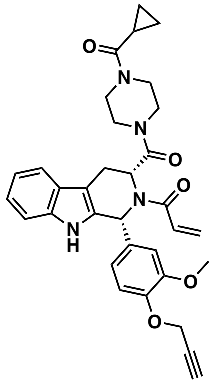

**1-((1*R*,3*R*)-3-(4-(cyclopropanecarbonyl)piperazine-1-carbonyl)-1-(3-methoxy-4-(prop-2-yn-1-yloxy)phenyl)-1,3,4,9-tetrahydro-2*H*-pyrido[3,4-*b*]indol-2-yl)prop-2-en-1-one (WX-02-515)**

$^1$H NMR (400 MHz, $CD_3OD$) δ 7.55 (dt, *J* = 7.6, 1.1 Hz, 1H), 7.42 – 6.84 (m, 7.5H), 6.64 – 6.31 (m, 1.5H), 6.08 – 5.34 (m, 2H), 4.73 (d, *J* = 2.4 Hz, 2H), 3.70 (s, 3H), 3.62 – 2.74 (m, 11H), 3.62 – 2.78 (br d, 1H), 1.84 (br s, 1H), 0.95 – 0.70 (m, 4H).

HRMS ESI-TOF m/z calculated for $C_{33}H_{35}N_4O_5$ $[M+H]^+$ 567.2602. Found 567.2621.

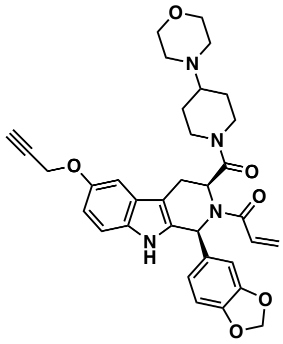

**1-((1*S*,3*S*)-1-(benzo[*d*][1,3]dioxol-5-yl)-6-ethoxy-3-(4-morpholinopiperidine-1-carbonyl)-1,3,4,9-tetrahydro-2*H*-pyrido[3,4-*b*]indol-2-yl)prop-2-en-1-one (WX-02-516)**

$^1$H NMR (400 MHz, $CD_3OD$) δ 7.32 – 7.02 (m, 3H), 6.94 (br s, 1H), 6.89 – 6.66 (m, 3.5H), 6.63– 6.21 (m, 1.5H), 6.18 – 5.17 (m, 4H), 4.73 (t, *J* = 2.5 Hz, 2H), 4.17 – 3.35 (m, 7H), 3.10 – 2.80 (m, 2.5H), 2.78 – 2.00 (m, 7H), 1.98 – 1.48 (m, 2.5H), 1.41 – 0.85 (m, 2H).

HRMS ESI-TOF m/z calculated for $C_{34}H_{37}N_4O_6$ $[M+H]^+$ 597.2708. Found 597.2714.

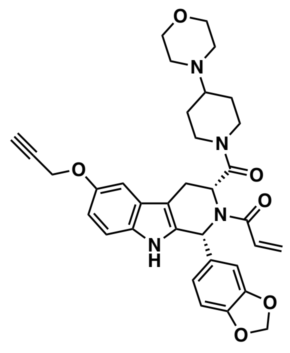

**1-((1*R*,3*R*)-1-(benzo[*d*][1,3]dioxol-5-yl)-6-ethoxy-3-(4-morpholinopiperidine-1-carbonyl)-1,3,4,9-tetrahydro-2*H*-pyrido[3,4-*b*]indol-2-yl)prop-2-en-1-one (WX-02-517)**

$^{1}$H NMR (400 MHz, $CD_3OD$) δ 7.29 – 7.04 (m, 3H), 6.93 (br s, 1H), 6.86 – 6.67 (m, 3.5H), 6.51 – 6.29 (m, 1.5H), 6.14 – 5.26 (m, 4H), 4.73 (t, *J* = 2.5 Hz, 2H), 4.08 – 3.55 (m, 7H), 3.03 – 2.86 (m, 2.5H), 2.70 – 2.06 (m, 7H), 1.93 – 1.47 (m, 2.5H), 1.41 – 1.05 (m, 2H).
HRMS ESI-TOF m/z calculated for $C_{34}H_{37}N_4O_6$ $[M+H]^+$ 597.2708. Found 597.2722.

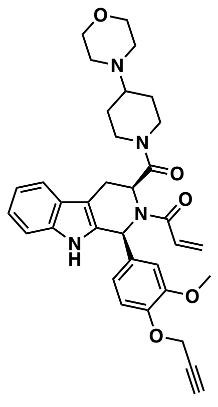

**1-((1*S*,3*S*)-1-(3-methoxy-4-(prop-2-yn-1-yloxy)phenyl)-3-(4-morpholinopiperidine-1-carbonyl)-1,3,4,9-tetrahydro-2*H*-pyrido[3,4-*b*]indol-2-yl)prop-2-en-1-one (WX-02-518)**

$^{1}$H NMR (400 MHz, $CD_3OD$) δ 7.53 (t, *J* = 6.7 Hz, 1H), 7.42 – 6.75 (m, 7.5H), 6.66 – 6.29 (m, 1.5H), 6.08 – 5.34 (m, 2H), 4.77 – 4.68 (m, 2H), 4.03 – 3.54 (m, 9.5H), 3.52 – 3.32 (m, 1H), 3.11 – 2.76 (m, 2.5H), 2.71 – 1.96 (m, 6.5H), 1.88 – 1.24 (m, 3H), 1.23 – 0.32 (m, 1.5H).
HRMS ESI-TOF m/z calculated for $C_{34}H_{39}N_4O_5$ $[M+H]^+$ 583.2915. Found 583.2927.

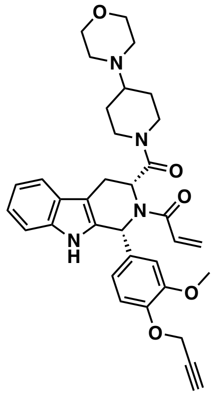

**1-((1*R*,3*R*)-1-(3-methoxy-4-(prop-2-yn-1-yloxy)phenyl)-3-(4-morpholinopiperidine-1-carbonyl)-1,3,4,9-tetrahydro-2*H*-pyrido[3,4-*b*]indol-2-yl)prop-2-en-1-one (WX-02-519)**

$^{1}$H NMR (400 MHz, $CD_3OD$) δ 7.53 (t, *J* = 6.7 Hz, 1H), 7.41 – 6.78 (m, 7.5H), 6.66 – 6.30 (m, 1.5H), 6.02 – 5.30 (m, 2H), 4.77 – 4.67 (m, 2H), 4.04 – 3.51 (m, 9.5H), 3.49 – 3.32 (m, 1H), 3.14 – 2.79 (m, 2.5H), 2.63 – 1.97 (m, 6.5H), 1.91 – 1.22 (m, 3H), 1.20 – 0.18 (m, 1.5H).
HRMS ESI-TOF m/z calculated for $C_{34}H_{39}N_4O_5$ $[M+H]^+$ 583.2915. Found 583.2934.

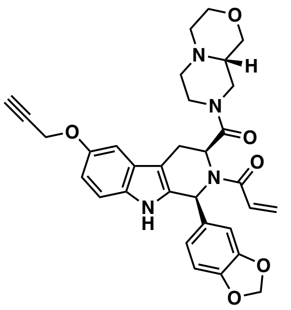

**1-((1*S*,3*S*)-1-(benzo[*d*][1,3]dioxol-5-yl)-6-methoxy-3-((*S*)-octahydropyrazino[2,1-*c*][1,4]oxazine-8-carbonyl)-1,3,4,9-tetrahydro-2*H*-pyrido[3,4-*b*]indol-2-yl)prop-2-en-1-one (WX-02-520)**
$^1$H NMR (400 MHz, $CD_3OD$) δ 7.34 – 7.06 (m, 3H), 6.97 – 6.67 (m, 4.3H), 6.65 – 6.29 (m, 1.7H), 6.11 – 5.27 (m, 4H), 4.73 (t, *J* = 2.4 Hz, 2H), 4.05 – 3.68 (m, 3H), 3.65 – 3.37 (m, 3H), 3.16 – 2.87 (m, 3H), 2.85 – 2.38 (m, 4H), 2.37 – 2.01 (m, 2H), 2.00 – 1.67 (m, 1H), 1.48 – 1.23 (m, 1H). $^{13}$C NMR (150 MHz, $CD_3OD$) δ 170.94, 168.93, 168.63, 153.27, 149.19, 149.03, 148.83, 134.59, 133.98, 133.93, 133.65, 133.06, 131.86, 131.45, 130.63, 130.17, 129.95, 129.39, 127.96, 127.83, 124.83, 124.14, 123.24, 113.59, 112.66, 110.70, 109.88, 109.58, 108.99, 108.82, 103.72, 102.61, 80.63, 76.18, 69.37, 69.24, 67.71, 60.69, 59.90, 57.85, 56.73, 54.92, 54.75, 54.63, 53.67, 46.74, 45.85, 43.54, 42.83, 30.76, 30.33, 26.92, 23.74, 23.01.
HRMS ESI-TOF m/z calculated for $C_{32}H_{33}N_4O_6$ $[M+H]^+$ 569.2395. Found 569.2416.

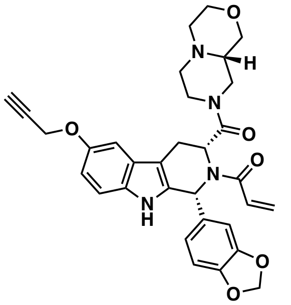

**1-((1*R*,3*R*)-1-(benzo[*d*][1,3]dioxol-5-yl)-6-methoxy-3-((*S*)-octahydropyrazino[2,1-*c*][1,4]oxazine-8-carbonyl)-1,3,4,9-tetrahydro-2*H*-pyrido[3,4-*b*]indol-2-yl)prop-2-en-1-one (WX-02-521)**
$^1$H NMR (400 MHz, $CD_3OD$) δ 7.32 – 7.08 (m, 3H), 6.99 – 6.60 (m, 4.3H), 6.59 – 6.19 (m, 1.7H), 6.11 – 5.24 (m, 4H), 4.73 (t, *J* = 2.4 Hz, 2H), 4.07 – 3.45 (m, 5H), 3.43 – 3.32 (m, 1.5H), 3.17 – 2.87 (m, 3.5H), 2.70 – 2.43 (m, 2H), 2.36 – 1.52 (m, 4H), 1.45 – 1.01 (m, 1H).
HRMS ESI-TOF m/z calculated for $C_{32}H_{33}N_4O_6$ $[M+H]^+$ 569.2395. Found 569.2409.

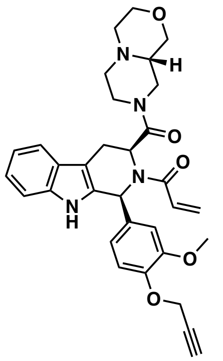

**1-((1*S*,3*S*)-1-(3-methoxy-4-(prop-2-yn-1-yloxy)phenyl)-3-((*S*)-octahydropyrazino[2,1-*c*][1,4]oxazine-8-carbonyl)-1,3,4,9-tetrahydro-2*H*-pyrido[3,4-*b*]indol-2-yl)prop-2-en-1-one (WX-02-522)**
$^1$H NMR (400 MHz, $CD_3OD$) δ 7.54 (dd, *J* = 7.9, 3.4 Hz, 1H), 7.29 (br t, 2H), 7.16 – 6.95 (m, 4H), 6.88 (br s, 1.5H), 6.66 – 6.32 (m, 1.5H), 6.09 – 5.24 (m, 2H), 4.74 (d, *J* = 2.4 Hz, 2H), 4.00 – 3.26 (m, 10H), 3.18 – 2.86 (m, 3H), 2.83 – 2.33 (m, 3.5H), 2.31 – 1.82 (m, 2H), 1.73 – 0.94 (m, 1.5H).
HRMS ESI-TOF m/z calculated for $C_{32}H_{35}N_4O_5$ $[M+H]^+$ 555.2602. Found 555.2617.

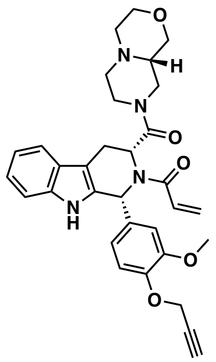

**1-((1*R*,3*R*)-1-(3-methoxy-4-(prop-2-yn-1-yloxy)phenyl)-3-((*S*)-octahydropyrazino[2,1-*c*][1,4]oxazine-8-carbonyl)-1,3,4,9-tetrahydro-2*H*-pyrido[3,4-*b*]indol-2-yl)prop-2-en-1-one (WX-02-523)**

$^1$H NMR (400 MHz, $CD_3OD$) δ 7.54 (t, *J* = 6.8 Hz, 1H), 7.45 – 6.72 (m, 7.5H), 6.66 – 6.27 (m, 1.5H), 6.04 – 5.35 (m, 2H), 4.74 (t, *J* = 2.9 Hz, 2H), 4.07 – 3.18 (m, 10H), 3.17 – 2.80 (m, 3H), 2.67 – 2.38 (m, 2H), 2.30 – 1.11 (m, 5H).

HRMS ESI-TOF m/z calculated for $C_{32}H_{35}N_4O_5$ $[M+H]^+$ 555.2602. Found 555.2615.

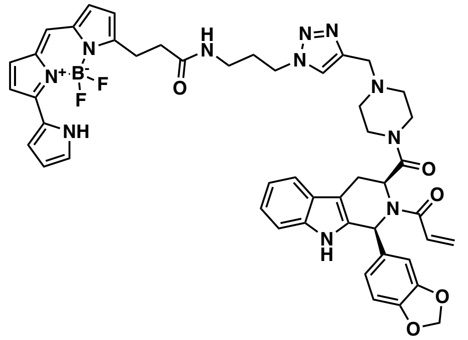

**N-(3-(4-((4-((1*S*,3*S*)-2-acryloyl-1-(benzo[*d*][1,3]dioxol-5-yl)-2,3,4,9-tetrahydro-1*H*-pyrido[3,4-*b*]indole-3-carbonyl)piperazin-1-yl)methyl)-1*H*-1,2,3-triazol-1-yl)propyl)-3-(5,5-difluoro-7-(1*H*-pyrrol-2-yl)-5*H*-5λ4,6λ4-dipyrrolo[1,2-*c*:2',1'-*f*][1,3,2]diazaborinin-3-yl)propanamide (YZ-01-A)**

$^1$H NMR (600 MHz, $CDCl_3$) δ 10.40 (s, 1H), 7.68 (s, 1H), 7.59 (d, *J* = 7.8 Hz, 1H), 7.45 (s, 1H), 7.29 – 7.23 (m, 3H), 7.21 – 7.15 (m, 2H), 7.13 (t, *J* = 7.4 Hz, 1H), 7.06 (d, *J* = 4.6 Hz, 1H), 7.02 (br s, 1H), 6.98 (s, 1H), 6.95 – 6.74 (m, 5H), 6.69 (d, *J* = 8.1 Hz, 1H), 6.46 – 6.36 (m, 2H), 6.30 (d, *J* = 3.9 Hz, 1H), 5.99 – 5.89 (m, 3H), 5.86 (dd, *J* = 10.5, 1.6 Hz, 1H), 4.23 (t, *J* = 6.7 Hz, 2H), 3.58 – 3.27 (m, 7H), 3.21 (q, *J* = 6.3 Hz, 2H), 3.09 – 2.89 (m, 2H), 2.68 (t, *J* = 7.4 Hz, 2H), 2.57 (br s, 1H), 2.37 – 2.06 (m, 4H), 2.01 (m, 2H).

HRMS ESI-TOF m/z calculated for $C_{48}H_{48}BF_2N_{11}O_5$ $[M]^+$ 907.4016. Found 907.4008.

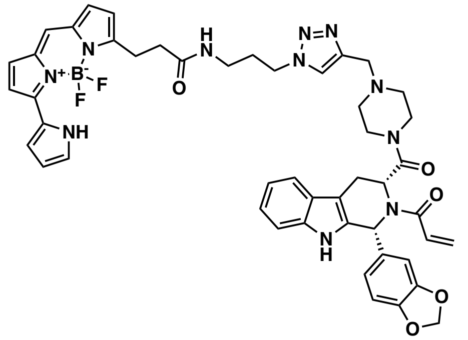

**N-(3-(4-((4-((1*R*,3*R*)-2-acryloyl-1-(benzo[*d*][1,3]dioxol-5-yl)-2,3,4,9-tetrahydro-1*H*-pyrido[3,4-*b*]indole-3-carbonyl)piperazin-1-yl)methyl)-1*H*-1,2,3-triazol-1-yl)propyl)-3-(5,5-difluoro-7-(1*H*-pyrrol-2-yl)-5*H*-5⎣4,6⎣4-dipyrrolo[1,2-*c*:2',1'-*f*][1,3,2]diazaborinin-3-yl)propanamide (YZ-01-B)**

$^{1}H$ NMR (600 MHz, $CDCl_3$) δ 10.40 (s, 1H), 7.74 (s, 1H), 7.58 (d, *J* = 7.8 Hz, 1H), 7.46 (s, 1H), 7.29 – 7.23 (m, 3H), 7.21 – 7.15 (m, 2H), 7.13 (t, *J* = 7.4 Hz, 1H), 7.06 (d, *J* = 4.6 Hz, 1H), 7.01 (br s, 1H), 6.98 (s, 1H), 6.94 – 6.74 (m, 5H), 6.69 (d, *J* = 8.1 Hz, 1H), 6.45 – 6.36 (m, 2H), 6.30 (d, *J* = 3.9 Hz, 1H), 6.03 – 5.89 (m, 3H), 5.85 (dd, *J* = 10.5, 1.6 Hz, 1H), 4.22 (t, *J* = 6.7 Hz, 2H), 3.58 – 3.26 (m, 7H), 3.20 (q, *J* = 6.3 Hz, 2H), 3.09 – 2.91 (m, 2H), 2.67 (t, *J* = 7.5 Hz, 2H), 2.57 (br s, 1H), 2.38 – 2.07 (m, 4H), 2.01 (m, 2H). $^{13}C$ NMR (150 MHz, $CDCl_3$) δ 172.17, 154.42, 154.23, 150.77, 147.77, 147.54, 137.46, 136.42, 133.35, 131.89, 130.17, 128.73, 127.82, 126.55, 126.12, 123.54, 123.27, 122.50, 120.62, 119.79, 118.76, 118.19, 116.67, 111.68, 110.84, 109.13, 108.00, 101.29, 66.81, 66.75, 52.80, 52.28, 51.94, 47.40, 44.49, 41.57, 40.95, 36.27, 35.79, 30.11, 24.64.
HRMS ESI-TOF m/z calculated for $C_{48}H_{48}BF_2N_{11}O_5$ $[M]^+$ 907.4016. Found 907.4014.

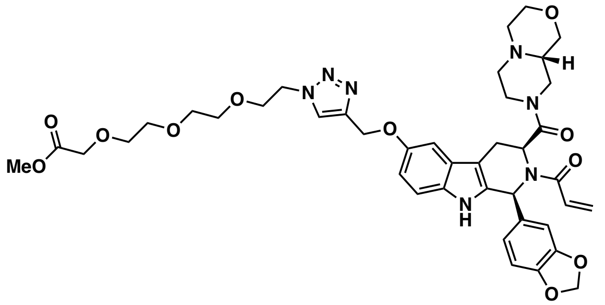

**methyl 2-(2-(2-(2-(4-((((1*S*,3*S*)-2-acryloyl-1-(benzo[*d*][1,3]dioxol-5-yl)-3-((*S*)-octahydropyrazino[2,1-*c*][1,4]oxazine-8-carbonyl)-2,3,4,9-tetrahydro-1*H*-pyrido[3,4-*b*]indol-6-yl)oxy)methyl)-1*H*-1,2,3-triazol-1-yl)ethoxy)ethoxy)ethoxy)acetate (WX-02-520-conjugate)**
HRMS ESI-TOF m/z calculated for $C_{41}H_{51}N_7O_{11}$ $[M+H]^+$ 816.3563. Found 816.3571.

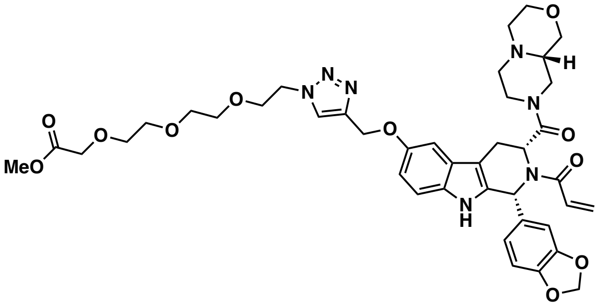

**methyl 2-(2-(2-(2-(4-((((1*R*,3*R*)-2-acryloyl-1-(benzo[*d*][1,3]dioxol-5-yl)-3-((*S*)-octahydropyrazino[2,1-*c*][1,4]oxazine-8-carbonyl)-2,3,4,9-tetrahydro-1*H*-pyrido[3,4-*b*]indol-6-yl)oxy)methyl)-1*H*-1,2,3-triazol-1-yl)ethoxy)ethoxy)ethoxy)acetate (WX-02-521-conjugate)**
HRMS ESI-TOF m/z calculated for $C_{41}H_{51}N_7O_{11}$ $[M+H]^+$ 816.3563. Found 816.3568.

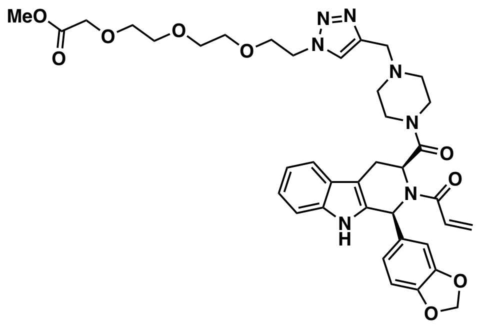

**methyl 2-(2-(2-(2-(4-((4-((1*S*,3*S*)-2-acryloyl-1-(benzo[*d*][1,3]dioxol-5-yl)-2,3,4,9-tetrahydro-1*H*-pyrido[3,4-*b*]indole-3-carbonyl)piperazin-1-yl)methyl)-1*H*-1,2,3-triazol-1-yl)ethoxy)ethoxy)ethoxy)acetate (WX-02-588-conjugate)**
HRMS ESI-TOF m/z calculated for $C_{38}H_{46}N_7O_9$ $[M+H]^+$ 744.3352. Found 744.3363.

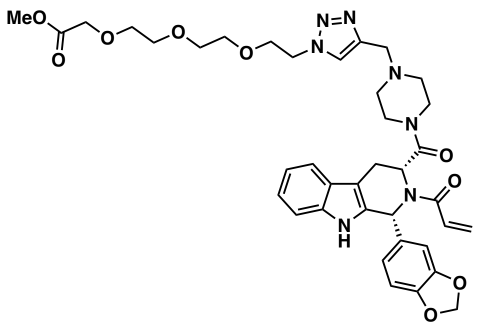

**methyl 2-(2-(2-(2-(4-((4-((1*R*,3*R*)-2-acryloyl-1-(benzo[*d*][1,3]dioxol-5-yl)-2,3,4,9-tetrahydro-1*H*-pyrido[3,4-*b*]indole-3-carbonyl)piperazin-1-yl)methyl)-1*H*-1,2,3-triazol-1-yl)ethoxy)ethoxy)ethoxy)acetate (WX-02-589-conjugate)**
HRMS ESI-TOF m/z calculated for $C_{38}H_{46}N_7O_9$ $[M+H]^+$ 744.3352. Found 744.3358.

### Chiral-phase SFC data

## WX-02-512

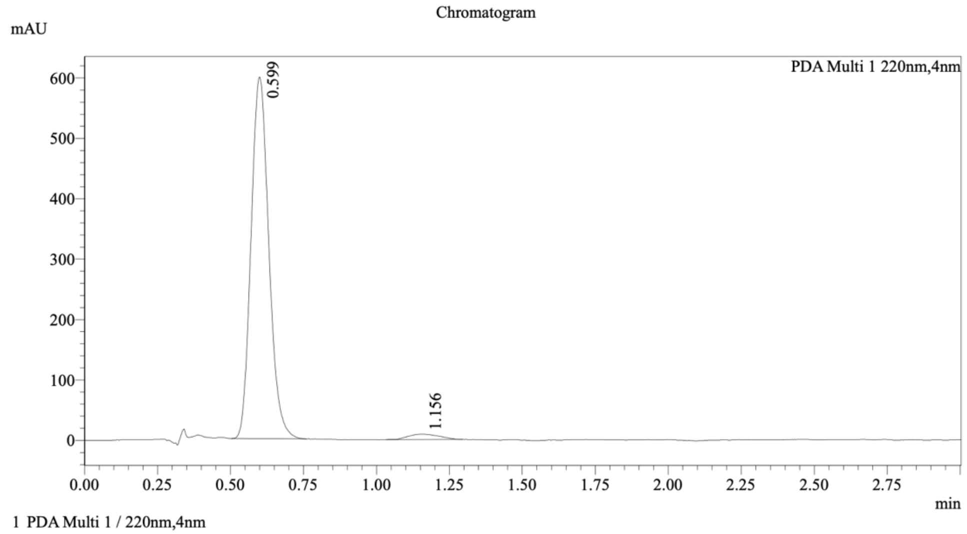


Integration Result

Peak Table

PDA Ch1 220nm

| Peak# | Ret. Time | Height | Height% | Resolution(USP) | Area | Area% |
|---|---|---|---|---|---|---|
| 1 | 0.599 | 595470 | 98.546 | -- | 2542970 | 97.516 |
| 2 | 1.156 | 8787 | 1.454 | 3.535 | 64786 | 2.484 |

## WX-02-513

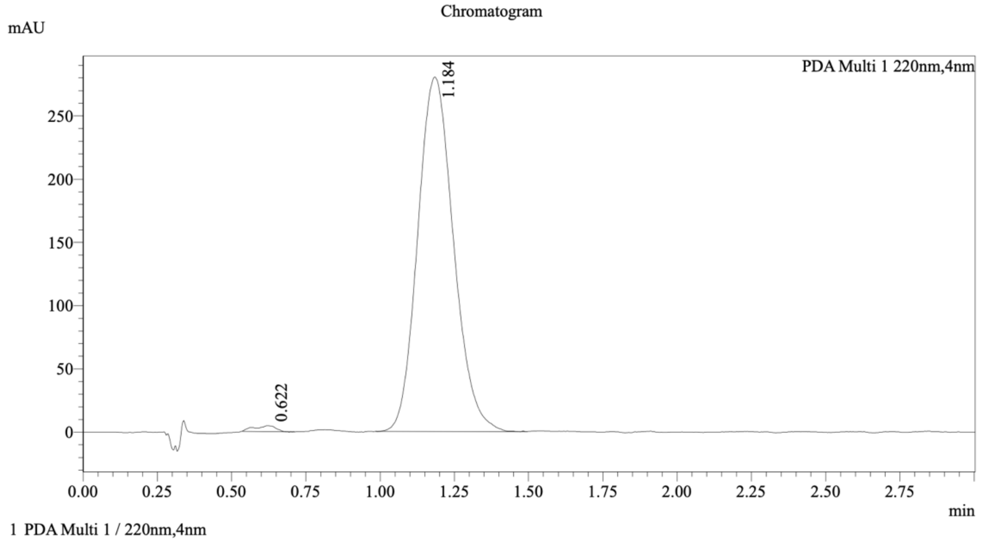


Integration Result

Peak Table

PDA Ch1 220nm

| Peak# | Ret. Time | Height | Height% | Resolution(USP) | Area | Area% |
|---|---|---|---|---|---|---|
| 1 | 0.622 | 4727 | 1.662 | -- | 23164 | 0.975 |
| 2 | 1.184 | 279728 | 98.338 | 3.175 | 2352238 | 99.025 |

**Co-injection of WX-02-512 and WX-02-513**

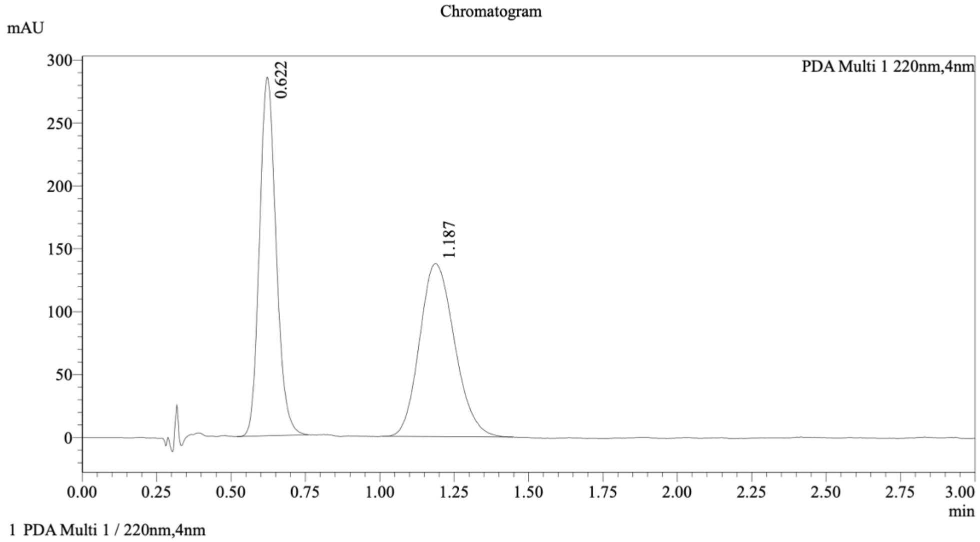


Integration Result

Peak Table

PDA Ch1 220nm

| Peak# | Ret. Time | Height | Height% | Resolution(USP) | Area | Area% |
|---|---|---|---|---|---|---|
| 1 | 0.622 | 283966 | 67.404 | -- | 1109455 | 49.607 |
| 2 | 1.187 | 137323 | 32.596 | 3.545 | 1127052 | 50.393 |

**Method details**

Column: Chiralpak AD-3, 50 mm length × 4.6 mm internal diameter, 3 µm particle size.
Mobile phase: A: supercritical $CO_2$; B: *i*-PrOH/$CH_3CN$/$Et_2NH$ (66.3:33.2:0.05).
Gradient elution: 50% B.

**WX-02-514**

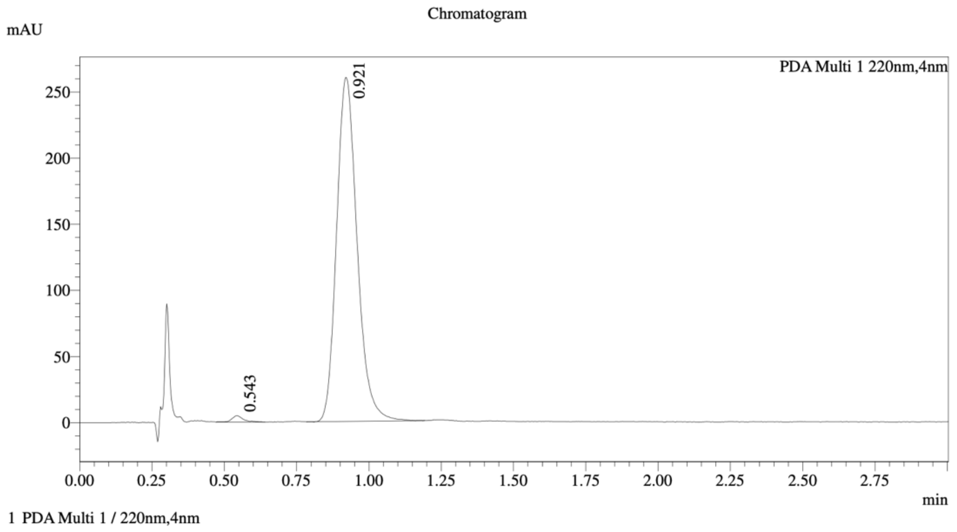


Integration Result

Peak Table

PDA Ch1 220nm

| Peak# | Ret. Time | Height | Height% | Resolution(USP) | Area | Area% |
|---|---|---|---|---|---|---|
| 1 | 0.543 | 4716 | 1.795 | -- | 12835 | 0.989 |
| 2 | 0.921 | 257966 | 98.205 | 3.831 | 1284315 | 99.011 |

**WX-02-515**

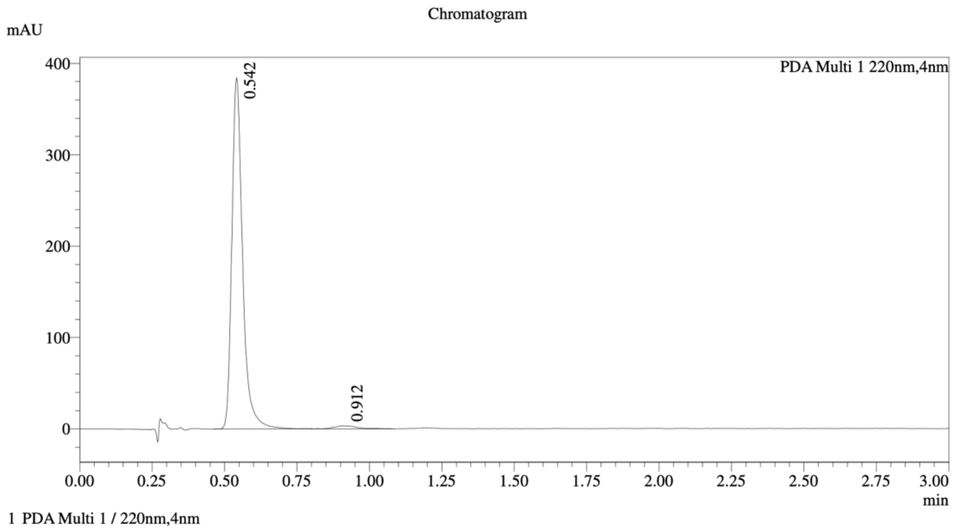


Integration Result

Peak Table

PDA Ch1 220nm

| Peak# | Ret. Time | Height | Height% | Resolution(USP) | Area | Area% |
|---|---|---|---|---|---|---|
| 1 | 0.542 | 380714 | 99.123 | -- | 993467 | 98.059 |
| 2 | 0.912 | 3369 | 0.877 | 3.533 | 19664 | 1.941 |

**Co-injection of WX-02-514 and WX-02-515**

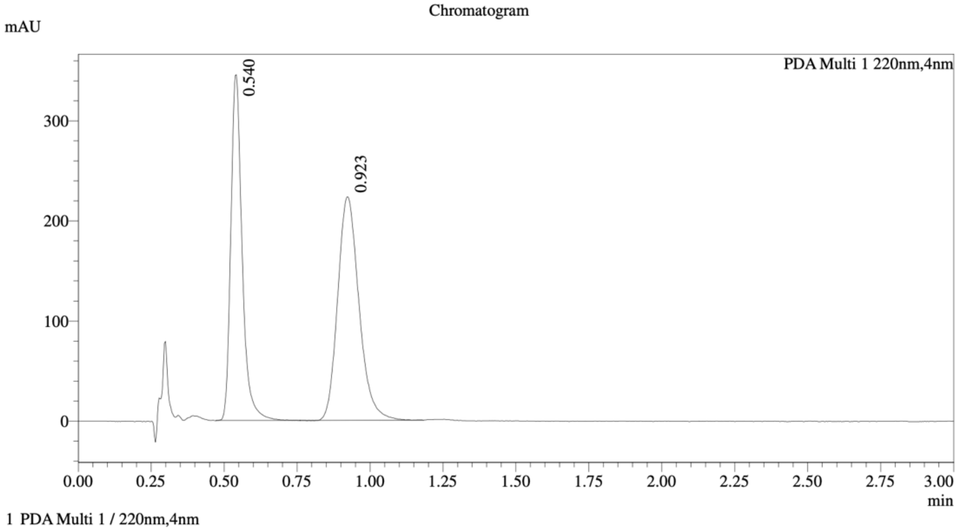


==========================================================================
Integration Result
==========================================================================

Peak Table

PDA Ch1 220nm

| Peak# | Ret. Time | Height | Height% | Resolution(USP) | Area | Area% |
|---|---|---|---|---|---|---|
| 1 | 0.540 | 342308 | 60.662 | -- | 930779 | 45.123 |
| 2 | 0.923 | 221979 | 39.338 | 3.754 | 1131960 | 54.877 |

**Method details**

Column: Chiralpak IC-3, 50 mm length × 4.6 mm internal diameter, 3 μm particle size.
Mobile phase: A: supercritical $CO_2$; B: MeOH/$CH_3CN$/$Et_2NH$ (66.3:33.2:0.05).
Gradient elution: 50% B.

**WX-02-516**

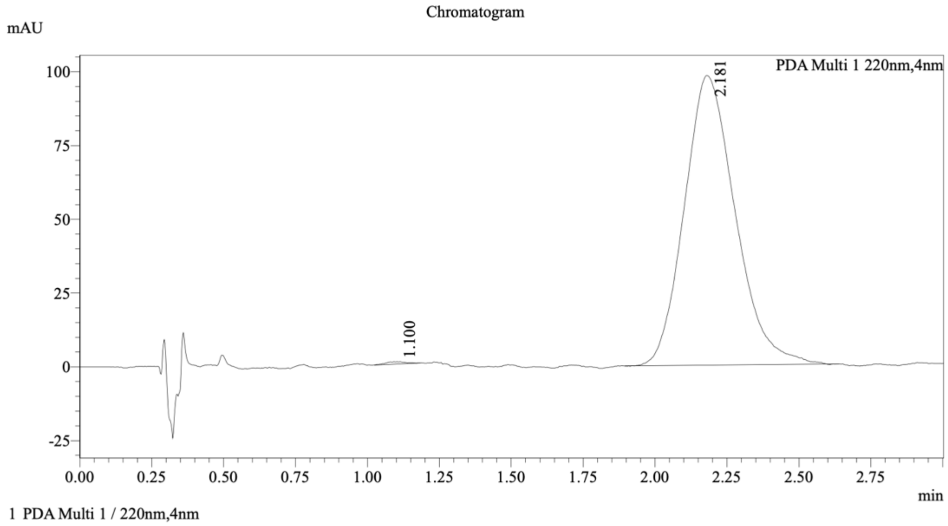


Integration Result

Peak Table

PDA Ch1 220nm

| Peak# | Ret. Time | Height | Height% | Resolution(USP) | Area | Area% |
|---|---|---|---|---|---|---|
| 1 | 1.100 | 751 | 0.760 | -- | 3644 | 0.300 |
| 2 | 2.181 | 98141 | 99.240 | 4.963 | 1209729 | 99.700 |

**WX-02-517**

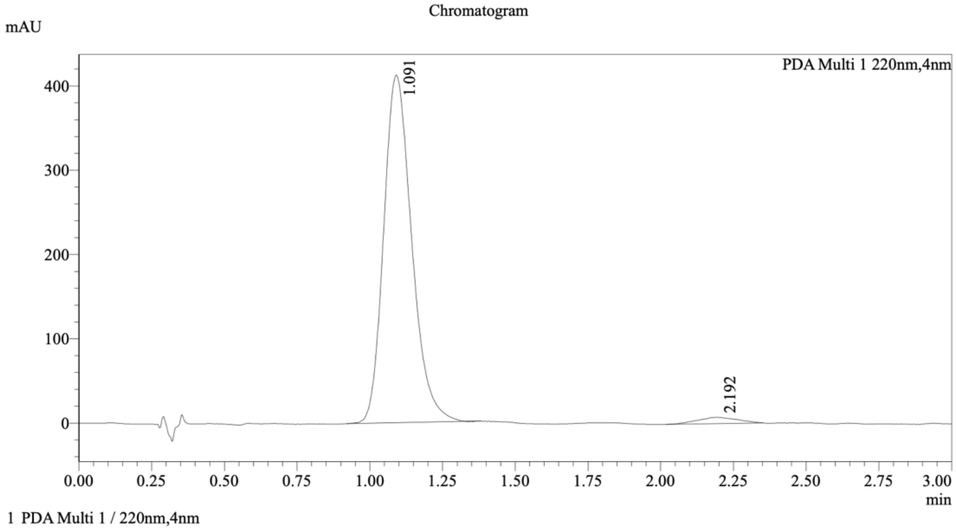


Integration Result

Peak Table

PDA Ch1 220nm

| Peak# | Ret. Time | Height | Height% | Resolution(USP) | Area | Area% |
|---|---|---|---|---|---|---|
| 1 | 1.091 | 411267 | 98.216 | -- | 2766502 | 97.474 |
| 2 | 2.192 | 7469 | 1.784 | 4.749 | 71700 | 2.526 |

### Co-injection of WX-02-516 and WX-02-517

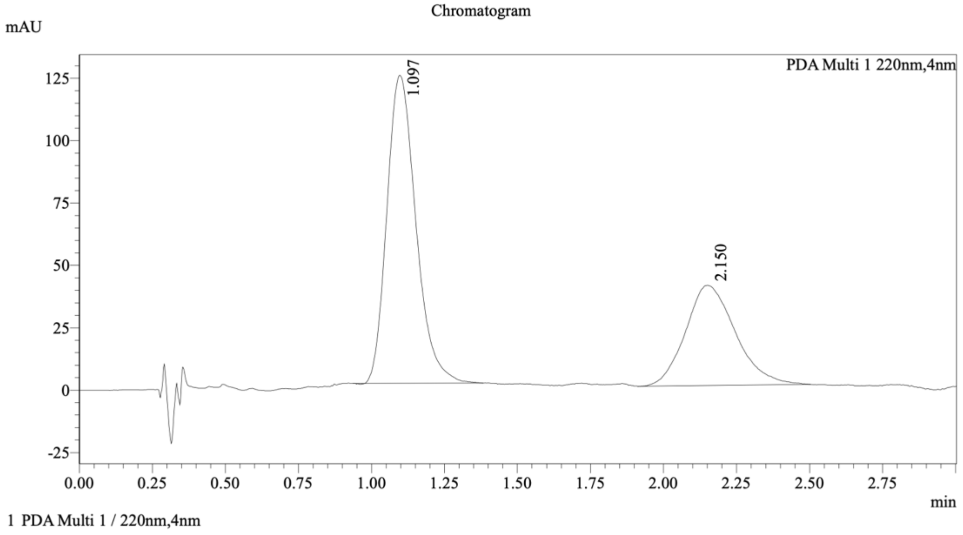


Integration Result

Peak Table

PDA Ch1 220nm

| Peak# | Ret. Time | Height | Height% | Resolution(USP) | Area | Area% |
|---|---|---|---|---|---|---|
| 1 | 1.097 | 122903 | 75.385 | -- | 854163 | 64.165 |
| 2 | 2.150 | 40131 | 24.615 | 4.320 | 477033 | 35.835 |

#### Method details

Column: Chiralpak AD-3, 50 mm length × 4.6 mm internal diameter, 3 µm particle size.
Mobile phase: A: supercritical $CO_2$; B: *i*-PrOH/$CH_3CN$/$Et_2NH$ (66.3:33.2:0.05).
Gradient elution: 40% B.

**WX-02-518**

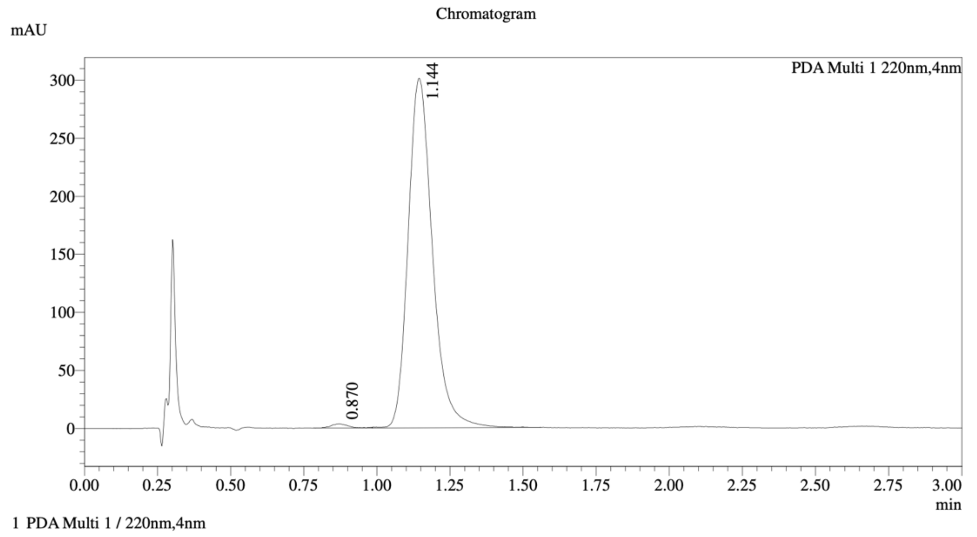


Integration Result

Peak Table

PDA Ch1 220nm

| Peak# | Ret. Time | Height | Height% | Resolution(USP) | Area | Area% |
|---|---|---|---|---|---|---|
| 1 | 0.870 | 3550 | 1.171 | -- | 14419 | 0.822 |
| 2 | 1.144 | 299476 | 98.829 | 2.098 | 1739745 | 99.178 |

**WX-02-519**

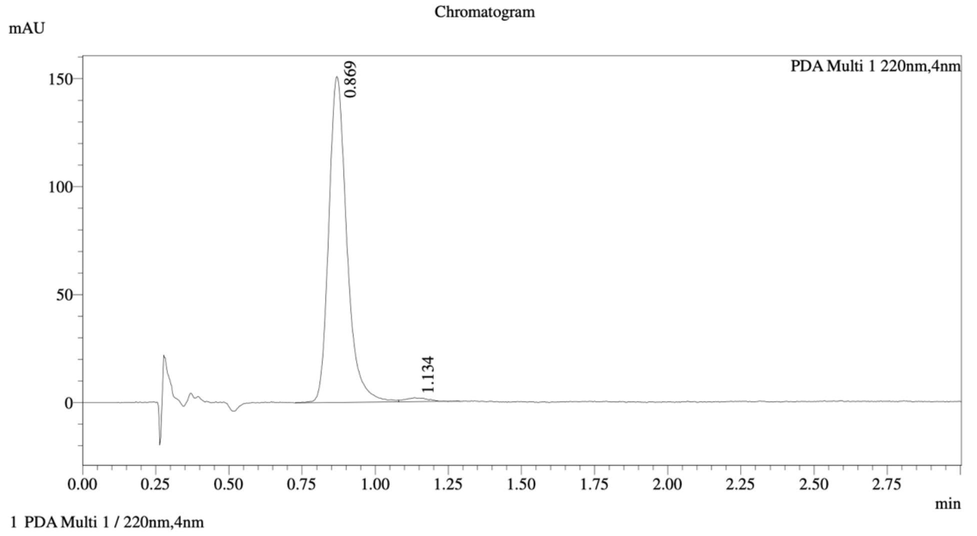


Integration Result

Peak Table

PDA Ch1 220nm

| Peak# | Ret. Time | Height | Height% | Resolution(USP) | Area | Area% |
|---|---|---|---|---|---|---|
| 1 | 0.869 | 150427 | 98.890 | -- | 648058 | 98.645 |
| 2 | 1.134 | 1689 | 1.110 | 2.033 | 8899 | 1.355 |

### Co-injection of WX-02-518 and WX-02-519

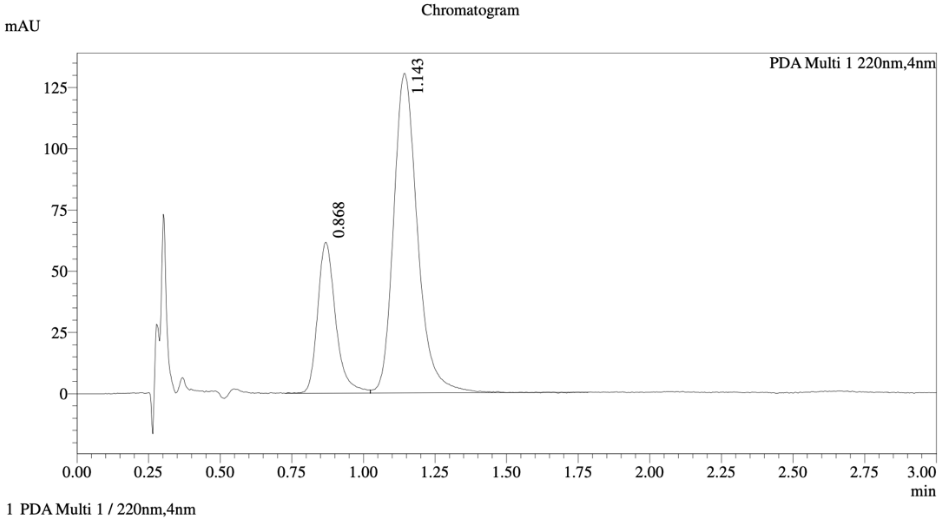


Integration Result

Peak Table

PDA Ch1 220nm

| Peak# | Ret. Time | Height | Height% | Resolution(USP) | Area | Area% |
|---|---|---|---|---|---|---|
| 1 | 0.868 | 61427 | 32.061 | -- | 270231 | 26.272 |
| 2 | 1.143 | 130167 | 67.939 | 2.105 | 758375 | 73.728 |

**Method details**

Column: Chiralpak IC-3, 50 mm length × 4.6 mm internal diameter, 3 μm particle size.
Mobile phase: A: supercritical $CO_2$; B: MeOH/$CH_3CN$/$Et_2NH$ (66.3:33.2:0.05).
Gradient elution: 40% B.

**Spectroscopic data**

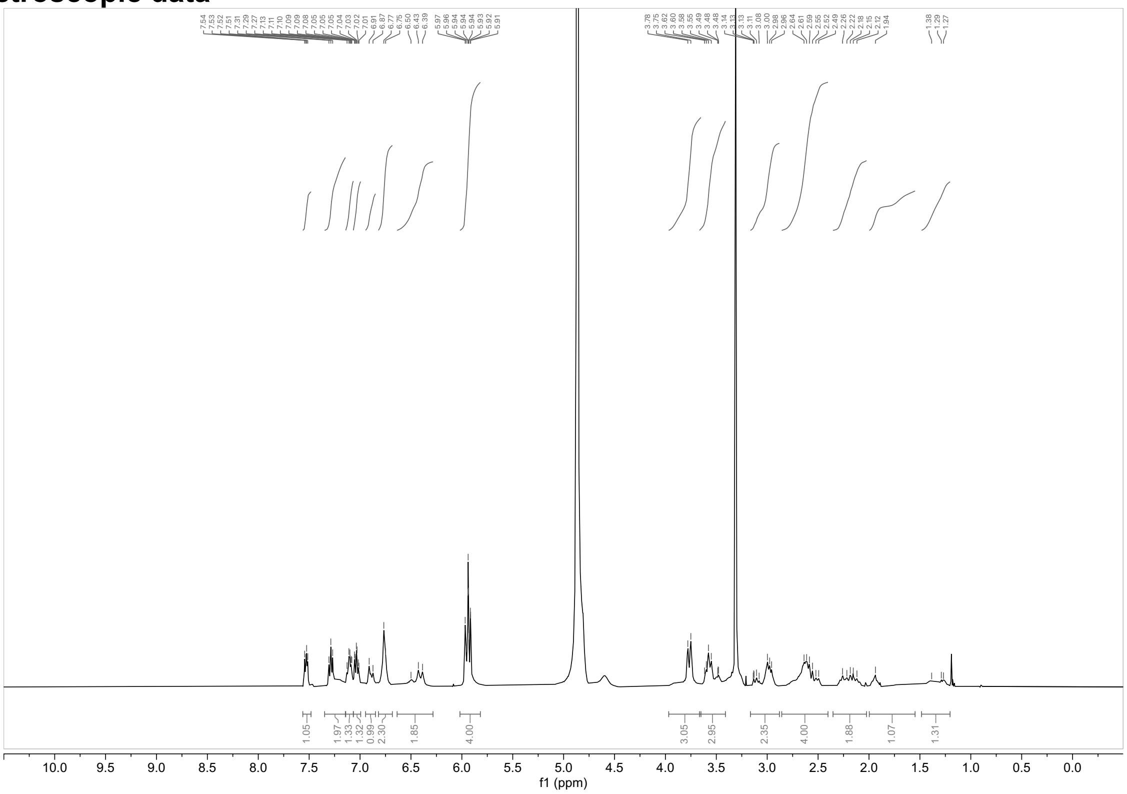

**$^{1}H$ NMR spectrum of WX-02-308 ($CD_3OD$, 400 MHz)**

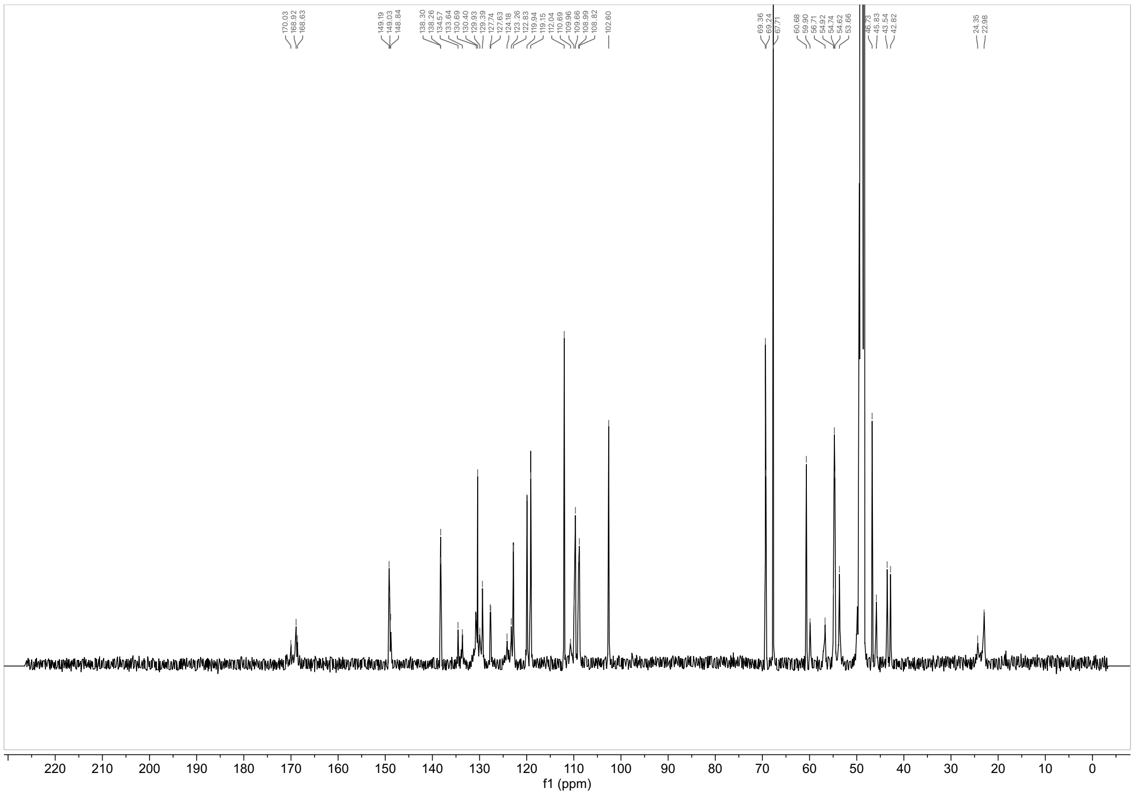

**$^{13}C$ NMR spectrum of WX-02-308 ($CD_3OD$, 150 MHz)**

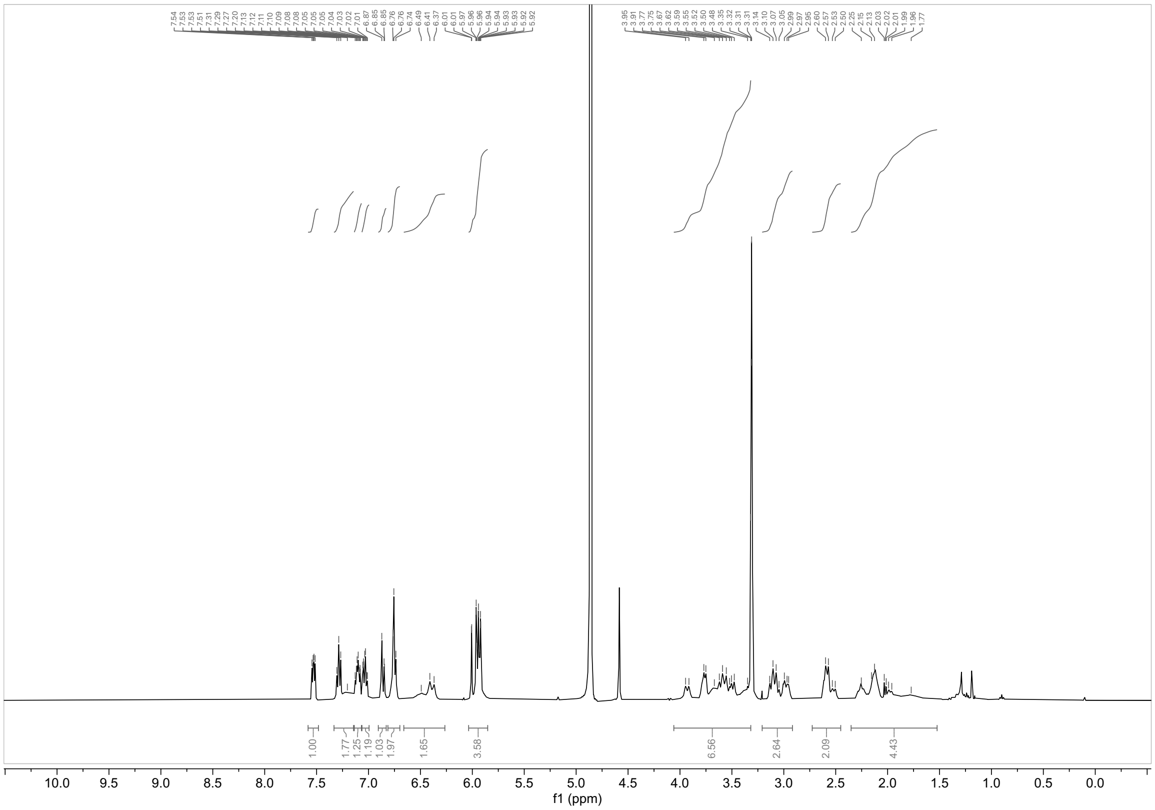

**$^1$H NMR spectrum of WX-02-326 ($CD_3OD$, 400 MHz)**

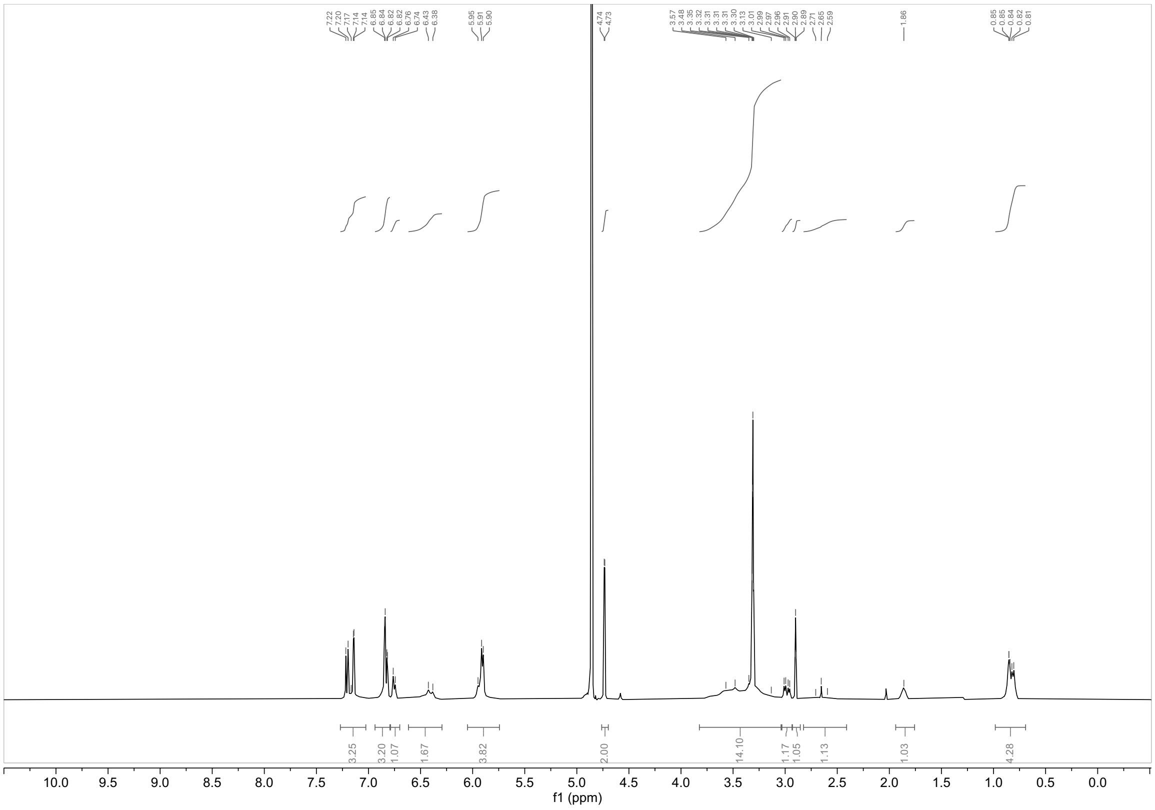

**$^1$H NMR spectrum of WX-02-512 ($CD_3OD$, 400 MHz)**

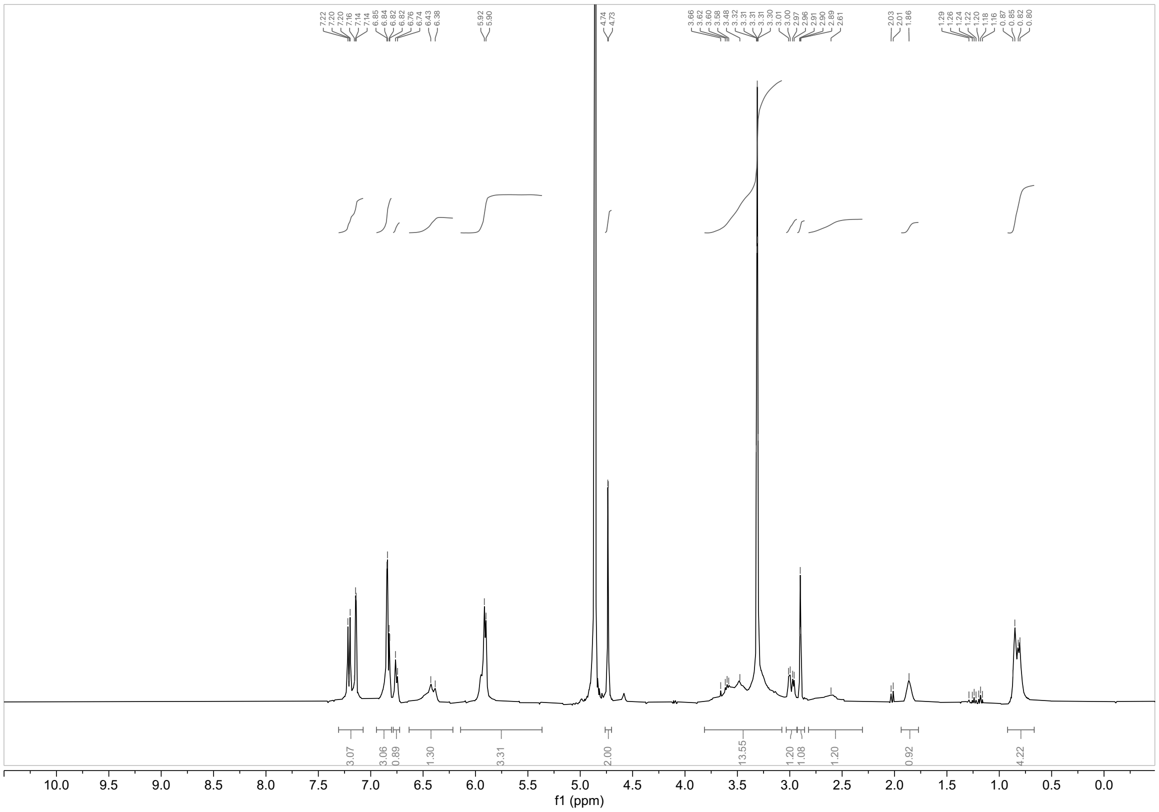

**$^1$H NMR spectrum of WX-02-513 ($CD_3OD$, 400 MHz)**

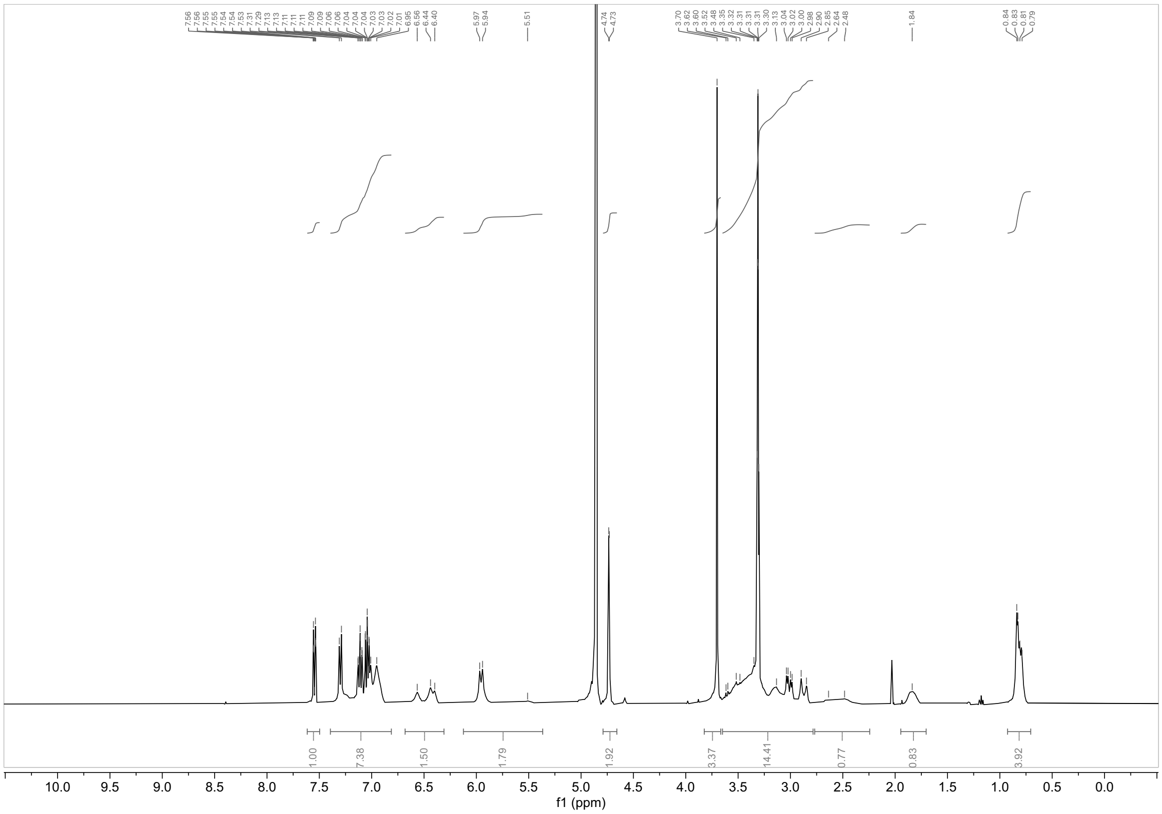

**$^1$H NMR spectrum of WX-02-514 ($CD_3OD$, 400 MHz)**

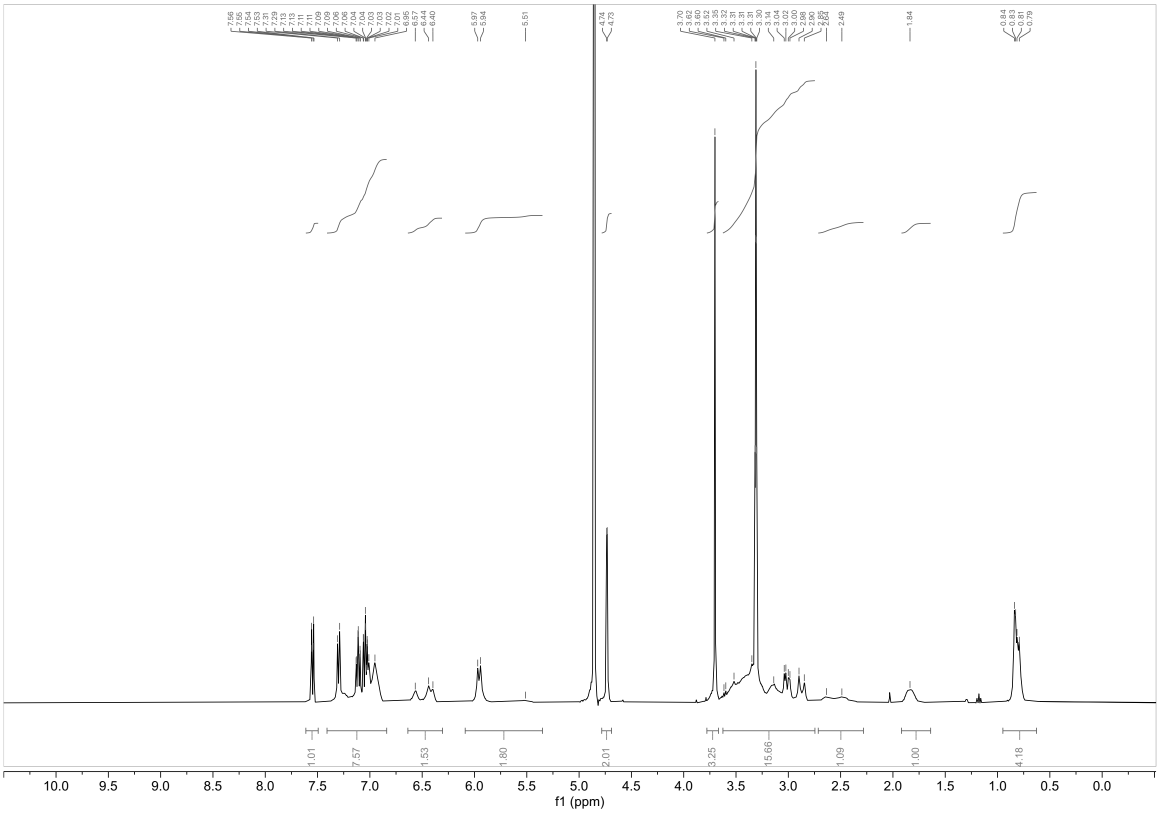

**$^{1}H$ NMR spectrum of WX-02-515 ($CD_3OD$, 400 MHz)**

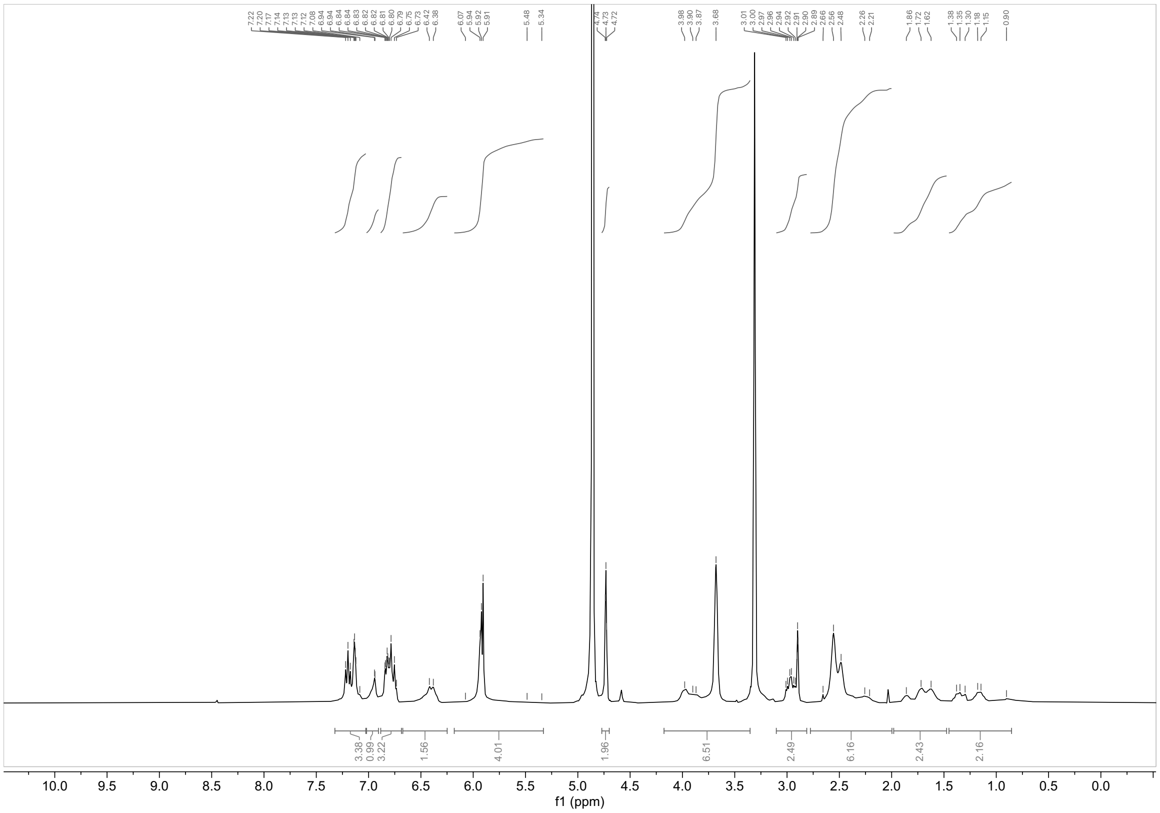

**$^{1}H$ NMR spectrum of WX-02-516 ($CD_3OD$, 400 MHz)**

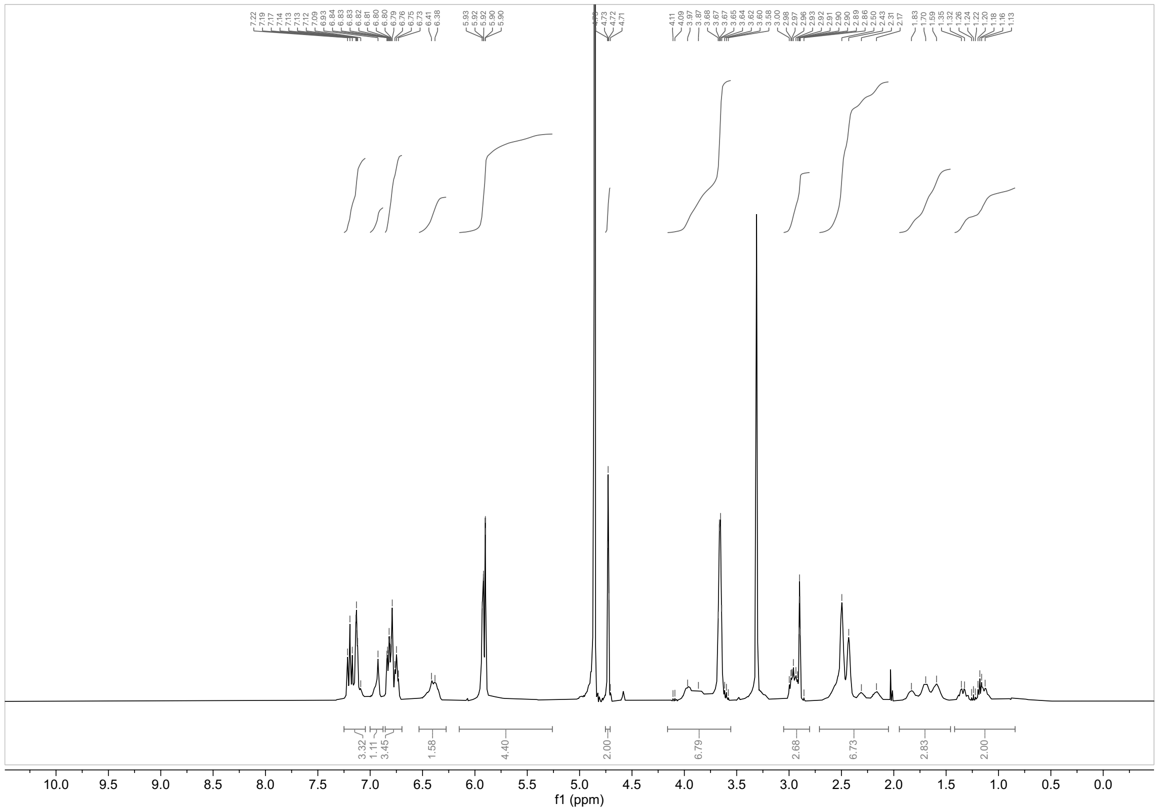


**$^1H$ NMR spectrum of WX-02-517 ($CD_3OD$, 400 MHz)**

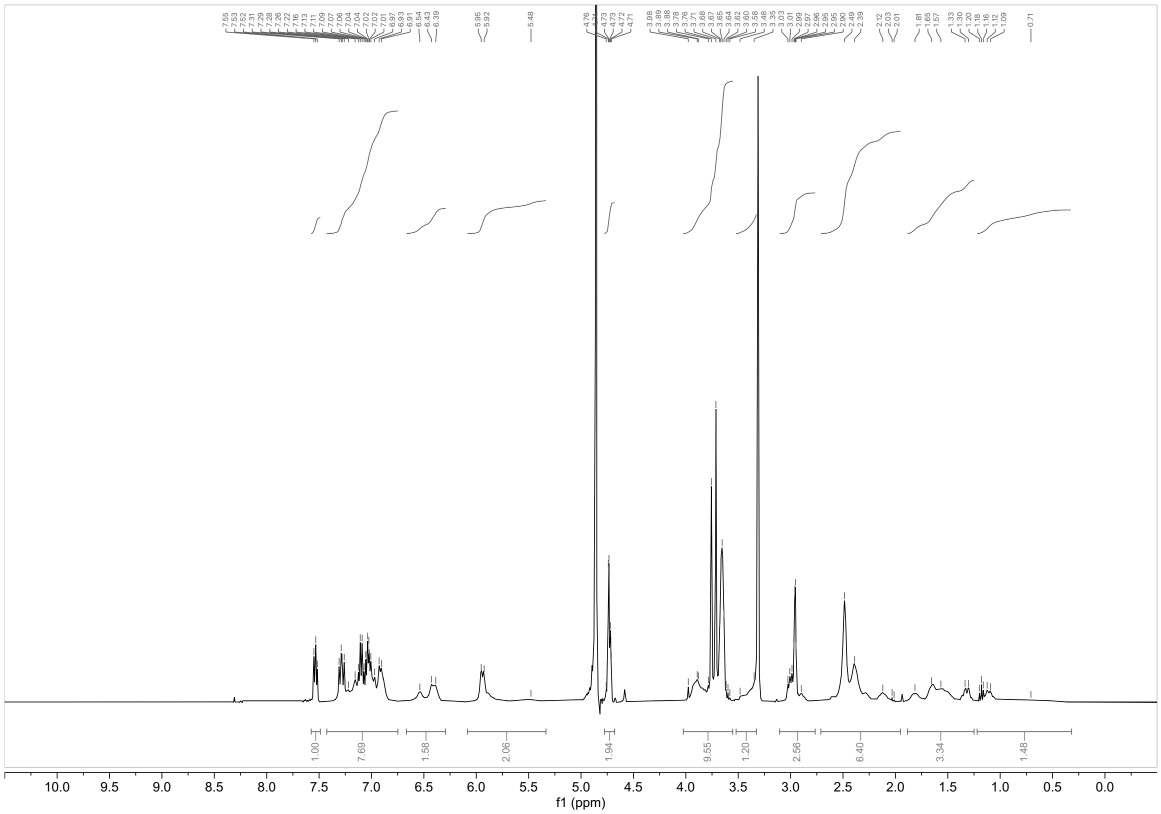


**$^1H$ NMR spectrum of WX-02-518 ($CD_3OD$, 400 MHz)**

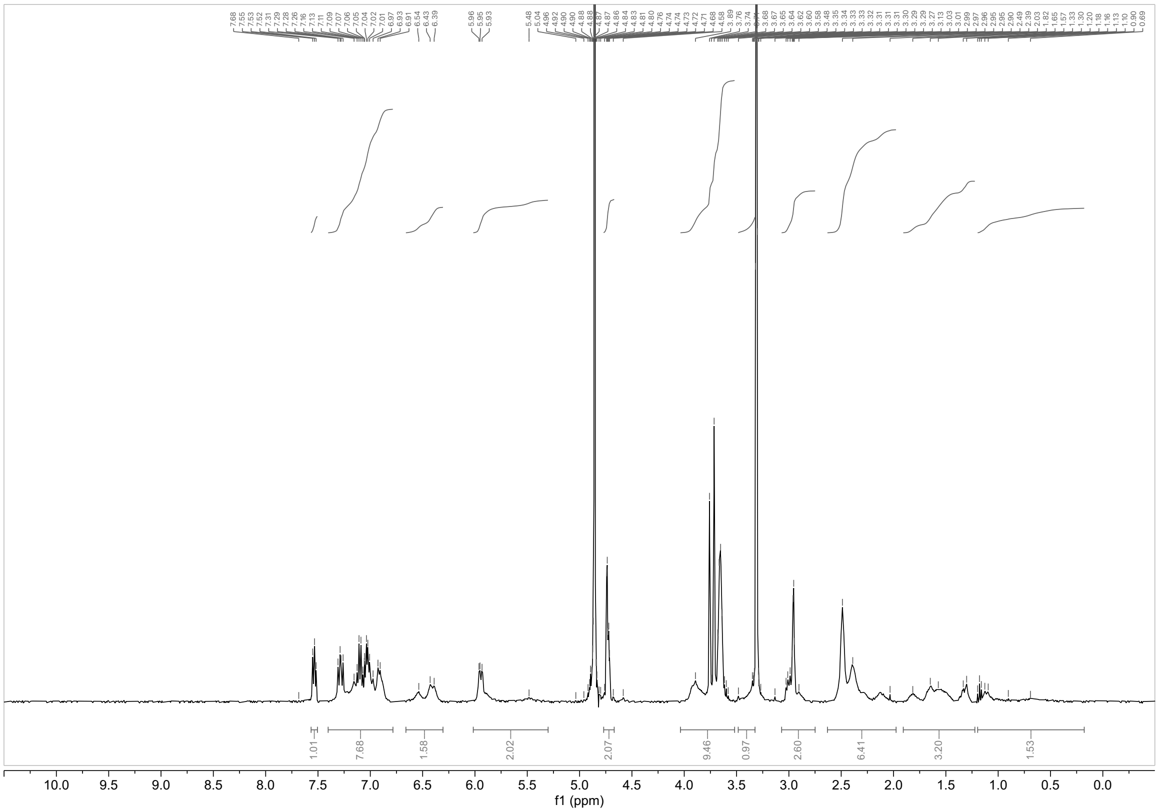

**$^{1}H$ NMR spectrum of WX-02-519 ($CD_3OD$, 400 MHz)**

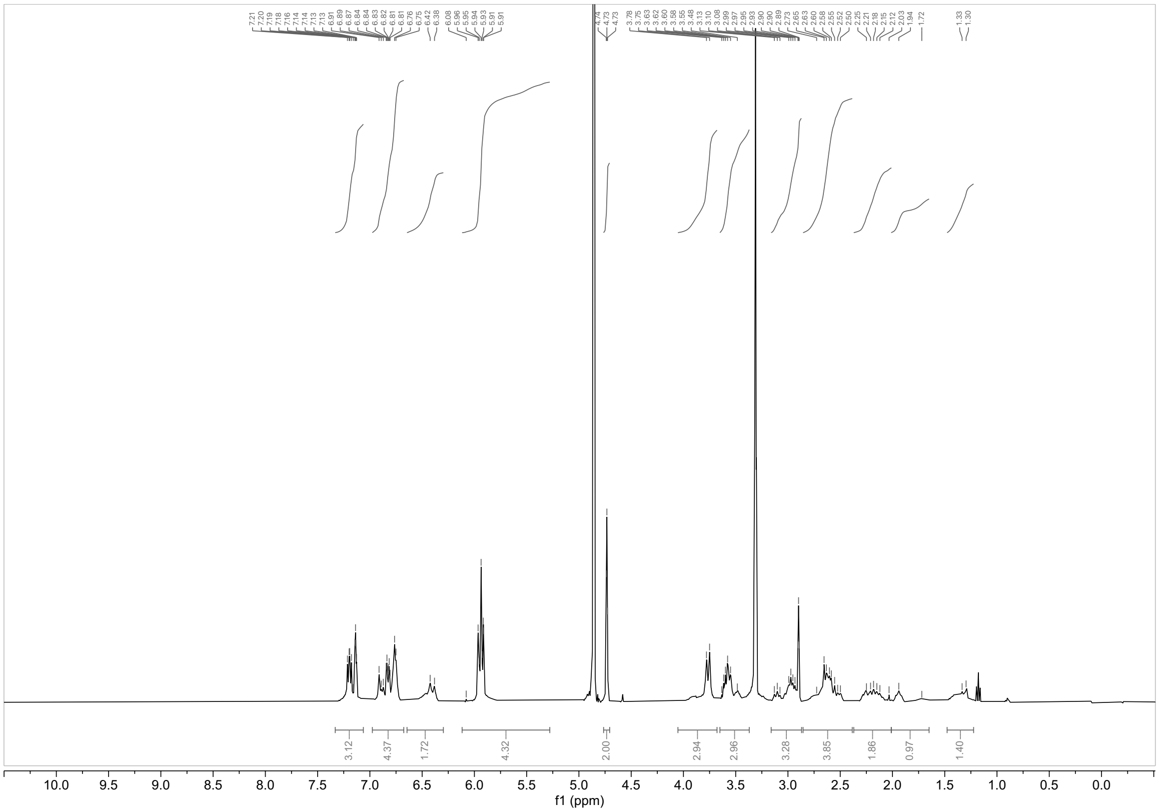

**$^{1}H$ NMR spectrum of WX-02-520 ($CD_3OD$, 400 MHz)**

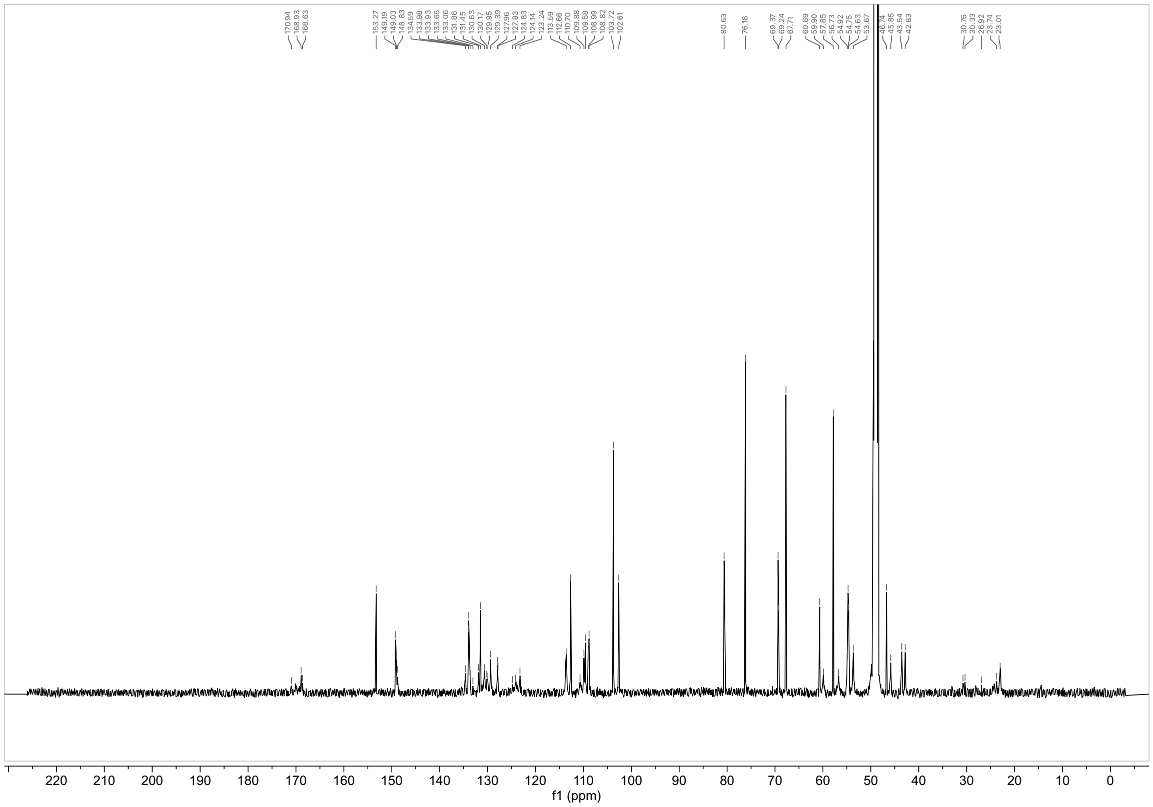

**$^{13}C$ NMR spectrum of WX-02-520 ($CD_3OD$, 150 MHz)**

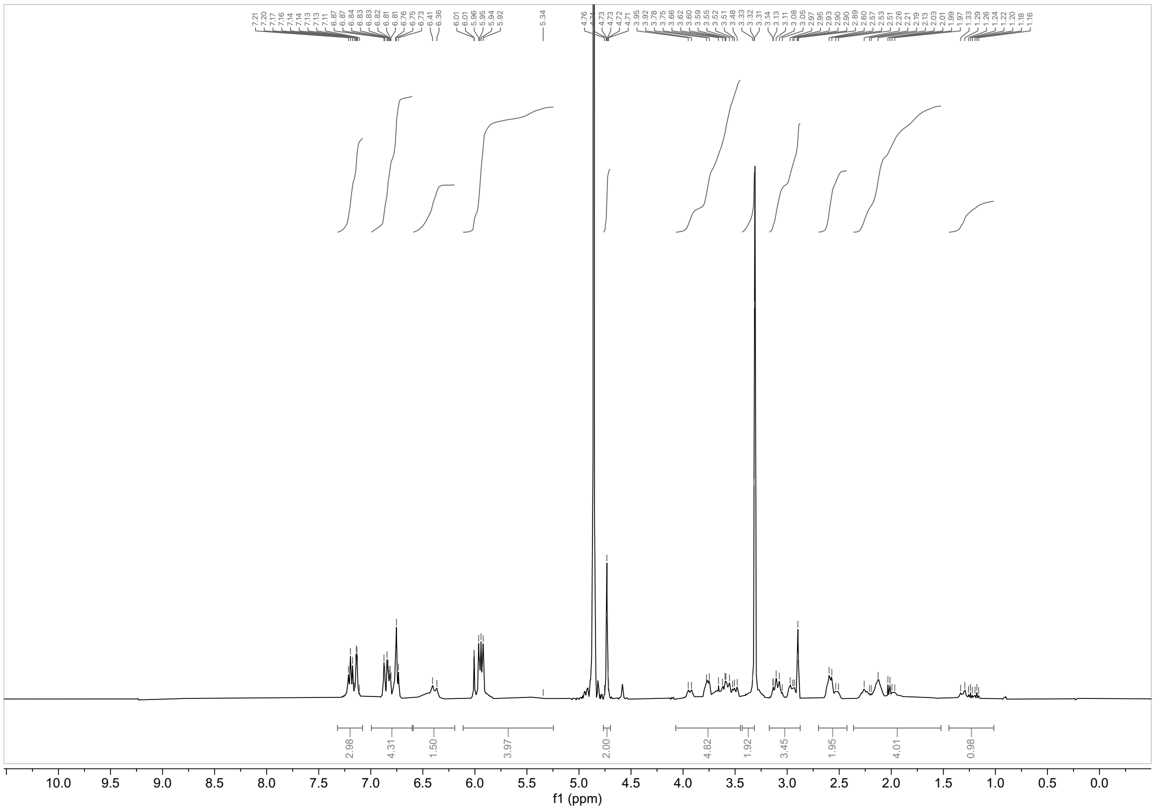

**$^{1}H$ NMR spectrum of WX-02-521 ($CD_3OD$, 400 MHz)**

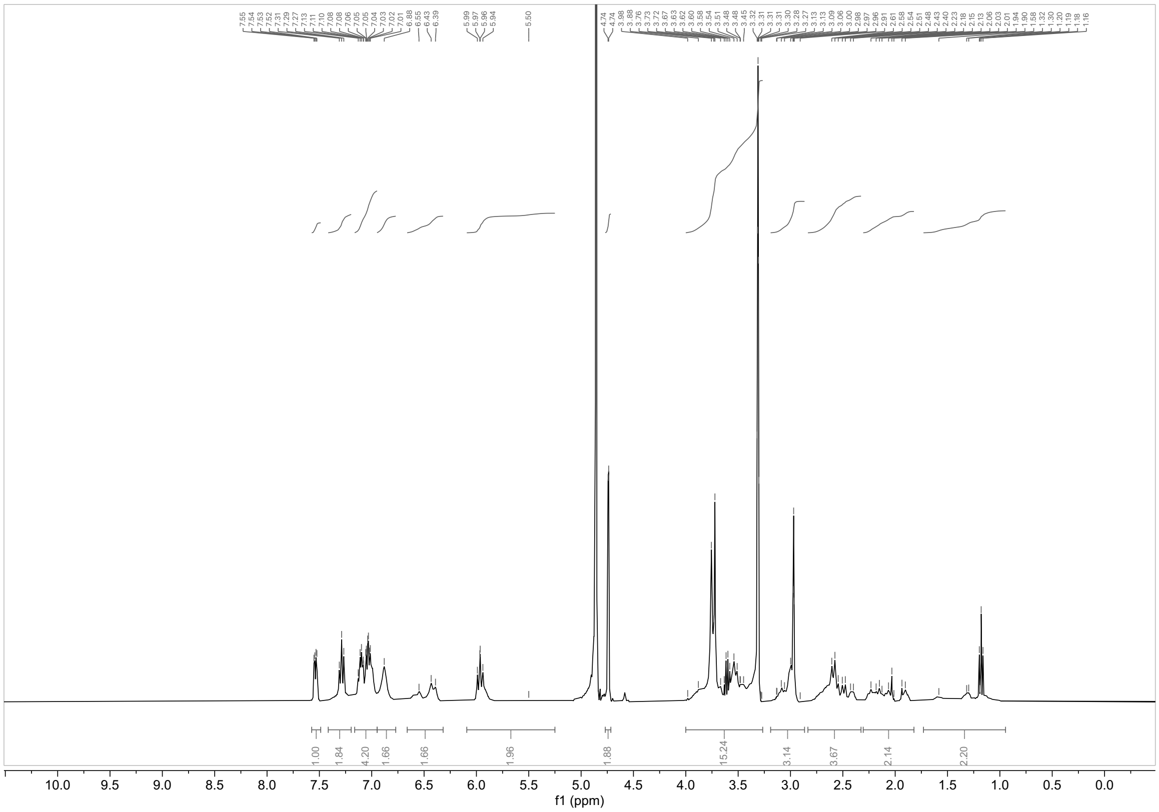


**$^{1}H$ NMR spectrum of WX-02-522 ($CD_3OD$, 400 MHz)**

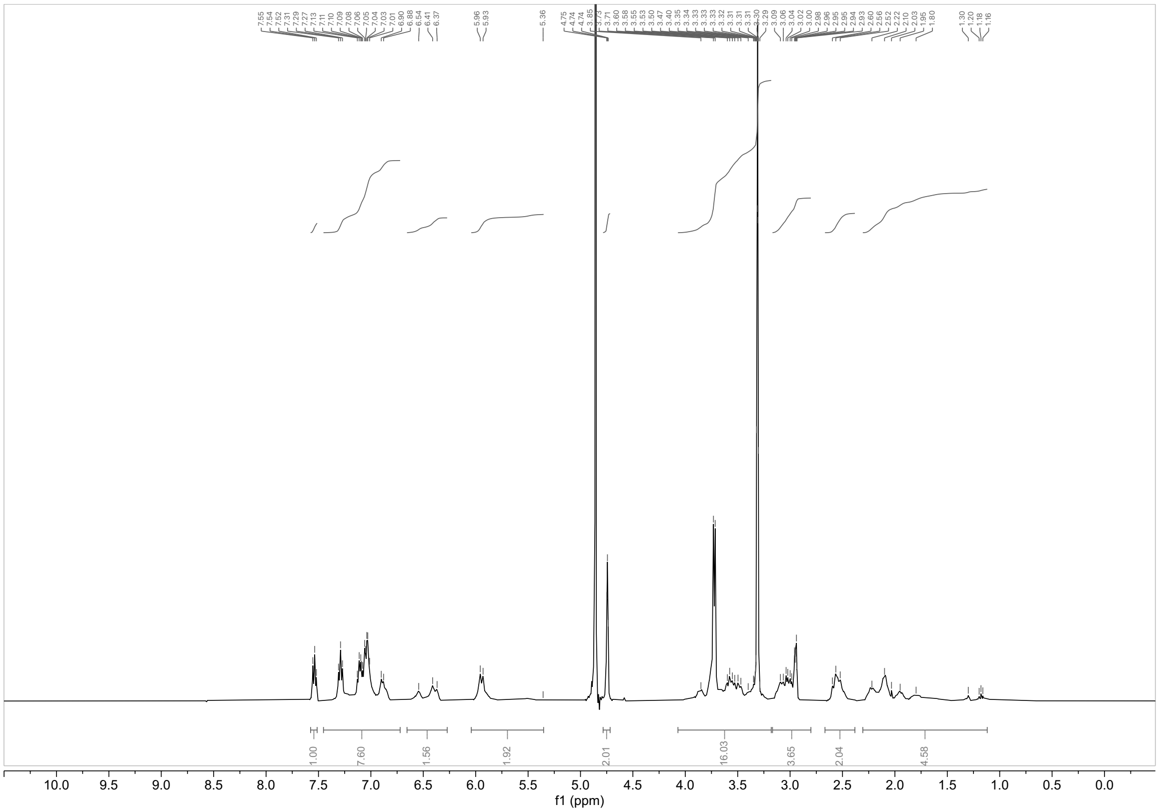


**$^{1}H$ NMR spectrum of WX-02-523 ($CD_3OD$, 400 MHz)**

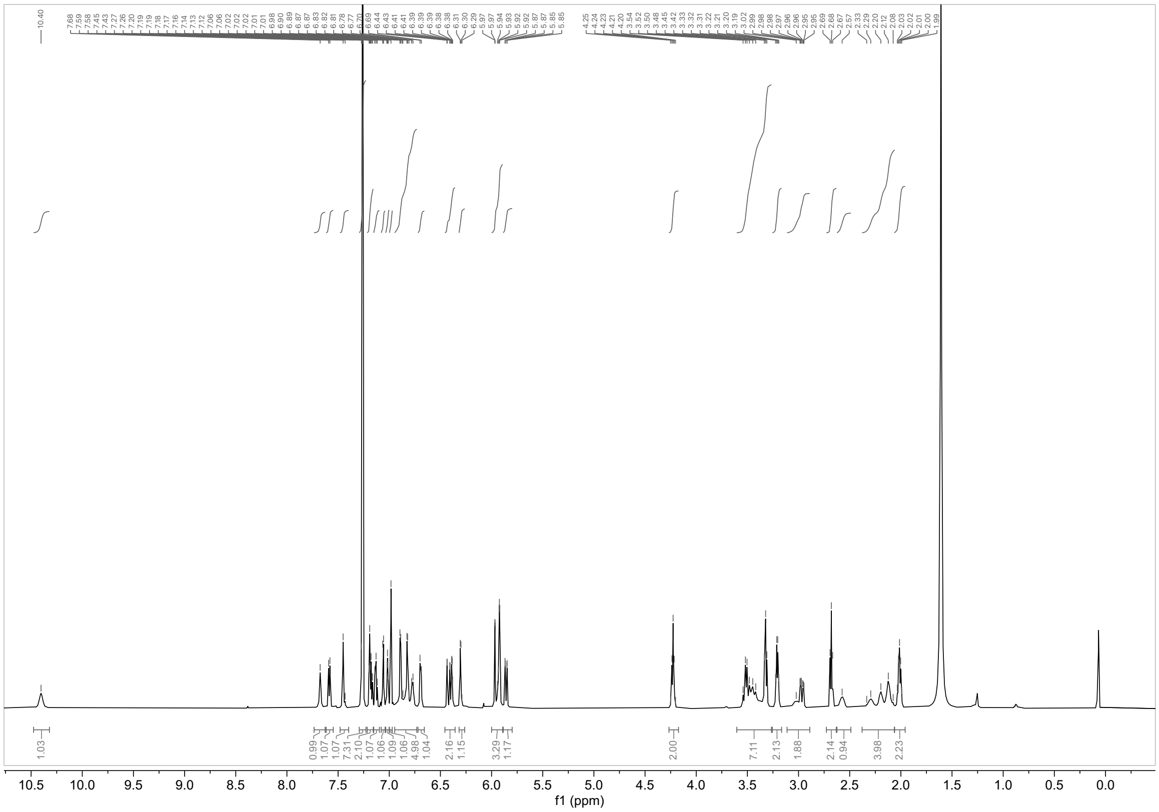

**$^1$H NMR spectrum of YZ-01-A ($CDCl_3$, 600 MHz)**

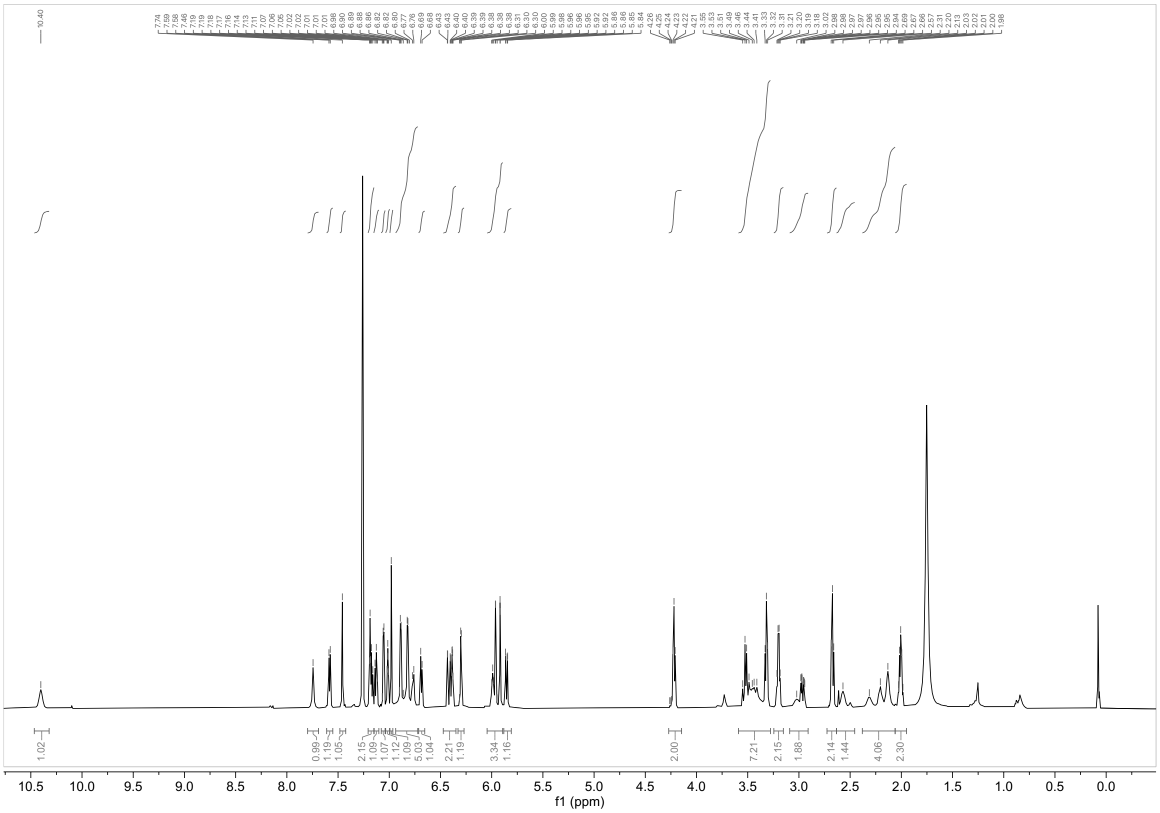

**$^1$H NMR spectrum of YZ-01-B ($CDCl_3$, 600 MHz)**

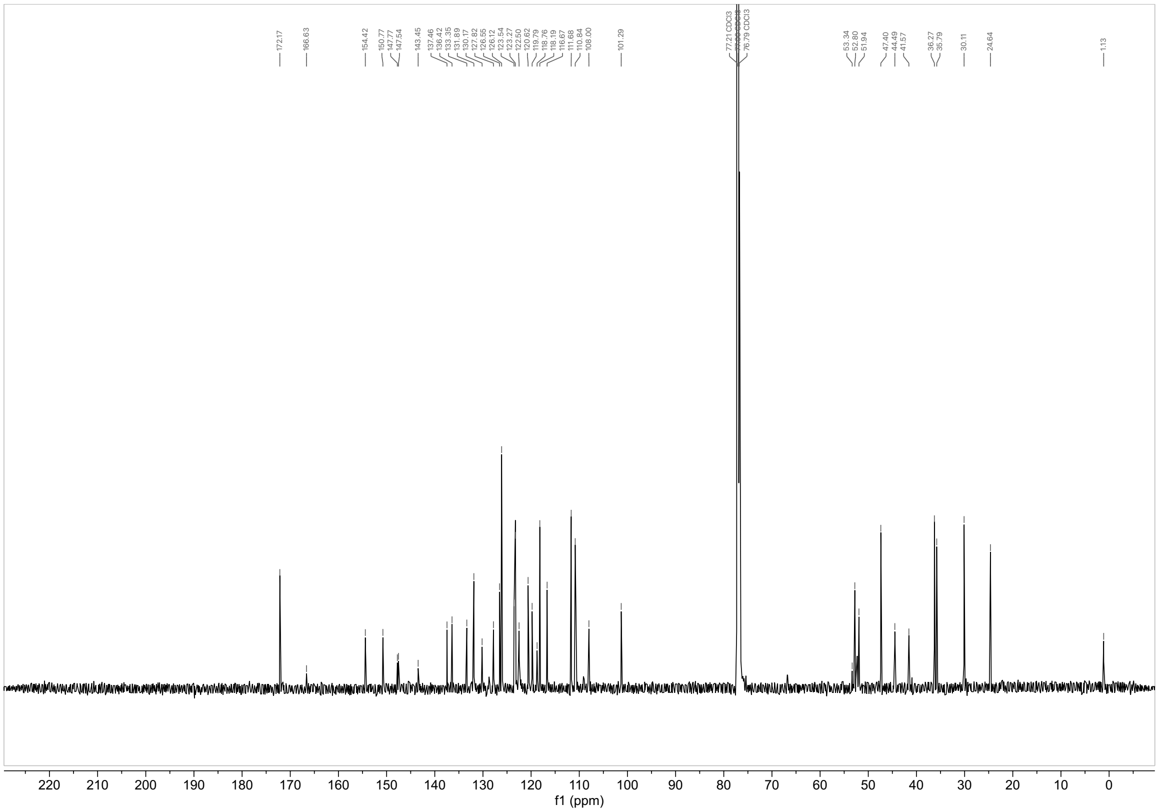

$^{13}C$ NMR spectrum of YZ-01-B ($CDCl_3$, 150 MHz)
