## Supplementary Table 1 for "Expanding the ligandable proteome by paralog hopping with covalent probes"

**Supplementary Table 1.** Crystallographic data collection and refinements statistics.

| | CDK2-cyclin E1 + I-125 | CDK2-cyclin E1 + I-198 |
|---|---|---|
| **Wavelength** | 1.000 | 1.000 |
| **Resolution range** | 84.280 - 1842 (1.960 - 1.842) | 84.271 - 1.896 (2.055 - 1.896) |
| **Space group** | P 41 21 2 | P 41 21 2 |
| **Unit cell** | 101.388 101.388 151.613 90 90 90 | 101.374 101.374 151.607 90 90 90 |
| **Total reflections** | 1869038 (178763) | 803761 (81845) |
| **Unique reflections** | 68650 (1043) | 63226 (329) |
| **Multiplicity** | 27.2 (26.6) | 12.7 (13.2) |
| **Completeness (%)** | 86.47 (25.7) | 79.50 (18.8) |
| **Mean I/sigma(I)** | 24.61 (1.6) | 23.5 (1.65) |
| **Wilson B-factor** | 38.95 | 43.15 |
| **R-merge** | 0.091 (4.49) | 0.071 (4.64) |
| **R-meas** | 0.093 (4.575) | 0.074 (4.82) |
| **R-pim** | 0.016 (0.467) | 0.017 (0.496) |
| **CC1/2** | 0.999 (0.649) | 1 (0.653) |
| **Reflections used in refinement** | 59380 (1043) | 50228 (329) |
| **Reflections used for R-free** | 3015 (49) | 2534 (14) |
| **R-work** | 0.199 (0.356) | 0.191 (0.301) |
| **R-free** | 0.217 (0.422) | 0.219 (0.490) |
| **CC(work)** | 0.958 (0.740) | 0.960 (0.780) |
| **CC(free)** | 0.958 (0.608) | 0.937 (0.301) |
| **Number of non-hydrogen atoms** | 4881 | 4790 |
| **macromolecules** | 4527 | 4516 |
| **ligands** | 44 | 48 |
| **solvent** | 310 | 226 |
| **Protein residues** | 568 | 565 |
| **RMS(bonds)** | 0.011 | 0.014 |
| **RMS(angles)** | 1.34 | 1.41 |
| **Ramachandran favored (%)** | 98.92 | 98.92 |
| **Ramachandran allowed (%)** | 0.90 | 1.08 |
| **Ramachandran outliers (%)** | 0.18 | 0 |
| **Rotamer outliers (%)** | 0.83 | 0.21 |
| **Clashscore** | 5.28 | 4.07 |
| **Average B-factor** | 46.20 | 49.14 |
| **macromolecules** | 45.71 | 48.92 |
| **ligands** | 45.41 | 48.28 |
| **solvent** | 53.49 | 53.61 |

Statistics for the highest-resolution shell are shown in parentheses.
Rmerge = ΣhklΣi|Ii − ⟨I⟩|/−ΣhklΣIi where Ii is the intensity of the ith observation, ⟨I⟩ is the mean intensity of the reflection, and the summations extend over all unique reflections (hkl) and all equivalents (i), respectively.
Rpim is a measure of the quality of the data after averaging the multiple measurements, and Rpim = Σhkl[n/(n − 1)]1/2Σi|Ii(hkl) − ⟨I(hkl)⟩|/ΣhklΣiIi(hkl) where n is the multiplicity and other variables are as defined for Rmerge ([70]).
CC1/2 is the Pearson correlation coefficient.
Rwork = Σ|Fo − Fc|/ΣFo where Fo and Fc are observed and calculated structure factors, respectively, Rfree was calculated from a randomly chosen 5% of reflections excluded from the refinement, and Rwork was calculated from the remaining 95% of reflections.

**Commented [BC1]:** This table does not need to be in the main manuscript file. it can be supplied as a separate Supp Table file.

Please move to a separate word doc named Supplementary Table 1
